# Gut-Initiated Alpha Synuclein Fibrils Drive Parkinson’s Disease Phenotypes: Temporal Mapping of non-Motor Symptoms and REM Sleep Behavior Disorder

**DOI:** 10.1101/2024.04.22.590542

**Authors:** Daniel Dautan, Wojciech Paslawski, Sergio G. Montejo, Daniel C. Doyon, Roberta Marangiu, Michael G. Kaplitt, Rong Chen, Valina L. Dawson, Xiaoaun Zhang, Ted M. Dawson, Per Svenningsson

## Abstract

Parkinson’s disease (PD) is characterized by progressive motor as well as less recognized non-motor symptoms that arise often years before motor manifestation, including sleep and gastrointestinal disturbances. Despite the heavy burden on the patient’s quality of life, these non-motor manifestations are poorly understood. To elucidate the temporal dynamics of the disease, we employed a mouse model involving injection of alpha-synuclein (αSyn) pre-formed fibrils (PFF) in the duodenum and antrum as a gut-brain model of Parkinsonism. Using anatomical mapping of αSyn-PFF propagation and behavioral and physiological characterizations, we unveil a correlation between post-injection time the temporal dynamics of αSyn propagation and non-motor/motor manifestations of the disease. We highlight the concurrent presence of αSyn aggregates in key brain regions, expressing acetylcholine or dopamine, involved in sleep duration, wakefulness, and particularly REM-associated atonia corresponding to REM behavioral disorder-like symptoms. This study presents a novel and in-depth exploration into the multifaceted nature of PD, unraveling the complex connections between α-synucleinopathies, gut-brain connectivity, and the emergence of non-motor phenotypes.

**One Sentence Summary:** Propagation of αSyn from the gut-to-the-brain induces RBD and several non-motor and motor phenotypes of Parkinson’s disease.

## INTRODUCTION

Non-motor symptoms in Parkinson’s disease (PD) encompass a diverse array of manifestations that can occur years before motor symptoms and impact very early on patients’ life (*1*, *2*). Rapid eye movement (REM)-sleep behavior disorder (RBD) is the prodromal manifestation with the highest likelihood ratio to phenoconvert to PD (*3*).

The pathological hallmark of PD is the accumulation of misfolded and aggregated alpha-synuclein (αSyn) that participates in Lewy Bodies formation (*4*, *5*). Aggregated forms of αSyn have increased phosphorylation of the serine 129 (pSer129) residue, although pSer129 phosphorylation may also regulate physiological synapse function (*6*). Most recent hypotheses suggest an origin of PD spreading throughout the central nervous system along the gut-brain axis and the vagus nerve (*7*, *8*). This spreading of pathologic αSyn through interconnected brain regions contributes to the heterogeneity of PD’s clinical representation, including the emergence of non-motor symptoms (*9*).

Most conventional models focus almost exclusively on the late stages of PD. This limitation has been a significant challenge for researchers and poses hurdles for understanding the disease. While not without limitations, the injection of αSyn pre-formed fibrils (PFF) in the stomach/duodenum represents a step forward in this direction, providing a clinically relevant model offering insight into the early- and late stages of PD (*10–13*).

To address these questions, we used behavioral and physiological characterization of clinical motor and non-motor representation of the disease in mice injected with αSyn-PFF at different time points. This approach demonstrated the role of dopaminergic structures, as well as their progressive degeneration in the appearance of non-motor symptoms, particularly impaired sleep architecture and RBD-like dysfunction.

## RESULTS

### Injection of αSyn-PFF induced the formation of αSyn aggregates along the digestive tract

Sonication of mature αSyn fibrils facilitates the generation of αSyn-PFF (*12*), with the majority exhibiting a length of less than 50 nm (**Fig. 1A**). Injection of αSyn-PFF into the muscularis layers of the duodenum and glandular stomach (**Fig. 1B**) enables rapid diffusion and aggregation of αSyn (pSer129 staining) throughout the entire stomach, including the forestomach (**Fig. 1C**). To assess the temporal diffusion in the periphery and central nervous system, 71 animals were injected with either monomeric αSyn (2 weeks or 7 months) or αSyn-PFF at different time points (2 weeks, 1 month, 3 months, 6 or 7 months). All animals were born around the same time and tested together to minimize group variability (**Fig. 1D**). Following behavioral assessments, animals were sacrificed, and tissues were collected. Staining for pSer129 revealed a trend towards an increase of pSer129 large clusters (>200px) in the upper gastrointestinal (Gi, i.e. Duodenum/Jejunum) tract (**Fig. 1E-H**) as well as an early increase in large-size aggregated clusters (∼3 months) in the colon (**Fig. 1I-L**) which is consistent with observation in prodromal PD patients where αSyn accumulation in the enteric nervous system precedes the progression of Lewy pathology (*14*). In the colon, the increase of large clusters was also paired with increase of the number of clusters independently of their sizes at 6 months suggesting an accumulation over time in all layers of the tissue (**Fig. 1I-L**).

**Fig. 1.**
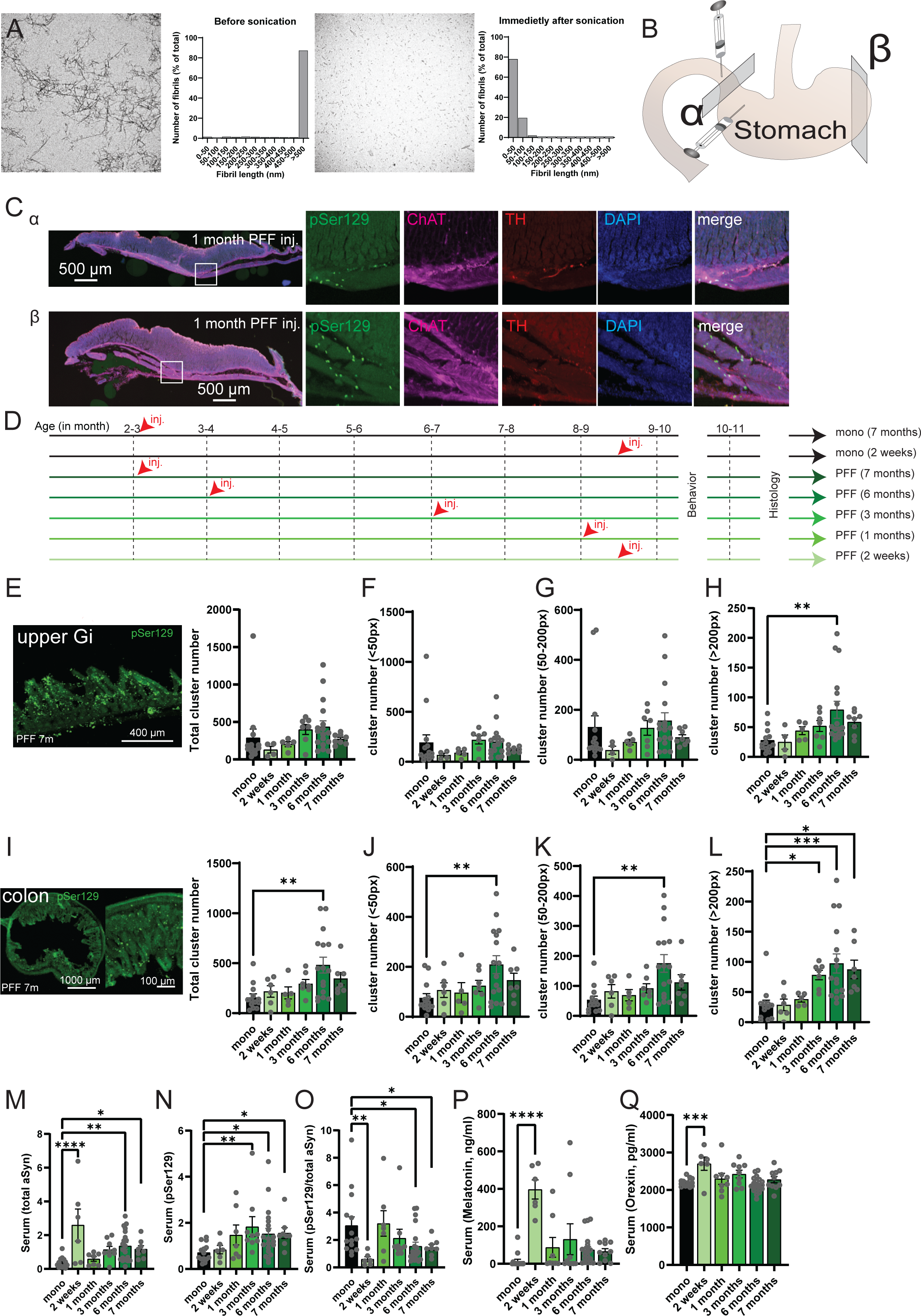
Propagation of alpha synuclein aggregates in the periphery following injections of preformed fibrils in the stomach. (**A**) Transition electron microscopy of fibrils before and after sonication revealed a change from long length fibrils to a majority of small sized fibrils. (**B**) Graphical representation of the injection sites in the muscularis layers of the duodenum and the stomach in mice. α represents transversal section at the level of the injection site (**C**) while β illustrates sections located on the distant side (**C**). (**C**) Confocal images of transversal sections of the stomach 1 month following injections at 2 different levels (α and β) showing staining of pSer129, ChAT, TH and DAPI. (**D**) Experimental design. Age-matched animals were born around the same period of time were tested for behavior and electrophysiology at the same time. Each group, including monomeric injected or PFF injected animals were receiving gut-injection at different time. (**E**) Confocal images of pSer129 in the upper gastrointestinal tract (intestine) and total number of pSer129 clusters in the upper Gastrointestinal (i.e. intestine) of mice injected with monomeric αSyn or αSyn-PFF in the stomach (F_5,47_=3.257, p=0.0132). (**F**) Total number of pSer129 small clusters (<50px) in the upper Gastrointestinal of mice injected with monomeric αSyn or αSyn-PFF in the stomach (F_5,47_=3.230, P=0.0138). (**G**) Total number of pSer129 medium sized clusters (50-200px) in the upper Gastrointestinal of mice injected with monomeric αSyn or αSyn-PFF in the stomach (F_5,47_=2.346, P=0.0482). (**H**) Total number of pSer129 large sized clusters (>200px) in the upper Gastrointestinal of mice injected with monomeric αSyn or αSyn-PFF in the stomach (F_5,48_=3.306, P=0.0121). (**I**) Confocal images of pSer129 in the colon and total number of pSer129 clusters in the upper colon of mice injected with monomeric αSyn or αSyn-PFF in the stomach (F_5,48_=4.059, P=0.0037). (**J**) Total number of pSer129 small clusters (<50px) in the colon of mice injected with monomeric αSyn or αSyn-PFF in the stomach (F_5,47_=3.348, P=0.0114). (**K**) Total number of pSer129 medium sized clusters (50-200px) in the colon of mice injected with monomeric αSyn or αSyn-PFF in the stomach (F_5,47_=4.632, P=0.0016). (**L**) Total number of pSer129 large sized clusters (>200px) in the COLON of mice injected with monomeric αSyn or αSyn-PFF in the stomach (F_5,47_=6.707, P<0.0001). (**M-O**) Quantification of total αSyn (**M**, F_5,58_=5.806, P=0.0002), pSer129 (**N**, F_5,58_=2.448, P=0.0433) or the ratio pSer129/total αSyn (**O**, F_5,58_=2.880, P=0.0217) in the serum of mice injected with monomeric αSyn or αSyn-PFF, post hoc Fisher LSD. (**P**) Concentration of melatonin in the serum of mice injected with monomeric αSyn or αSyn-PFF in the stomach (F_5,62_=9.481, P<0.0001), post hoc Fisher LSD. (**Q**) Concentration of orexin in the serum of mice injected with monomeric αSyn or αSyn-PFF in the stomach (F_5,63_=5.078, P=0.0006), post hoc Fisher LSD. Data are expressed as mean±SEM. All individual points represent individual animal. * P<0.05, ** P<0.01, *** P<0.001, **** P<0.0001. Statistical data represent post hoc analyses compared to monomeric αSyn group following 1-way ANOVA.

Furthermore, plasma from intra-cardiac blood was analyzed using western blot, demonstrating an increase in total αSyn (**Fig. 1M**), pSer129 (**Fig. 1N**), as well as a reduction in the ratio of pSer129/total αSyn (**Fig. 1O**). Similarly, a rapid increase in serum concentrations of Melatonin (**Fig. 1P**) and Orexin (**Fig. 1Q**), suggesting possible sleep disturbances, was observed immediately following αSyn-PFF injection (∼2 weeks) persisting, but to a lower extent, for the duration of the experiments. Collectively, our data suggest that αSyn-PFF, after injection into the stomach/duodenum, propagates retrogradely and anterogradely along the vagus nerve, affecting various tissues and may contribute to non-motor phenotypes in PD patients.

### Injection of αSyn-PFF is sufficient to alter microbiota with a time-sensitive pattern

Analyses (16S – OPU) of the freshly collected feces RNA_microbiota show a strong reduction of alpha diversity following a specific time distribution (**Fig. 2A-B, S1**) as well as beta diversity (**Fig. 2C, S2**) as early as 1 month following the injection of the αSyn-PFF. Specific analyses of phylum suggest an early reduction of cyanobacteria (∼2 w) as well as decrease of bacteroidota and increase of firmicutes (∼1 m) following injection (**Fig. 2D, S3**). Similarly, we observed a reduction of families including Bacteroidaceae and Rikenellacea at an early timepoint following injection of αSyn-PFF (∼2 w, **Fig. S4**), while Lactobacillaceae and Lachnospiraceae families increased following injection and remain high for the remaining of the experiments (**Fig. S4**). We also observed an increase of the Lactobacillus genus starting 1 m after injection (**Fig. S5**), and a decrease of the Bacteroides_acidifaciens species (∼2 w, **Fig. 2E, S6**) and the order Bacteroidales (**Fig. 2F, S7**), while the class Vampirivibronia and Bacteroida decrease strongly and remain low following injection of αSyn-PFF (**Fig. 2G, S8**). Interestingly, the linear discriminant analyses effect size (LFEfSE) (**Fig. 2H, S9**) and the cladogram (**Fig. 2I**) compared to monomeric αSyn injected mice revealed several clusters like the one observed in inflammation as well as in PD cohorts.

**Fig. 2.**
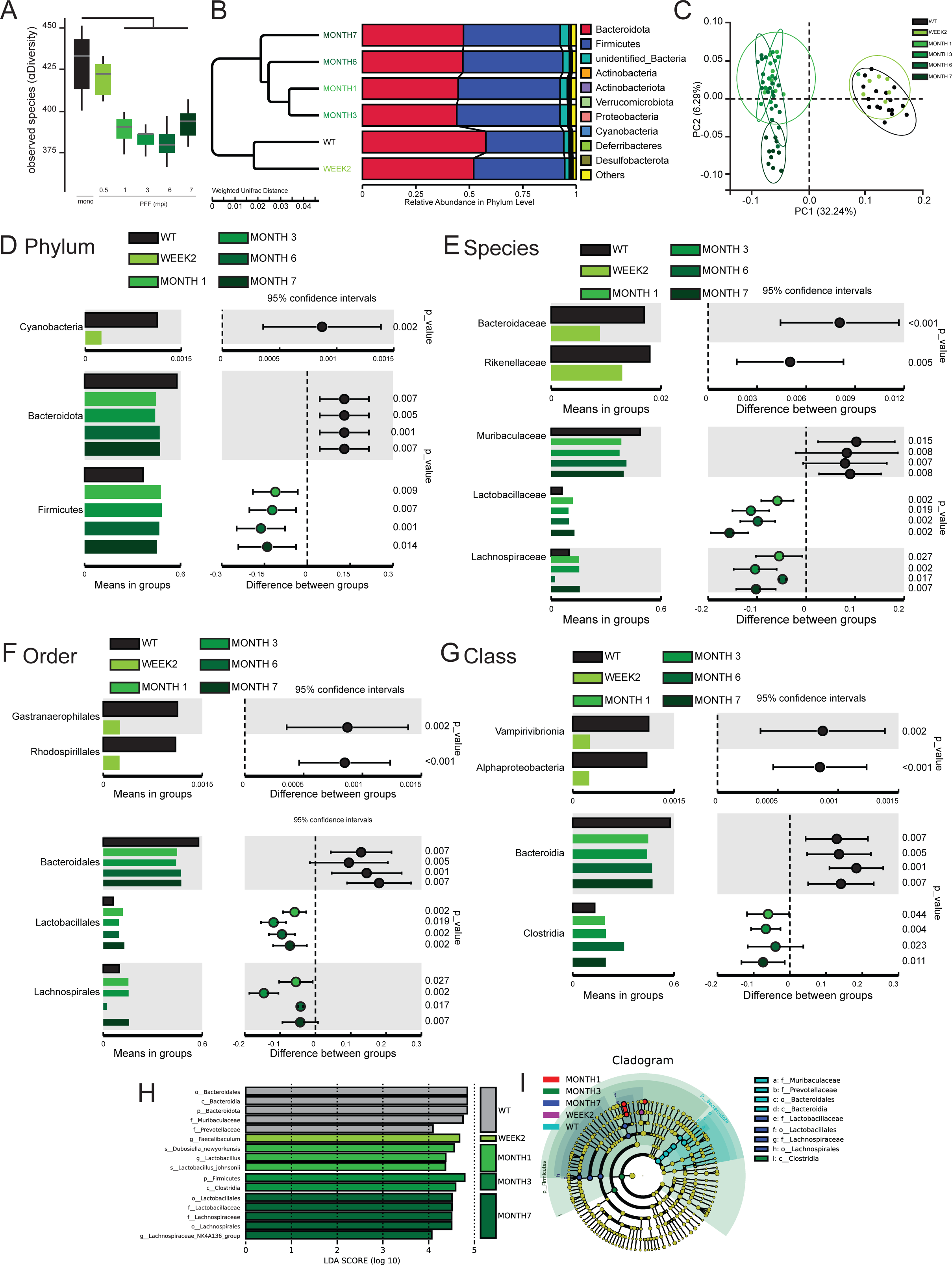
Time dependent alteration of microbiota following injections of preformed fibrils in the stomach. (**A**) Observed alpha diversity in the microbiota of mice injected with monomeric αSyn or αSyn-PFF. (**B**) Cluster-based representation of the relative abundance at the Phylum levels of mice injected with monomeric αSyn or αSyn-PFF. (**C**) Principal component analyses of the Beta diversity of mice injected with monomeric αSyn or αSyn-PFF. (**D-G**) Relative abundance of specific Phylum (**D**), Species (**E**), Order (**F**) or Class (**G**) with comparison between groups of mice injected with monomeric αSyn or αSyn-PFF. (**H**) LDA score of specific order in mice injected with monomeric αSyn or αSyn-PFF. (**I**) Cladogram of specific order in mice injected with monomeric αSyn or αSyn-PFF. Data are expressed as mean ± SEM. Box plots are representing the median, lower and upper quartile as well as the minimum and maximum value. All individual point represents individual animals Statistical P-value are given following post hoc analyses compared to monomeric αSyn group following 1-way ANOVA, n=71 values.

Altogether, we found that αSyn-PFF injection in the stomach impacts gut microbiota in mice, with similar modulation of bacteria diversity and sub-type to the one observed in PD patients (*15–17*).

### Time-dependent manner propagation of pathologic alpha-synuclein in the central nervous system correlates with sleep manifestations and other non-motor phenotypes

According to the Braak model of gut-brain axis αSyn propagation, the formation of pathologic alpha-synuclein in the gut will then propagate to the central nervous system along the vagus nerve and induce non-motor phenotypes based on the structures impacted at each stage (*7*, *18*). To see if such a time-dependent manner between the propagation in the brain and non-motor phenotypes can be observed using the αSyn-PFF model, we first whole-brain mapped pSer129 staining in the brain of all the animals using DAB-staining (**Fig. 3, S10**) (*19*, *20*). After processing the entire brain slices, and scanning at high resolution, we cropped and aligned the different sections before proceeding to a normalized density plot and statistical analyses on 30 µm^2^ ROIs (**Fig. S10)** (*21*). This allowed us to confirm that immediately following injection of αSyn-PFF in the gut, we observed increased level expression of pSer129 in the dorsal vagus nerve (DMV; ∼2w to 1m, **Fig. 3H**) followed by the gigantocellular nuclei (Gi, ∼1 m, **Fig. 3H**), tractus solitarus (NTS, ∼1m, **Fig. 3H**), the pons (∼3 m, **Fig. 3G**) and the Globus Pallidus (GPe)/Striatum (∼6 m, **Fig. 3B**). Interestingly, increased staining in the Substantia nigra (SN, **Fig. 3E**) was observed early (∼3 m), similar to cortical structure (∼3-6 m, **Fig. 3H, A-D**). Despite the significant increase of pSer129 in the thalamus (**Fig. 3D)**, we were not able to define the exact stage at which the effect was significant, most probably due to the complexity of thalamus anatomy (*22*). Similarly, an increase of pSer129 was found in vestibular structures (**Fig. 3G-H)**, including the trigeminal nuclei, but post hoc analyses suggest the effect to be significant only in late stages (∼6 m).

**Fig. 3.**
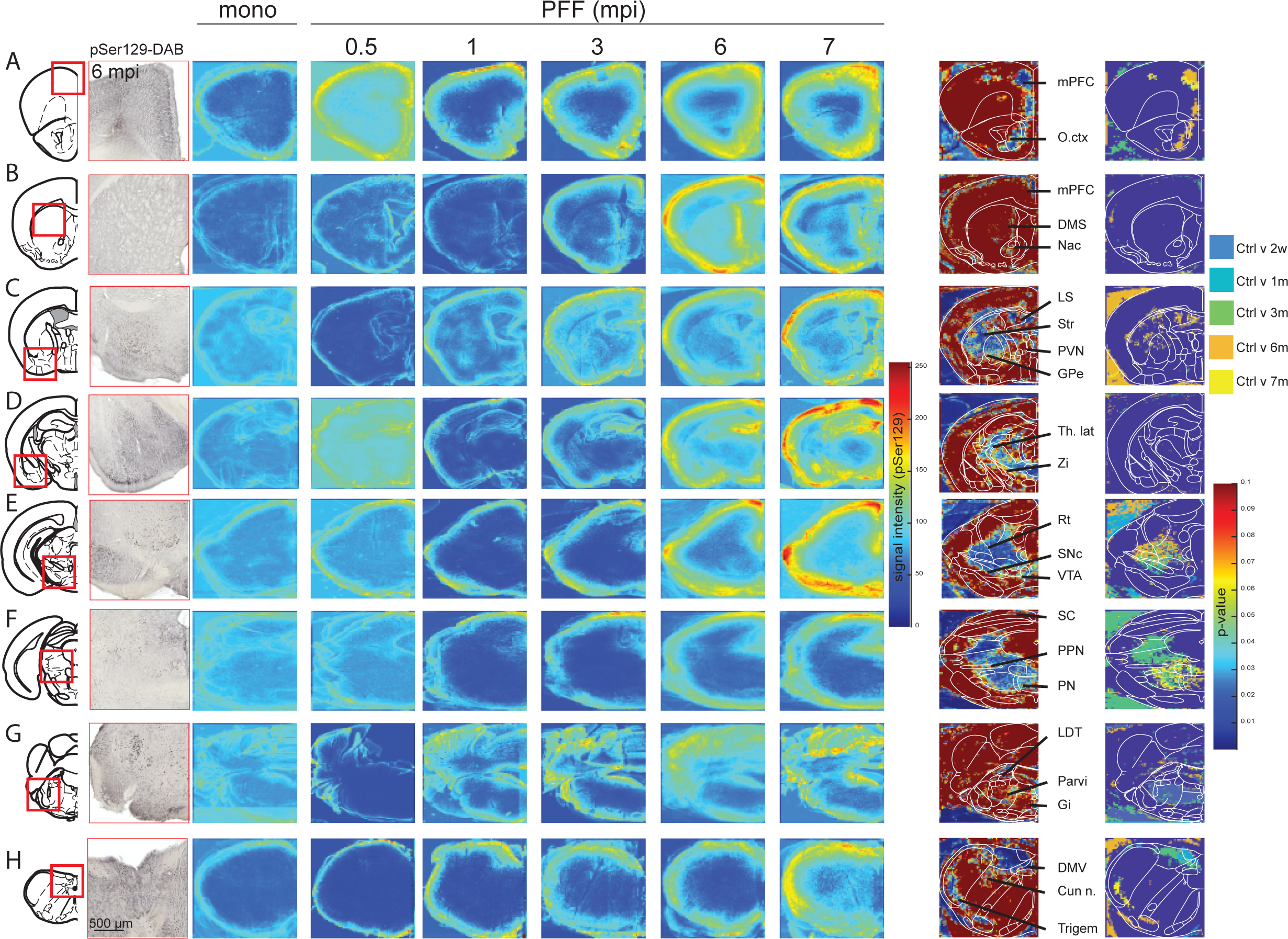
Time dependent propagation of pSer129 in the brain. Whole brain mapping of the temporal propagation of pSer129 DAB staining in (from top to bottom) frontal cortex (**A**), striatum (**B**), forebrain (**C**), thalamus (**D**), midbrain (**E**), brainstem (**F**), Pons (**G**) and medulla (**H**). Density-plot are estimated on a 5×5 pixels ROIs for each animal. P-value is obtained from “ROIs by ROIs” comparison between groups. Post hoc analyses are obtained using multiple comparison to the monomeric αSyn group, n=71 values. DAB images are obtained from the same animals injected with PFF in the stomach for 6 months, localization of the insert is represented with the red square on the brain outlines. Statistical data represent post hoc analyses compared to monomeric αSyn group following 1-way ANOVA.

To check the possible effect of pSer129 on neuronal viability, we performed immunofluorescent labeling using marker specific for cell types that have been described to be reduced post-mortem in PD (**Fig. S11**) including Tyrosine hydroxylase (**TH**) or acetylcholine transferase (**ChAT**). In the DMV (**Fig. S11A**) (*23*), staining for pSer129, 4’,6-diamidino-2-phenylindole (DAPI) and ChAT revealed a decrease of ChAT+ neurons as early as 1 month following injection (**Fig. S11B**) that was in strong anticipation with the reduction of DAPI (**Fig. S11C**) or the ratio ChAT^+^/DAPI^+^ (**Fig. S11D**) suggesting a loss of cholinergic markers in early stages rather than a neuronal degeneration. Similarly, in the midbrain (**Fig. S11E**) we found a reduction of TH+ neurons in the SNc at 3 months post injection (**Fig. S11F**) while the DAPI reduction and the trends towards a reduced ratio TH/DAPI ratio occurred at 6m (**Fig. S11G-H**), while the VTA present a decrease of TH+ neurons around 6 months post-injection (**Fig. S11I**) was observed with no significant reduction of DAPI or TH/DAPI ratio (**Fig. S11J-K**). This seems to confirm TH-markers reduction in early stages that preceded neurodegeneration.

Interestingly, we observed the presence of pSer129 in the brainstem including the Laterodorsal tegmental nucleus (LDTg) and the Pedunculopontine nucleus (PPN, **Fig. S11L**) and the ventral dorsal raphe (DR, **Fig. S11M**) (*24*, *25*). This Confirm previous observations in PD patients (*26*, *27*), where increased accumulation of alpha-synuclein in the LDTg and the DR correlated with sleep disorders and depressive-like phenotypes (*28*, *29*).

To characterize if specific phenotypes, typically observed early in PD patients, occurred following injection of αSyn-PFF in the stomach, we tested the cohort of mice for specific behaviors assessing non-motor symptoms.

One of the main non-motor symptoms, which appears in the prodromal and early stages of the disease and correlate with increased risk of PD, is shortened REM as well as RBD (*30*). To assess to what extent αSyn-PFF injection into the stomach could impair sleep, we implanted all the animals with a subcutaneous telemetry device collecting frontal electroencephalogram (EEG) and neck-muscles electromyogram (EMG). Automated sleep scoring allows us to define the different stages of sleep, including awake, REM and non-REM (NREM). Using significant (>2SD) muscle contraction in the root mean square of the EMG signal during REM events (Paradoxical sleep), we determined the number of events without atonia defined as RBD (**Fig. 4A)**. We found that ∼3 months following injection the number of RBD there was a trend toward an increase in the number of RBD that reached a significant effect at 6 months (**Fig. 4B**), while the percentage of REM events with RBD increased significantly starting at 3 months (**Fig. 4C**). Interestingly, when comparing cortical oscillation during RBD events in animals that display at least 1 RBD-event, we found that theta frequencies increase around the 3^rd^-month post injection and reach significance at 6 months (**Fig. S12B**) during RBD events without affecting other oscillations (**Fig. S12A, C-E**). Moreover, αSyn-PFF injection seems to modify cortical oscillation only during RBD, with no impact during awake event (**Fig. S12F**), REM (**Fig. S12G**), or NREM (**Fig. S12H**) despite an increased trend for delta oscillation during awake and NREM (*31*, *32*).

**Fig. 4.**
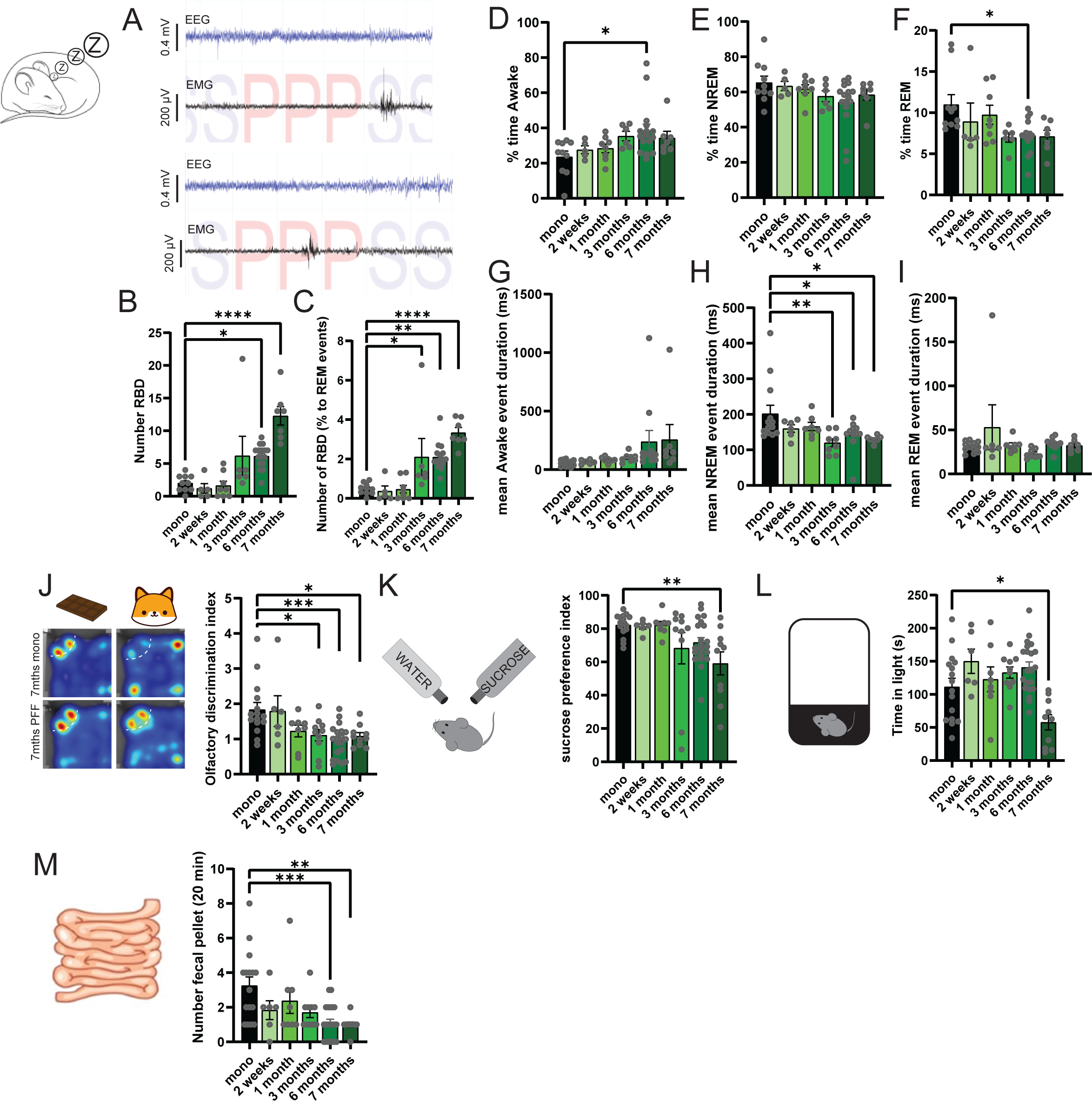
Time dependent development of non-motor phenotypes following injection of PFF in the stomach. (**A**) Example of EEG (blue) and EMG (back) signal during NREM (S) or REM (P) sleep. The top panel represents muscle contraction during NREM sleep, while the bottom panel represent REM-sleep behavior disorders (RBD). (**B**) Number of RBD events per 24h in mice injected with monomeric αSyn or αSyn-PFF (n=52 values, F_5,46_=13.40, P<0.0001). (**C**) Number of RBD events normalized by the individual number of REM events in mice injected with monomeric αSyn or αSyn-PFF (n=52 values, F_5,46_=12.87, P<0.0001). Percentage of time awake (**D**, F_5,46_=2.823, P=0.0264), in NREM (**E**, F_5,46_=1. 696, P=0.1545) or REM (**F**, F_5,46_=2.960, P=0.0213) following injection of monomeric αSyn or αSyn-PFF (n=52 values). Average duration of awake (**G**, F_5,46_=1.923, P=0.1088), NREM (**H**, F_5,44_=3.914, P=0.0051) or REM (**I**, F_5,46_=1.426, P=0.2327) events following injection of monomeric αSyn or αSyn-PFF (n=52 values). (**J**) Olfactory discrimination index expressed as the ratio spent in the corner associated with chocolate or TMT odors in mice injected with monomeric αSyn or αSyn-PFF (n=71 values, F_5,63_=5.052, P=0.0006). (**K**) Sucrose preference index expressed as the ratio of water and sugar-water following 24h in mice injected with monomeric αSyn or αSyn-PFF (n=71 values, F_5,65_=3.493, P=0.0074). (**L**) Time spent in the light area of the dark-light test in mice injected with monomeric αSyn or αSyn-PFF (n=71 values, F_5,65_=6.247, P<0.0001). (**M**) Number of fecal pellets produced in 20 min in mice injected with monomeric αSyn or αSyn-PFF (n=71 values, F_5,65_=5.313, P=0.0004). Data are expressed as mean ± SEM. All individual point represents individual animal. * P<0.05, ** P<0.01, *** P<0.001, **** P<0.0001. Statistical data represent post hoc analyses compared to monomeric αSyn group following 1-way ANOVA.

When analyzing sleep pattern in more details, we found a trend to increase the % of time awake and reduced % time in REM but not NREM around the 3^rd^-month post-injection that became significant at 6 months (**Fig. 4D-F**), an effect that was diffused along the dark and light phase (**Fig. S12I, L, O**) but does not seems to be consistent over time. However, this was associated with a trend towards an increase of awake and a significant reduction of the NREM without impacting REM event duration (**Fig. 4G-I**). Interestingly, the number of events as well as the longest duration does not seem to affect awake (**Fig. S12J-K**) or NREM (**Fig. S12M-N**), while it does consistently impact REM by a reduction of events but not the longest duration (**Fig. S12P-Q**). These data suggests that RBD appear simultaneously with the increase of pSer129 in the Pons or the midbrain, including structures that have been associated with sleep regulation such as the DMV, LDT, or the SNc.

### Time-dependent manner propagation of pathologic alpha-synuclein in the central nervous system correlates with other prodromal non-motor phenotypes

In addition to sleep disturbance, in patients, hyposmia is highly correlated with the risk of developing PD (*33*). To test hyposmia in our mice, we performed an adjusted version of the olfactory discrimination test (*34*), where mice were successively exposed to chocolate- or fox urine-smell in a classical open field (**Fig. 4J, S13A-J**). This test allows us to assess olfactory preference for a positive- or a mild anxious odor. Interestingly, we found that αSyn-PFF-injected mice present an impairment in the discrimination index as early as 3 months following injection (**Fig. 4J**) that was also linked with an abnormal increase in activity in the olfactory test as early as the 1^st^-month post-injection (**Fig. S13A-J**). This confirms an hyposmia-like phenotypes at around 3 months post-PFF injection, a period where pSer129 increases in the Parabigeminal nucleus and the dorsal raphe, the two structures involved in innate behaviors and sensory processing (*35*). We next tested whether the injection of αSyn-PFF could induce a depressive-like phenotype using the classical sucrose-preference test (**Fig. 4K, S13K-M**). We found a reduction of the sucrose-preference index in the late stage (∼7 m) with a decreasing trend starting at the 3^rd^ month (**Fig. 4K**) that was due to a reduction of sucrose consumption rather than total intake (**Fig. S13K-M**). In addition, we found an increase in anxiety-like behavior using the dark-light box test (**Fig. 4L, S13N**). Constipation also seems to be one of the earliest pre-motor symptoms and a major predictive factor of PD (*36*). We found a reduction in the number of fecal pellets ejected over 20 minutes in mice injected with αSyn-PFF as early as 2 weeks and significant at late stage compared to control group (**Fig. 4M**); this resulted in a reduction of the total fecal weight (**Fig. S13O**), but not their average weight (**Fig. S13P**). Drying the fecal pellet shows a reduction of the dry fecal pellet (**Fig. S13Q**) but no effect on the water content (**Fig. S13R**) suggesting an impairment of intestine motility rather than constipation, an effect that can be linked to early propagation in the DMV. Altogether, it appears that the spread of pathologic alpha-synuclein from the gut to the brain correlated with the impairment of structures that are involved in non-motor phenotypes observed in early-stages of PD.

### Fine motor planning impairment predates motor symptoms in the gut-brain model

PD is characterized by specific motor symptomatology, including freezing of gait as well as loss of motor coordination. Here, we tested motor symptoms in the high-speed rotarod (40 RPM) in untrained mice and found a reduction of the average time to fall across 3 sessions in the late stage of the model (**Fig. 5A**) at session 3 of testing (**Fig. S14A-C**). To reproduce the time Up&Go test, a classic test employed to assess sequential locomotor tasks, we used the docking setting at 20 rpm, again in untrained mice. Interestingly, we found a significant reduction in the average time to fall across 3 sessions as early as 1 month after injection (**Fig. 5B**) that was consistent in sessions 2 and 3 (**Fig. S14D-F**). This suggests that fine movement sequential planning might be impaired at the earlier stage compared to motor execution (*37*). To confirm such a hypothesis, we tested the mice in the bedding test for 24h (**Fig. S14G**) and found a reduction of the overall score at around 1-month post-injection (**Fig. 5C**). In comparison, testing the mice on the descending pole revealed an increased time to descend after 7 months post-injection (**Fig. 5D**), which was independent of the anxiety effect linked to the test (**Fig. S14H-I**). Similarly, testing the mice on the declined platform revealed a trend toward increased time to descend (**Fig. 5E**) together with a significant clasping phenotype as early as 3 months post-injection (**Fig. 5F**). Finally, we found a reduction of alpha and beta oscillation in the late stages (∼7 months) that was independent of the movement of the animal (**Fig. S14J-S**).

**Fig. 5.**
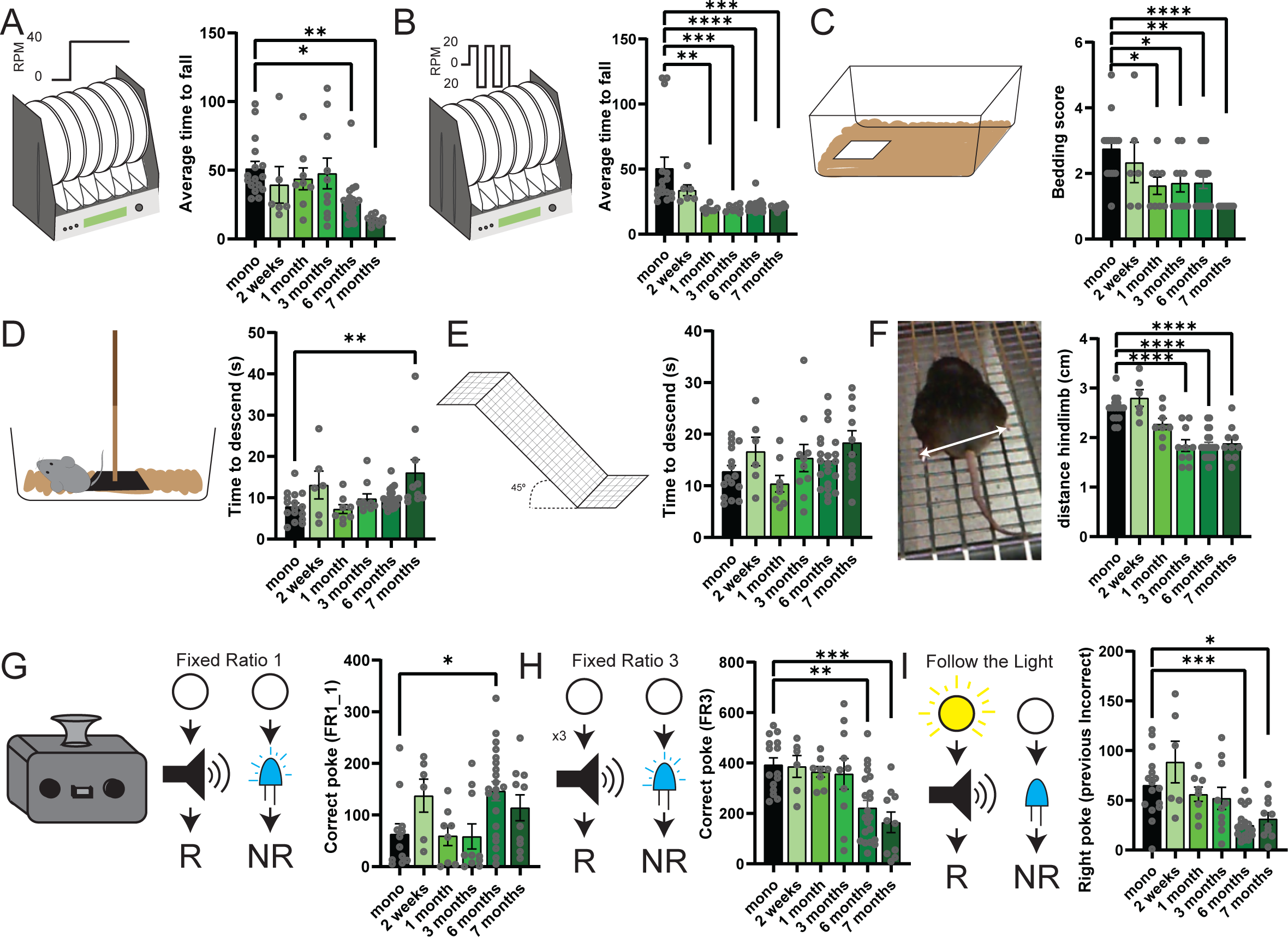
Development of locomotor symptoms following injection of αSyn-PFF in the stomach. (**A-B**) Latency to fall in the 40 RPM constant speed (**A**, n=71 values, F_5,65_=5.123, P=0.0005) or the 20 RPM docking (**B**, n=71 values, F_5,65_=7.601, P<0.0001) rotarod protocol. (**C**) Bedding scores following 24h in mice injected with monomeric αSyn or αSyn-PFF (n=71 values, F_5,65_=6.279, P<0.0001). (**D**) Time to descend the vertical pole in mice injected with monomeric αSyn or αSyn-PFF (n=71 values, F_5,65_=4.312, P=0.0019). (**E**) Time to descend the inclined platform in mice injected with monomeric αSyn or αSyn-PFF (n=71 values, F_5,65_=1.976, P=0.0939). (**F**) Average distance between hindlimbs during the inclined platform test in mice injected with monomeric αSyn or αSyn-PFF (n=71 values, F_5,65_=19.55, P<0.0001). (**G-I**) Number of correct pokes during the first session of fixed ratio 1 (**G**, n=67 values, F_5,61_=3.499, P=0.0076) or fixed ratio 3 (**H**, F_5,64_=6.539, P<0.0001) or in the follow the light protocol (**I**, F_5,65_=7.600, P<0.0001) in mice injected with monomeric αSyn or αSyn-PFF. Data are expressed as mean±SEM. All individual point represents individual animal. * P<0.05, ** P<0.01, *** P<0.001, **** P<0.0001. Statistical data represent post hoc analyses compared to monomeric αSyn group following 1-way ANOVA.

Altogether, these results suggest that fine motor execution might be impaired at an early stage, probably when pSer129 is present in the mesencephalic locomotor region (MLR), while motor function declines when dopaminergic degeneration becomes significant and affects the entire basal ganglia (*38–40*).

### Cognitive declines are linked with a late-stage of the αSyn-PFF gut-brain model

In late-stage PD, cognitive flexibility is strongly reduced and correlates with the dementia phase of the disorder (*41–43*). Here, we used in-home cage cognitive tasks (*44*) to test the ability of the mice to correctly shift between different cognitive paradigms. In the 3 sessions of Fixed-ratio 1 (FR1), we found that mice injected with αSyn-PFF for a long period (∼6-7 months) display a longer time to associate the correct poke with reward delivery (**Fig. 5G, S15A-G**), but still maintain the ability to associate action-outcome. Interestingly, when the cognitive paradigm is modified either changing the intra-dimensional states (FR3, **Fig. 5H**) or the extra-dimensional (follow the light, **Fig. 5I**) mice injected with αSyn-PFF for ∼6-7 months display a significant impairment of cognitive planning (**Fig. S15H-L**).

Altogether, these data suggest that cognitive planning is impaired in parallel with the propagation of pSer129 in the striatum and the frontal cortex, two structures extensively linked with cognitive planning and set-shifting (*45*, *46*).

### The surgery does not affect behavior

The fast arising of some non-motor phenotype alteration, as soon as 2 weeks or 1-month post-surgery suggests that the effects observed could be linked to the recovery rather than the injection of the αSyn-PFF. To rule out such an effect, we compared the effects of the surgery in animals injected with monomeric αSyn after 2 weeks or 7 months (**Fig. S16**). In all non-motor phenotypes tested before, we found no significant differences between animals.

### Sex effects observed in motor behavior recapitulate to some extent PD pathology

Despite the predominance of PD in males, motor and non-motor symptoms are present in both males and females. Interestingly, some of them are predominantly found in females including hallucination, depression, and insomnia (*47*, *48*). To assess whether there is an influence of biological sex in our gut-PFF model phenotypes, we first compared the propagation of pSer129 in function of the group and the sex (**Fig. S17A-H**). Interestingly, we found several structures that were not showing significant group effect (**Fig. 3**) that now present a sex*group effect with a faster propagation of pSer129 in the motor cortex (**Fig. S17A**), striatum, medial septum (**Fig. S17B**) and the Pons (**Fig. S17G**) in male, while the propagation in the DMV appears to be slightly faster in female (**Fig. S17H**). No effect was observed for other analyzed regions (**Fig. S17C-F**). Interestingly, comparing the number of ChAT-positive neurons in the DMV, we found a more significant effect in the female (**Fig. S17I**), while the loss of TH-positive neurons does not present any sex-dependent effects in either the SNc (**Fig. 17J**) or VTA (**Fig. S17K**), despite a non-significant group*sex effect in all 3 structures. However, such effect need to be taken carefully as the sex-effect in PD is often correlated with menopause (*49*).

We next analyzed non-motor phenotypes (**Fig. S18A-M**) and found no significant sex*group effects in sleep (**Fig. S18A-H**), olfactory discrimination (**Fig. S18I-K**), depressive-like behavior (**Fig. S18L**) anxiety-related behavior (**Fig. S18M**). We also observed no sex*group effect in the constipation test (**Fig. S19A-C**). In sharp contrast, we found a significant sex*group effect in the motor test, with the stronger, or at least fastest, impairment in males for the constant speed rotarod (**Fig. S19D-F**), pole test (**Fig. S19K**) or clasping (**Fig. S19N**), while fine motor learning seems to be impaired faster in female (**Fig. S19G-I**). No significant sex*group effect was found in the bedding test **(Fig. S19J)**, the time to descend inclined platform **(Fig. S19M)**, or cognitive flexibility test (**Fig. S20**).

Altogether, these shows that alpha-synuclein propagation and motor impairment are influenced by sex. However, such data need to be interpreted carefully and further characterization is required, including larger group sizes.

### Time-dependent propagation recapitulates various sleep impairment phenotypes

Following injection of αSyn-PFF in the stomach, we found impairment of sleep including impairment of sleep stability as well as REM-sleep behavior disorder around 3-month post injection (**Fig. 4**), like the parallel propagation of pSer129 in the SNc (**Fig. 3**). In particular, a reduction of dopamine is observed in patient with idiopathic RBD (*50*, *51*). To test whether alpha synuclein expression in the SNc might impact the sleep pattern, we used viral overexpression of human αSyn (AAV-hSNCA) or control (AAV-mCherry) in the SNc (**Fig. 6A**). The overexpression of human αSyn (hαSyn) does not cause αSyn aggregation with endogenous wildtype (WT) αSyn while allowing somatic and axonal expression (**Fig. 6B**). Following bilateral injection, mice were implanted with a telemetry head stage and sleep was scored 8 weeks post-injections. Interestingly, we found no modification of the time awake following injection (**Fig. 6C-F**), the injection increased the time in REM (**Fig. 6G**), without impacting the event pattern outside of longest REM event (**Fig. 6H-J**). Not surprisingly, the increase of time in REM with overexpression of hαSyn in the SNc induces a reduction of the time in NREM (**Fig. 6K-N**). Finally, analyses of the number of RBD events (**Fig. 6O**) as well as the normalized RBD events (**Fig. 6P**) suggest that overexpression in the SNc is sufficient to increase RBD significantly without affecting muscle activity (**Fig. 6Q**). Altogether, this suggests that overexpression of hαSyn in the SNc is sufficient to alter atonia during REM sleep (*52*, *53*).

**Fig. 6.**
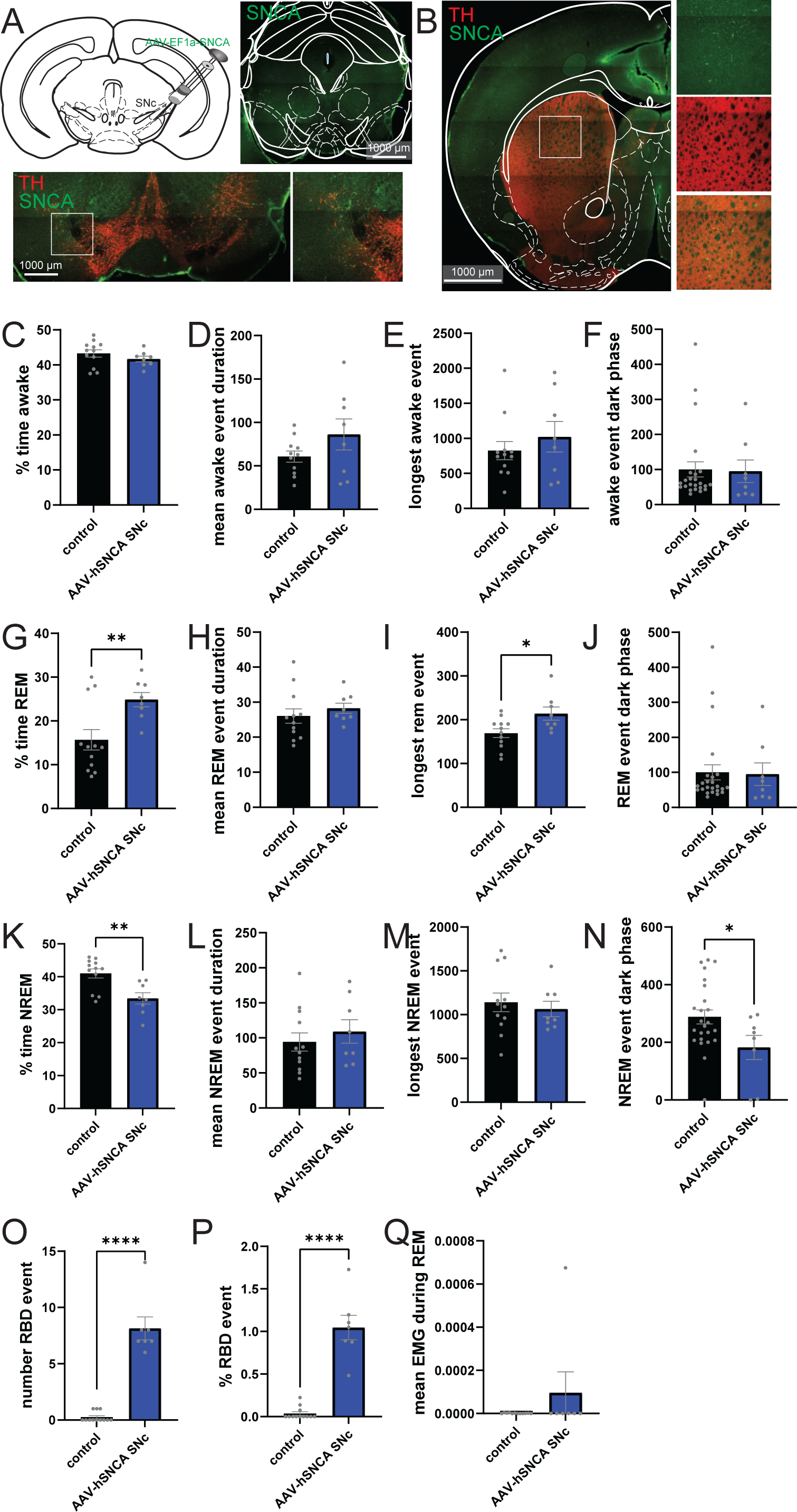
Sleep alteration with overexpression of alpha-synuclein in the SNc. (**A, B**) Confocal images of the injection of AAV-hSNCA in the SNc of WT mice showing expression of αSyn in inputs structure of the SNc **(A)** and output (i.e. striatum, **B**) repeated for all mice. (**C**) Percentage of time awake during the 24h recording in control mice (AAV-mCherry) or following injection of AAV-hSNCA in the SNc. (**D**) Average awake event duration during the 24h recording in control mice (AAV-mCherry) or following injection of AAV-hSNCA in the SNc. (**E**) Duration of the longest awake event during the 24h recording in control mice (AAV-mCherry) or following injection of AAV-hSNCA in the SNc. (**F**) Average time awake during dark phase during the 24h recording in control mice (AAV-mCherry) or following injection of AAV-hSNCA in the SNc. (G) Percentage of time REM during the 24h recording in control mice (AAV-mCherry) or following injection of AAV-hSNCA in the SNc. (**H**) Average REM event duration during the 24h recording in control mice (AAV-mCherry) or following injection of AAV-hSNCA in the SNc. (**I**) Duration of the longest REM event during the 24h recording in control mice (AAV-mCherry) or following injection of AAV-hSNCA in the SNc. (**J**) Average time REM during dark phase during the 24h recording in control mice (AAV-mCherry) or following injection of AAV-hSNCA in the SNc. (**K**) Percentage of time NREM during the 24h recording in control mice (AAV-mCherry) or following injection of AAV-hSNCA in the SNc. **(L)** Average NREM event duration during the 24h recording in control mice (AAV-mCherry) or following injection of AAV-hSNCA in the SNc. (**M**) Duration of the longest NREM event during the 24h recording in control mice (AAV-mCherry) or following injection of AAV-hSNCA in the SNc. (**N**) Average time NREM during dark phase during the 24h recording in control mice (AAV-mCherry) or following injection of AAV-hSNCA in the SNc. (**O**) Number of RBD events during the 24h recording in control mice (AAV-mCherry) or following injection of AAV-hSNCA in the SNc. (**P**) Percentage of RBD events over the number of REM events during the 24h recording in control mice (AAV-mCherry) or following injection of AAV-hSNCA in the SNc. (**Q**) Average EMG signal during REM events during the 24h recording in control mice (AAV-mCherry) or following injection of AAV-hSNCA in the SNc. Data are expressed as mean±SEM. All individual points represent individual animals. * P<0.05, ** P<0.01, **** P<0.0001. Statistical data represent unpaired t-test analyses.

### Somatic versus axonal propagation of alpha-synuclein in dopaminergic neurons

Viral overexpression of αSyn in the SNc suggests its role in the development of RBD. However, in PD, despite the early propagation of alpha-synuclein in the soma, degeneration seems to occur first in the terminals located in the striatum (*54*, *55*). To assess whether αSyn seeding in the striatum or the SNc impacts different PD-related phenotypes, we injected WT mice with αSyn-PFF in the SNc or the striatum. As a control, a group of mice was injected with monomeric αSyn in either the SNc or the striatum. The unilateral injection of αSyn-PFF in the SNc (**Fig. 7A-C**) is sufficient to increase pSer129 staining after 1 month in the cortex in sharp contrast to pSer129 staining observed following injection in the stomach (**Fig. 3**) and to induce a strong reduction of TH-expressing neurons in the SNc (**Fig.S21A-C**) but not the VTA (**Fig. S21D-F**), interestingly correlating the number of TH-positive neurons to the number of DAPI positive neurons revealed no changes, suggesting a loss of TH-neurons rather than decrease expression of TH (**Fig. S21C, F**). Similar unilateral injection in the striatum is sufficient to increase pSer129 in the motor cortex, and other structures (**Fig. 7D-F**) with a reduction of TH-positive neurons in the SNc more limited than the one observed in SNc injected animals (**Fig. S21A-F**). Recently, the increased expression of the apolipoproteins, in particular, ApoE, in dopaminergic neurons has been suggested to participate in αSyn spreading (Paslawski et al., 2019). While ApoE is synthesized mainly by astrocytes, microglia and immature neurons, neuronal ApoE levels are increased in injured or stressed neurons and therefore postulated to play a role in the growth and repair of cell in the CNS (*56*). Co-staining for TH and ApoE shows that injection of αSyn-PFF in the striatum increases ApoE inclusion within the SNc (**Fig. S21G-J**) while the expression of ApoE inside TH+ neurons seem to increase no matter if the injection of αSyn-PFF was in the SNc or the striatum (**Fig. S21K**), showing similarity between the used model and data obtained from PD patient’s tissue.

**Fig. 7.**
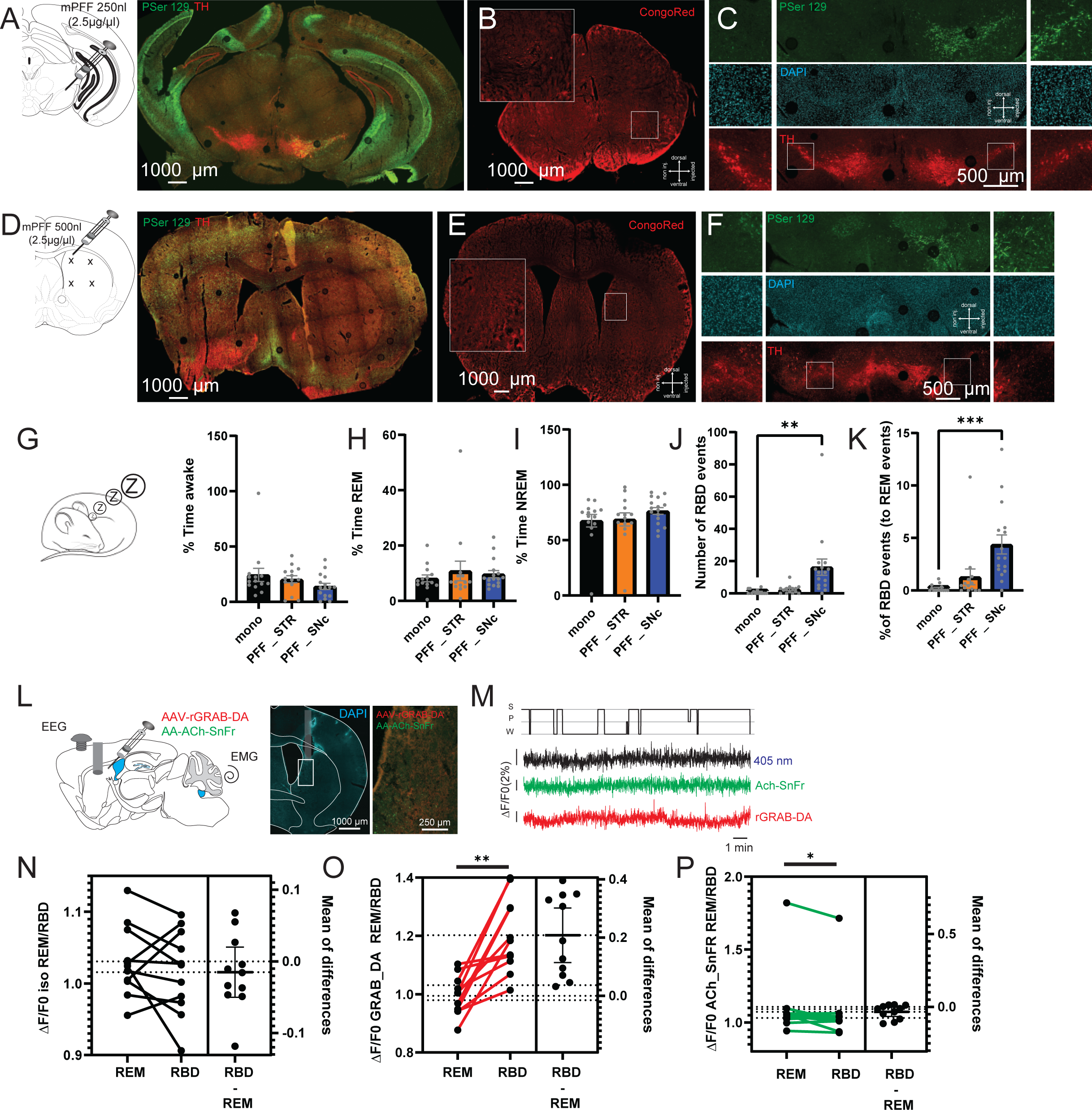
REM-sleep behavior disorder is paired with alpha-synuclein aggregates in the SNc. Unilateral injection of αSyn-PFF in the SNc (**A-C**) or the striatum (**D-F**) revealed the presence of amyloid-like structures (**B, E**) in the vicinity of the injection site brain. The SNc injection at the soma level (**C**) or their terminals (**F**) affect TH expression differently. (**G-I**) Percentage time awake (**G**, n=44 values, F_2,41_=1.660, P=0.2026), REM (**H**, n=44 values, F_2,41_=0.3546, P=0.7036) or NREM (**I**, n=44 values, F_2,41_=0.9903, P=0.3802) following injection of monomeric αSyn (black, or αSyn-PFF in the striatum (orange) or the SNc (blue). (**J-K**) Number of RBD events (**J**, n=44 values, F_2,41_=7.100, P=0.0023) and normalized RBD event to REM events (**K**, n=44 values, F_2,41_=9.625, P=0.0004) following injection of monomeric αSyn (black, or αSyn-PFF in the striatum (orange) or the SNc (blue). (**L**) Schematic representation of the fiber photometry experiments with recording of dopamine and acetylcholine release in the dorsomedial striatum (DMS) during sleep. Confocal images of the implantation of the optic fiber and the spread of the adenoviruses. (**M**) Example of traces showing sleep scoring within 20s windows, isosbestic, Ach-SnFr and rGRAB-DA signal in the DMS. (**N**) Variation of the isosbestic signal in the DMS between REM and RBD events in mice injected with αSyn-PFF in the SNc (n=11 mice, paired t-test t_10_=0.9574, p=0.3609). (**O**) Variation of the rGRAB-DA signal in the DMS between REM and RBD events in mice injected with αSyn-PFF in the SNc (n=11 mice, paired t-test t_10_=4.936, p=0.0006). (**P**) Variation of the Ach-SnFr signal in the DMS between REM and RBD events in mice injected with αSyn-PFF in the SNc (n=11 mice, paired t-test t_10_=2.318, p=0.0429). Data are expressed as mean±SEM. All individual point represents individual animal. * P<0.05, ** P<0.01, *** P<0.001. Statistical data represent post hoc analyses compared to monomeric αSyn group following 1-way ANOVA or paired t-test for photometry data.

Altogether, these data confirmed that injection of αSyn-PFF at the somatic or axonal level induces retrograde and anterograde spreading of pathologic αSyn as well as neuronal degeneration through different mechanisms (*57*).

Next, we assessed the effects of αSyn-PFF bilateral injection on mouse behavior. Animals were tested every 2 weeks following injection. In the constant speed rotarod, we found that injection of αSyn-PFF in the striatum induced a fast reduction of the time to fall (∼4^th^ week post-injection) while this impairment was delayed in the mice with PFF injection in the SNc (∼6 weeks, **Fig. S22A-D**). In the docking rotarod test, we found that the time to fall decreased in both groups very quickly (∼4^th^ week, **Fig. S22E-H**). Interestingly, when tested in fixed ratio 1 task, we found that the number of incorrect (**Fig. S22M-P**), but not correct (**Fig. S22I-L**), as well as the number of pellets collected (**Fig. S22Q-T**) is altered only in SNc injected animals at 4 weeks after injection, while at 6 weeks post-injection in striatum-injected mice.

Altogether, these data suggest that targeting the soma or the terminals of dopaminergic neurons with αSyn-PFF impact differently behavioral outcomes linked with motor and cognition function.

### Targeting soma of dopaminergic neurons impacts REM-sleep atonia, but not sleep pattern by modulating release in the dorsomedial striatum

Following 8 weeks post-injection, sleep patterns were recorded using our telemetry implant. We found no significant difference in the pattern of awake (**Fig. 7G, S23A-C**), REM (**Fig. 7H, S23D-F**), or NREM (**Fig. 7I, S23G-I**) events following injection of αSyn-PFF in both the striatum and the SNc. However, quantification of the number of REM events with atonia suggests an increase of RBD events exclusively in mice injected with αSyn-PFF in the SNc (**Fig. 7J-K**) despite the absence of effects on the EMG signal during REM sleep (**Fig. S23J**). To test whether the effects observed were due to alteration of dopaminergic release in the striatum, we injected the animals with a mix of adenovirus, the first one green-shifted to measure acetylcholine release (AAV-Ach-SnFr) and the second one red-shifted to measure dopamine release (AAV-rGRAB-DA) together with EEG and EMG (**Fig. 7L**). Based on a previous report on the role of dopamine release in the dorsomedial striatum (DMS) during sleep, an optic fiber was then placed above the injection site (*58*, *59*). We next recorded sleep and photometry, including a control isosbestic signal. Sleep schedule was then scored on 20s windows and each photometry signal average for each sleep-scored window (**Fig. 7M; Fig S23K-R**). As isosbestic signal in control, striatum or SNc injected animals were not different during awake (**Fig. S23K**), REM (**Fig. S23N**), or NREM (**Fig. S23Q**). However, pooling together their average to compare to other signal variations, during awake events showed a significant increase in dopamine release in all three groups compared to isosbestic signal (**Fig. S23L**) with a highest increase found in SNc-injected animals compared to control mice. In the same context, we found that acetylcholine release in the striatum increased in control, but not αSyn-PFF injected mice compared to isosbestic signal (**Fig. S23M**). During REM events, we found that classically dopamine release is reduced in control animals as well as animals injected with αSyn-PFF in the striatum. Interestingly, the release of dopamine in DMS remains high in animals injected with αSyn-PFF in the SNc (**Fig. S23O**). In contrast, while acetylcholine release in the striatum during REM events does not change if αSyn-PFF are injected in the SNc compared to controls, we found that injection of αSyn-PFF in the striatum is sufficient to reduce the cholinergic signal compared to monomeric αSyn injected animals (**Fig. S23P**). Finally, during NREM events, we found that dopamine release in the striatum does not change significantly compared to isosbestic variation or monomeric αSyn injected mice if the αSyn-PFF are injected in the SNc, while dopamine release decrease in animals injected in the striatum compared to isosbestic variation (**Fig. S23R**). Finally, we found that acetylcholine release decreases slightly in monomeric αSyn injected animals compared to isosbestic variation, but not αSyn-PFF injected animals during NREM events (**Fig. S23S**).

Altogether, these confirmed that the different spread of pathologic αSyn following injection at the soma or terminals level of dopaminergic neurons impacts the modulation of activity and/or axonal release differently.

As animals injected with αSyn-PFF in the SNc demonstrate an increase of dopamine release during REM events compared to isosbestic as well as control mice signals together with a reduction of atonia during REM events (RBD), we hypothesized that the effect could be explained by movement during REM events. To assess that, we compared photometry signals during REM and RBD events in SNc-injected animals (**Fig. 7N-P**). Not surprisingly, we found no significant difference in isosbestic signal during REM and RBD (**Fig. 7N**). In both dopamine (**Fig. 7O**) and acetylcholine signals (**Fig. 7P**), we found a significant difference between REM and RBD, where dopamine signal increases strongly during RBD events, while acetylcholine signal decreases. Interestingly, in basal condition, cortical activity changes during movement (**Fig. S14J-S**), suggesting that parasitic movement observed during RBD can also be encoded at the cortical level. We next compare cortical oscillation during REM and RBD events in mice injected with αSyn-PFF in the SNc (**Fig. S23T-X**). Despite the presence of movement during RBD, we found no significant variation of cortical bands frequency between REM and RBD (**Fig. S23T-X**).

Altogether these suggest that movement that occurs during RBD is linked with the activity of SNc dopaminergic neurons, and thus dopaminergic neuron inputs modulate the firing rate rather than cortical modulation of movement.

## DISCUSSION

Our investigation into the impact of αSyn pathology on the gastrointestinal tract, central nervous system, and associated non-motor phenotypes in the context of PD has yielded substantial insights with significant implications. The multifaceted nature of our study has allowed us to draw conclusions that bridge the gap between the gastrointestinal origin of PD, alpha-synuclein propagation, and the manifestation of non-motor symptoms in PD, particularly RBD-like behaviors. Crucially, we were able to highlight a simultaneous spread of αSyn from the periphery to the brain with the temporal development of motor and non-motor symptoms, suggesting that the alteration of specific brain structures increased levels of pathological αSyn is correlated with specific behavioral impairments (*60*) in light with the body first hypothesis of idiopathic PD (*30*). First, our data support the notion that pathological αSyn introduced via stomach injections of αSyn-PFF propagates from the stomach via the vagus nerve and then towards the brain (central manifestation) (Kim *et al.*, 2019) or towards peripheral tissues (colon, peripheral manifestations). This bidirectional manifestation was demonstrated *ex vivo* as well as *in vivo* (*61*, *62*). This phenomenon not only correlates with alterations in gut microbiota, mirroring some of the changes observed in PD patients but also suggests a potential mechanistic link to the non-motor manifestations of the disease. The direction of αSyn propagation is of high interest, as it does define the possible model and so, possibility to use imaging for prodromal forms of PD. Previous studies in human tissue focusing on the pattern of Lewy pathology propagation suggested that the spread does occur primarily in a retrograde fashion (*63*, *64*). Our data seems to confirm such propagation, where αSyn is first taken up by neurons somewhere along the axon or in the presynaptic terminal. This is particularly visible following injection of PFF in the striatum, with a strong staining for pSer129 was found in the amygdala. However, the presence of aggregated αSyn markers in the colon also suggests that the spread may also occurs anterogradely most probably to a lesser extent. This still raises the question of the circuit involved in the αSyn transmission from the enteric nervous system to the brain. In recent reports using αSyn-PFF injection in the duodenum, a vagotomy was sufficient to prevent most of the phenotypes linked with αSyn aggregation (*12*, *65*). Interestingly, similar injection in the rat do not result in spinal cord αSyn pathology suggesting a possible propagation mostly by the parasympathetic system (*66*). However, some of these data have been difficult to reproduce and possible bidirectional propagation in both system seems likely (*67*).

Moreover, our findings highlight the critical role of αSyn in the gut-brain axis, demonstrating that the spread of aggregated αSyn from the gut to the brain is closely associated with the impairment of brain regions involved in non-motor phenotypes observed in PD (*68*). This raise the question of a possible degenerative path between the gut and the SNc involving several brain structures and their respective functions (*26*). However, in our whole brain mapping, not all structures were found to accumulate αSyn along a caudal-to-rostral pattern. In particular, it is interesting to find that the Gigantocellular and the medullary nuclei, both located near the DMV and connected with it (*69*) were presenting an increase of pSer129 staining in very late stages of the model. This highlight the selective neuronal vulnerability in Parkinson’s disease (*70*).

Furthermore, our study elucidates the temporal dynamics of αSyn propagation, revealing concurrent presence of the phosphorylated form in key brain regions associated with sleep duration, motor function, and cognitive planning. Among them sleep disturbances are particularly noteworthy, affecting a substantial proportion of individuals with PD. Our data, suggest that RBD-like behavior, characterized by an absence of atonia during REM-sleep, robustly occurs in the gut-brain PFF model. A previous study, demonstrated that injections of αSyn-PFFs or inactivation of neurons in the sublaterodorsal tegmental nucleus (i.e., the subcoeruleus nucleus in humans) can induce RBD-like behaviors in animals(*71*). The authors also found transneuronal spread of α-synuclein pathology to several other brain regions, and the mice developed parkinsonian locomotor dysfunction, depression-like disorder, olfactory dysfunction and gastrointestinal dysmotility demonstrating a potential path for phenoconversion of RBD to motor signs of parkinsonism. In another study, using A53T SNCA transgenic mice, RBD-like behaviors occurred at 5 months of age (*72*). The RBD-like behaviors correlated with pSer129 expression in the sublaterodorsal tegmental nucleus along with nuclei in the ventromedial medullary reticular formation and the pedunculopontine nuclei. In the present study, we have not specifically targeted the sublaterodorsal tegmental nucleus, but our work highlights the function of dopaminergic structures in RBD-like behaviors. Interestingly, targeting nigral dopaminergic neurons or their striatal terminals exhibit no comparable influence on sleep structure when contrasted with stomach injections while recapitulating RBD. This might imply that the Substantia Nigra pars compacta per se or structures connected with this region are implicated in RBD-like behaviors. Moreover, our data using Fiber photometry suggest that while dopamine release in the dorsomedial striatum is involved in RBD generation, acetylcholine release might be crucial for wakefulness and switch between REM and NREM sleep. Crucially, while the majority of Acetylcholine in the striatum is originating from interneurons, a subset is released from cholinergic neurons of the brainstem including the Pedunculopontine and the Laterodorsal Tegmental nuclei (*73*), two structures involved in slow-wave cortical activity and sleep (*74*).

Interestingly, PD patients present Lewy-bodies like inclusion in peripheral organs including skin, heart and digestive tract, which correlate with excessive sweating, hypotension, urinary incontinence or constipation (*2*). In recent years, accumulation of αSyn in the digestive tract is observed in early stages of the disease (*75*) and suggest an implication of intestinal microbiota (*15*). While it is unknown whether the modification observed in microbiota facilitate αSyn aggregation or whether aggregation of αSyn in specific intestinal cell type modify the environment to favor the development of specific microbiote (*76*), the outcome of metanalyses seems to be coherent (*77*). In several studies of bacterial family, bifidobacteriacea, Rikenellaceae, Lactobacillae and Verrucamicrobiaceae were increased in PD patients compare to healthy controls, while Lachnospiraceae, Prevotellaceae were reduced (*77*). Interestingly, in our study in mice, we found a reduction of Lachnospiracea as early as 1 month following injection of αSyn-PFF in the gut while Lactobacillae were reduced within the same period. Similarly, at the genus level multiple studies found increase of Bifidobacterium, Lactobacillus and Akkemansia, while Blautia, Facaelibacterium and Rosaburia were decrease (*77*). In our analyses, we were able to find an increase of Lactobacillus within one month of the injection and a reduction of Blautia following 7 months post injection, suggesting that despite significant differences between human and rodent microbiota, some similarities between our model and PD patients. It is interesting to note that in our study, we found several families linked with anti-inflammatory functions that were decrease in mice injected for as less as 2 weeks, including Prevotellaceae, suggesting that αSyn-PFF injection might impair gut-barrier function.

The αSyn-PFF model with the injection in the muscularis layers of the stomach and the duodenum appear as a clinically relevant model, but often failed to show progression of the disease (*66*) or the propagation beyond the Pons (*65*). While most studies were initially performed in rats, the same approach done in mutant mice confirm the pathology. (*78–80*) In our hands, we found that the variance of the phenotypes and immunostaining differs based on the sex of the animals, as well as the time following injection. One limitation of our approach is that the laparotomy during PFF injection surgery induce a local inflammation that does not allow to investigate earlier than 2 weeks the spread at the injection site, nor it is not possible to exclude that the injection induces local bleeding and thus partial propagation through the blood stream. A second limitation is that, while pSer129 is considered as a marker for αSyn aggregation, the presence of a small but quantifiable amount in the central nervous system (*20*) that participate in synapses function (*6*) make it difficult to estimate possible increases in the early timeframe. In particular, this is enhanced by the different morphologies of pSer129 species observed using immunofluorescent versus widefield microscopy (*19*). In order to overcome that limitation, all our experiments included a group of animals injected similarly with monomeric αSyn, as well as all mouse tissues were processed and labelled at the same time using same solution. Finally, the last limitation of our study is the possible role of the age in αSyn spreading (*13*). In their work, Challis and colleagues found that aged animals (16 months or older) display a age related propagation of αSyn as well as progression of phenotypes. While this is most likely the case, as native αSyn increase with age (*81*), and thus increasing the seeding between native αSyn and fibrils, it was not possible for our experiments due to the age limit and the aversive effects of the surgery. To overcome a possible age-related effect, we used age-matched mice and behavioral testing at the same time, where only the age at the time of injection differ across experimental groups, but the older group (2 weeks post-injection) was including animals 9 to 12 months old. In addition, we divided our control groups with the injection of monomeric αSyn for 2 weeks or 7 months in order to avoid the possible effect linked with the surgery. It would be of interest to use other models of animals that either present with abnormal αSyn expression or that can live past the ∼2 years.

In conclusion, our findings offer a novel and deepened perspective on the etiology and progression of PD, emphasizing the interconnectedness of gastrointestinal neurological, and behavioral aspects (*12*). The identification of specific regions implicated in the αSyn propagation cascade opens new avenues for targeted diagnosis and therapeutic strategies. Our research not only expands the current understanding of PD but also provides a solid foundation for the development of interventions based on the mice model.

## MATERIALS AND METHODS

### Study design

A comparable number of animals in similar studies was used for the entire experiments. No power analyses were done for sample size estimation. Data exclusions were defined by animal that complete the entire study, quality of tissue collected (bad perfusion, noise in immunostaining signal) as well as statistical exclusion analyses. Data collection timepoint were defined before the beginning of the experiment as well as allocation of animals for each group. Sex was balanced between group. Group design and behavior analyses were defined before the beginning of the experiment, tissue collection and analyses were changed during the study. To reduce inter group variation, all experiments were completed at the same period of the day, in a group of animals of same age, in a similar environment and with experimenter/animal caretaker blind to animal group allocation.

### Chemicals and antibodies

Unless stated otherwise, all chemicals were purchased from Sigma Aldrich (Merck KGaA, Germany) and were of an analytical grade. All solutions were prepared using Milli Q deionized water (Millipore, MA, US).

### Animals

Wild-type C57Bl6 mice were bred and maintained in the animal facility at the Karolinska Institute, Sweden, adhering to the guidelines set forth by the local ethical committee at Karolinska Institute (3218–2022, 24297–2022) and the European Communities Council Directive (86/609/EEC). To minimize within-experiment variability, experimental animals and their respective controls were sourced from the same litters, ensuring identical age and housing conditions.

### αSyn purification and preparation of the αSyn-PFF

aSyn-PFF were prepared similarly to previous publication(*12*) Detailed protocol can be found in supplementary material and methods and dx.doi.org/10.17504/protocols.io.dm6gp3nm5vzp/v1. When required for injection αSyn-PFF were diluted to 2mg/ml by adding sterile PBS and sonicated. For all αSyn-PFF preparation, a small aliquot was proceeded for transmission electron microscopy to validate αSyn-PFF quality, while the remaining was used for injection. See dx.doi.org/10.17504/protocols.io.dm6gp3nm5vzp/v1.

### Stereotaxic surgery

Animals were injected bilaterally with virus and αSyn-PFF for behavior and electrophysiology experiments, or unilaterally for anatomy mapping. Viral vector or αSyn-PFF were injected using a Hamilton Syringe. Overexpression of hSNCA was obtained with injection of AAV6-CAG-a-Synuclein. All injections were performed for a period of 10 minutes followed by 5 minutes of diffusion time. Stereotaxic injections were aiming to inject in the Substantia Nigra pars compacta (SNc, 250nl, AP: −3.6mm, ML1.5mm, DV: 4.2mm) and striatum (Str, 500nl, AP: +0.5mm, ML: ±1 and 2mm, DV: 2.5 and 3.5mm). All coordinates were defined from the Bregma and the surface of the brain.

### Intra-muscularis layer injection in stomach and proximal-duodenum

At 2 months of age, two discrete PFF injections (2.5 µl each) were administered along the lesser curvature of the stomach and two additional injections (2.5µl each) in the muscularis layers of the duodenum adjacent to the sphincter.

### Tissue processing

After behavioral and physiological experiments, all animals underwent transcardial perfusion. Post-perfusion, various tissues, including the brain, spinal cord, kidney, liver, lungs, eyes, heart, stomach, intestine, and additional skin from the thoracic region, were harvested, and immersed in 4% PFA for post-fixation, a duration not exceeding 4-6 hours. During the processing phase, all tissues were gently dried and subsequently embedded in an optimal cutting temperature (OCT) block prior to sectioning at a thickness of 40 µm using a cryostat.

### Immunostaining – Antibody list

For fluorescent staining, primary antibody were as followed : Tyrosine Hydroxylase (TH, raised in Rabbit, 1:1000, AB152, Sigma-Aldrich), TH (raised in chicken, 1:1000, AB76442, Abcam), Choline acetyltransferase (ChAT, raised in goat, 1:1000, AB144P, Abcam), Alpha synuclein phosphorylated on serine 129 (pSer129, raised in Rabbit, 1:500, AB51253, Abcam), apolipoprotein E (ApoE, raised in goat, 1:500, 178479, Millipore).

Secondary antibody were as follow: 488-Anti Rabbit (raised in Donkey, 1:1000, A2206, Jackson Immunoresearch); CY3-Anti Rabbit (raised in Goat, 1:1000, A11011, Thermofisher); CY5-Anti Rabbit (raised in Goat, 1:1000, A21244, Thermofisher); 488-Anti Chicken (raised in Goat, 1:1000, A11039, Thermofisher); CY3-Anti Chicken (raised in Donkey, 1:1000, A78951, Jackson Immunoresearch); CY5-Anti Chicken (raised in Donkey, 1:1000, A21449, Jackson Immunoresearch); CY3-Anti Goat (raised in Donkey, 1:1000, A11058, Jackson Immunoresearch); CY5-Anti Goat (raised in Donkey, 1:1000, A214447, Jackson Immunoresearch); 488-Anti Mouse (raised in Donkey, 1:1000, a21202, Thermofisher); CY3-Anti Mouse (raised in Donkey, 1:1000,715-175-150, Jackson Immunoresearch); CY5-Anti Mouse (raised in Donkey, 1:1000, 715-165-150, Jackson Immunoresearch); Biotin-conjugated goat Anti-Rabbit (1:500, B-6648, Sigma Aldrich).

### Immunostaining – Fluorescent

Freshly sectioned slices of tissue was first washed 2-3 times with 1X PBS to remove OCT. Next, sections were blocked with 5% Normal donkey/Goat serum in 0.3% TRITON X-100 in 1X PBS for 1h at room temperature before washing 5 times in 1X PBS. Following washes, all sections were transferred to a primary antibody solution containing a mix of primary antibody diluted in 0.3% TRITON X-100 in 1X PBS with 1% Normal Donkey/Goat serum overnight at 4°c temperature. The following day, sections were washed 5 times in 1X PBS before to be transferred in secondary solution containing the adequate secondary antibody diluted in 0.3% TRITON X-100 in 1X PBS with 1% Normal Donkey/Goat serum for 4h at room temperature. Following staining, sections were washed 3 times in 1X PBS before to be mounted on microscope slides and mounted with medium (Vectashield with 4’,6-diamidino-2-phenylindole (DAPI), H-1800-10). To avoid variability in staining linked with the protocol, all animals from the same experiments (Gut injection, AAV-αSyn or αSyn-PFF injection in SNc/STR were stained at the same time with their respective controls.

### Immunostaining – DAB

Freshly sectioned slices of tissue were first washed 2-3 times with 1X PBS to remove OCT. Next sectioned were quenched for 15 minutes in 3ml of quenching solution (0.1ml 30% H_2_O_2_, 0.1ml methanol and 0.8 ml 1X PBS) then washed 4-5 times in 1X PBS. Next, sections were blocked and stained with primary antibody similarly to fluorescent staining (pSer129 primary antibody 1:500) overnight at 4°c. The following day, sections were washed 4-5 times with 1X PBS and transferred into secondary solution for 2h at room temperature. Next, sections were washed 4-5 times in 1X PBS and transferred into ABC Kit solution (PK4000, Vector Laboratories) containing 10µl of solution A and 10µl of solution B for 1ml of 1X PBS. Sections were incubated for 1h at room temperature with ABC Kit solution and quickly washed 4-5 times in 1X PBS. Finaly, sections were transferred to the DAB working solution (SK-4100, Vector Laboratories) under a fume hood. Well-plate containing sections were gently shaken during staining and the reaction was stopped with transfer to 1X PBS based on dark-signal intensity on the fastest arising staining (i.e. limiting factor is group with strongest pSer129 staining). Sections are then washed 3-5 times in 1X PBS and quickly mounted on microscope slides. Following drying overnight at room temperature, slides were then transferred to sequential baths of distilled water (∼2min), 70% ethanol (2 times ∼2min), 95% ethanol (2 times ∼2min), 100% ethanol (2 times ∼2min) and then 10% xylene (2 times ∼5min) to allow section dehydration. Following the last bath, sections were dried at room temperature (∼5 min) and covered with DPX mounting medium). To avoid variability in staining linked with the protocol, all animals from the same experiments (Gut injection, AAV-αSyn or αSyn-PFF injection in SNc/STR) were stained at the same time with their respective controls. See dx.doi.org/10.17504/protocols.io.n2bvj34zplk5/v1.

### Immunostaining – CongoRed

Freshly sectioned slices of tissue were first washed 2-3 times with 1X PBS to remove OCT. Next, all sectioned were mounted on microscope slides and left to dry at room temperature overnight. Next slides were transferred into a 1% CongoRed solution for 30 minutes, followed by alkaline bath (1% Sodium hydroxide in 50% ethanol) for 5 minutes. Slides were then transferred in 3 sequential baths of water to remove excess staining. Slides were then transferred into 0.4% Toluidine blue solution for 10 minutes before being washed in 3 sequential baths of water to remove excess staining. Finaly slides were dehydrated using sequential baths of distilled water (∼2min), 70% ethanol (2 times ∼2min), 95% ethanol (2 times ∼2min), 100% ethanol (2 times ∼2min) and then 10% xylene (2 times ∼5min). Following the last bath, sections were dried at room temperature (∼5 min) and covered with DPX mounting medium). To avoid variability in staining linked with the protocol, all animals from the same experiments (Gut injection, AAV-αSyn or αSyn-PFF injection in SNc/STR) were stained at the same time with their respective controls. See dx.doi.org/10.17504/protocols.io.4r3l22k1jl1y/v1.

### Imaging – Confocal

Sections that were mounted and covered underwent imaging using confocal microscopes (Carl Zeiss LSM 880). All sections were captured at a 10X magnification in air, featuring a resolution of either 1024×1024 or 2048×2048 pixels for high-resolution images. Tile scans were subjected to processing with a 0.6 zoom factor and 10% overlap for automated reconstruction. Z-stack imaging was performed with 1 to 4 µm spacing, and stack projection was executed using ImageJ (https://imagej.net/, RRID:SCR_003070), excluding approximately 10% of the section’s surface. Subsequently, all sections underwent processing in ImageJ software, incorporating consistent steps such as overlapping adjustments, cropping, color correction, and color attribution. It is noteworthy that the post-processing of images for sections within the same experiment, specifically those involving gut injection, AAV-αSyn, or αSyn-PFF injection in SNc/STR, adhered to identical protocols and parameters.

### Imaging – bright field

Mounted and covered sections were imaged with a slide scanner (Hamamatsu, NanoZoomer) with a 40X magnification in air. All sections were extracted using the NDP.view software and saved as tiff file.

### Image processing – whole brain DAB

TIF files extracted from NDP.view software were imported into ImageJ for further analysis. Sections corresponding to distinct brain structures, such as the Prefrontal cortex, striatum, GPe, Thalamus, SNc, Pons, Cerebellum, and DMV, were merged, cropped, and aligned within ImageJ.

To prevent alterations in pixel size, adjustments were made in both the x and y directions. Only complete sections (with the exception of the cerebellum, where most lobes are folded or damaged) were included in subsequent analyses. After positioning each section accurately, files were saved in designated folders for subsequent analyses. Utilizing a custom Matlab script, images from specific folders were opened, converted to an 8-bit grayscale (0-255 scale), and normalized to the signal across the entire images. This normalization process aimed to reduce signal variations associated with the transition between background and brain tissue.

To facilitate adequate structural overlapping, pixels were downsampled using a 5×5 filter, enabling the definition of regions of interest (ROIs) at approximately 30 µm. For each section within a specific group (monomeric αSyn, 2 weeks, 1 month, 3 months, 6 months, or 7 months), specific ROIs were then averaged for density plots and compared between groups using a 1-way ANOVA. Post hoc analyses were subsequently conducted for significant ROIs displaying group effects. The resulting images were plotted with a consistent scale bar using the imagesc – Jet Matlab function.

### Cell counting

Following immunostaining for TH, ChAT, and DAPI, 2-3 sections encompassing the SNc/VTA, DMV, or LDTg were collected for high-resolution scanning (2048×2048 pixels) utilizing tile scanning and z-stack acquisition. Subsequently, images were imported into ImageJ, subjected to maximum average projection, and color-adjusted to facilitate the differentiation of distinct anatomical structures.

For each section, the quantification of TH/ChAT/DAPI-positive neurons was manually performed using the multi-selection tool, and the counting area’s surface was determined using the selection tool. The number of cells in the SNc, VTA, or DMV was then normalized by the surface area to derive the relative density.

Regarding ApoE quantification, 3-4 high-magnification images were acquired for each section within the SNc for each animal. Within the SNc region delineated by TH staining, the manual counting involved TH-positive neurons, ApoE puncta, and TH-positive neurons with at least one ApoE punctum within the cell.

### Cluster counting

Structures stained for pSer129 were subjected to high-resolution scanning with a pixel size of 6.25 µm. Subsequently, the images were imported into ImageJ, converted to grayscale, and the signal intensity was adjusted to a standardized threshold of 95% (consistent thresholds, exposure settings, and laser intensities were applied across all scans). Following this step, the “Process – Binary – Convert to Mask” function was employed. The mask function was determined using the default method against a black background. Next, a watershed filter was applied to select clusters with similar distribution. Finally, the “Analyze Particles” function was executed with a size range of 0 to infinity, displaying and summarizing the results. This facilitated the extraction of all clusters, including their coordinates and areas.

The subsequent step involved extracting clusters with a size equal to or smaller than 1 pixel (6.25 µm²) and clusters identified as artifacts (>15000 µm²), yielding the total number of clusters. Utilizing the Excel COUNTIF function, the number of small clusters (6.25 to 50 pixels), medium-sized clusters (50 to 200 pixels), and large clusters (>200 pixels) were calculated as proxies for the size of pSer129 aggregates.

Following the extraction of tissue surface and cluster count data, their relative density was determined for further analyses. Automated ImageJ macros were employed for the same procedures across all tissues (peripheral and brain) and animals, ensuring consistency in the analytical process.

### Serum analyses

Plasma Orexin A was measured by ELISA (Novus Biologicals, NBP2-80231, MO, US) according to manufacturer instructions. See dx.doi.org/10.17504/protocols.io.kxygx365zg8j/v1.

Plasma Melatonin was measured by ELISA Kit (Enzo Life Sciences, ENZ-KIT150-0001, NY, US) according to manufacturer instructions. See dx.doi.org/10.17504/protocols.io.81wgbx35ylpk/v1. Total and PS129 αSyn levels were measured using western blotting (WB) as described previously See dx.doi.org/10.17504/protocols.io.n92ldmk78l5b/v1.

### Microbiota analyses

#### DNA extraction

DNA extraction was carried out using the QiAamp PowerFecal Pro DNA kit (Qiagen, Sweden) and following the recommended protocol.

Amplicon, Quality control and OT production were carried out by Novogene see supplementary material and methods.

#### Alpha-Diversity

Alpha diversity is applied in analyzing complexity of species diversity for a sample through 6 indices, including Observed-species, Chao1, Shannon, Simpson, ACE, Good-coverage. Rank abundance curve Rank abundance curve can reflect the richness and evenness of species with samples through observing the width and shape of the curves.

#### Beta-Diversity

Beta diversity analysis was used to evaluate differences of samples in species complexity, Beta diversity on both weighted and unweighted unifrac were calculated by QIIMEsoftware (Version 1.9.1).

#### Community difference analyses

A series of statistical analyses, which include Anosim, Adonis, Multi-response permutation procedure (MRPP), Simper, T-test, MetaStat and LEfSe, were performed to reveal the community structure differentiation.

### Behavior testing

#### Olfactory discrimination test

Animals were transferred to a 30 x 30 cm grey open field chamber equipped with overhead housing lights and a camera recording system positioned above the apparatus.

On the first day of the experiment, mice were introduced to the apparatus for a duration of 20 minutes, accompanied by a 5 cm plexiglass circle cage filled with chocolate-flavored pellets (05684, Bio-Serv). On the subsequent day, a different apparatus was utilized, and the chocolate-flavored pellets were replaced with cotton pads infused with 1 ml of a 1% solution of 2,3,5-trimethyl-3-thiazoline (TMT, Sigma Aldrich) in water. See dx.doi.org/10.17504/protocols.io.36wgq3wwxlk5/v1.

#### Sucrose preference test

Individually housed mice in single cages were provided access to two novel bottles, one bottle contained a 1% sucrose solution, while the other contained regular water. See dx.doi.org/10.17504/protocols.io.4r3l22wwxl1y/v1.

#### Dark light box

A 45 x 20 x 20cm custom box divided into 1/3 dark (covered with lid, <5lux) and 2/3 white (∼100lux) was used for the experiment. Animals were released in the dark compartment before positioning the lid and left to explore for 6 minutes. See: dx.doi.org/10.17504/protocols.io.ewov19jz7lr2/v1.

#### Rotarod 40 RPM

The animals were positioned on an apparatus (Ugo Basile, cat # 47650, https://ugobasile.com/products/categories/motory-coordination/rotarod-for-mice-and-rats) with the receptacle platform in the upward position. Upon commencing the protocol, the speed was increased to 40 rotations per minute (40 RPM) and maintained at this speed for 120 seconds. See dx.doi.org/10.17504/protocols.io.3byl4qo5zvo5/v2.

#### Rotarod 20 RPM – docking

The animals were positioned on an apparatus (Ugo Basile, cat # 47650, https://ugobasile.com/products/categories/motory-coordination/rotarod-for-mice-and-rats) with the receptacle platform in the upward position. Upon commencing the protocol, the speed was increased to 20 rotations per minute (20 RPM) and maintained at this speed for five rotations before stopping. Subsequently, the platform rotated in the opposite direction for five rotations at 20 RPM (docking). See dx.doi.org/10.17504/protocols.io.3byl4qo5zvo5/v2.

#### Bedding test

Mice were individually housed for a minimum of 48 hours before testing. On the test day at 09:00, the mice were transferred to a clean cage with ad libitum access to food and water, featuring a 5×5 cm cotton pad positioned on the floor. After 24 and 48 hours, a picture was taken of the apparatus, and a bedding quality score (on a scale of 0 to 5 points) was assigned based on specific parameters: 0) did not touch the pad, 1) unfolded the pad, 2) folded the pad in a corner, 3) folded the pad in a corner with bite marks, 4) initiated shredding the pad and created a nest, and 5) the pad is shredded, and a proper nest is visible. See dx.doi.org/10.17504/protocols.io.n2bvj3ko5lk5/v1.

#### Descending pole test

Mice were positioned on top of a wooden pole (50 cm long, ∼1 cm diameter) leading to a clean cage.

#### Descending inclined platform test

At the initiation of the test, mice were positioned on the horizontal section of a 60 cm long, 45° inclined platform featuring a 1 cm grid. Subsequently, mice were gently prompted to the starting point of the inclined path and allowed to move down freely. The time taken to reach the clean cage positioned at the bottom was then measured utilizing video recorded sessions. See DOI: dx.doi.org/10.17504/protocols.io.14egn3ko6l5d/v2.

#### Operant behavior

Operant behavior testing was conducted in individually housed mice without access to regular food. In each animal cage, a FED3.0 operant box was positioned (https://open-ephys.org/fed3/fed3, Matikainen-Ankney *et al.*, 2019).

Initially, mice were exposed to the FED3.0 for 24 hours with 6g of chow pellets placed on the floor. In the following two days, mice were exposed to free-feeding sessions where 20mg sugar pellets were randomly delivered to habituate the mice. The next three days involved a switch to a Fixed Ratio 1 (FR1) protocol starting at 09:00. Following three consecutive sessions of FR1, the protocol was increased to Fixed Ratio 3 (FR3) for one day. Finally, on the last day, the protocol was switched to a “Follow the Light” protocol for 24 hours. See: dx.doi.org/10.17504/protocols.io.j8nlk8b31l5r/v1.

#### Telemetry implantation

An electroencephalogram (EEG) electrode (Agnhtos.se, MCS1×2) was then screwed into the skull above the frontal cortex (AP: 1.5, ML: 1.0), and a reference electrode was secured into the skull above the cerebellum. Electromyogram (EMG) electrodes were positioned in the nuchal muscles on the same side with approximately 1mm spacing; these electrodes were then affixed using small heat-shrink tubing. Subsequently, a sizable chamber was created in the abdominal area to accommodate the wireless telemetry recording device (F20-EET, DSI, USA). See : https://www.protocols.io/view/procedure-for-eeg-surgery-kxygx3dxog8j/v1

#### Telemetry recording

Following a recovery period of at least 1 week post-surgery, animals were transferred to the sleep-recording room and allowed to acclimate for a minimum of 48 hours. During the recordings, animals were group-housed with 1-2 cage mates throughout the entire procedure, enabling the characterization of sleep independently of any social stress factors. On the day of the recording, the telemetry device was activated using a strong magnet, and the cages were placed on the telemetry receiver (DSI, https://www.datasci.com/products/software/neuroscore). A total of 16 animals were recorded simultaneously.

EEG, EMG, and movement activity were recorded via telemetry using Neuroscore v3.0 software (DSI) at a frequency of 500Hz for EEG/EMG and 1Hz for activity. Mice were recorded for 24 hours without interruption in a soundproof room, with ad libitum access to food and water.

#### Sleep data analysis

Sleep data files were imported into Neuroscore V3.0 software and visually inspected.

The contributing factors of Delta, Theta/Delta, and EMG were considered equally important for all recordings. Sleep scoring was then obtained within 20s windows, and any recording with more than 1% artifacts was excluded from further analyses. See dx.doi.org/10.17504/protocols.io.yxmvm3rrbl3p/v1.

#### REM sleep behavior disorder scoring and Periodogram analyses

Following the scoring of sleep stages, EMG was converted into the root mean square (RMS) using the signal grid function. For all recordings, a signal grid was generated, including the timestamps (20s windows) of the GMT time, the RMS of the EMG, the sleep scoring, and the EEG. Additionally, a periodogram Power Band (PB) for each 20s window was obtained within the 0.3 to 80 Hz band range. The analysis employed a 10/1024 FFT order, an overlap of 50%, and a Hamming spectral function applied to the window. PB values were expressed as a relative percentage of the power band value. Delta oscillation was determined within the 0.5 to 4Hz window, Theta within the 4 to 8Hz window, Alpha within the 8 to 12Hz window, Sigma within the 12 to 16Hz window, Beta within the 16 to 24Hz window, Low Gamma within the 24 to 49 Hz window, and High Gamma within the 51 to 80Hz window. See dx.doi.org/10.17504/protocols.io.81wgbxjdolpk/v1.

#### Fiber Photometry during sleep

A subset of mice from each group received a combination of two Adenoviruses (AAV) at a ratio of 1:1 for imaging dopamine release (AAV9-hSyn-GRAB-rDA1h, 250nl, addgene #140557) or acetylcholine release (AAV1-CAG-iAChSnFr, 250nl, addgene #137955) in the dorsomedial striatum (coordinates from the Bregma: AP 0.5mm, ML 1.1, DV 3.0mm) in the same injection. After a 2-week interval, mice were implanted with a sleep recording telemetry device and an optic fiber (0.50NA, 400µm, 4mm long, Thorlabs) above the virus injection site. For each animal, a minimum of 30 minutes of data were acquired and repeated multiple times to cover multiple sleep cycles. Recordings were limited to 30 minutes to prevent the excitation light from affecting the mice’s sleep schedule and to minimize interference from the patch cord on their ability to sleep.

Following recording, sleep scoring was processed as described above, extracting timestamps of Wake, REM, NREM, and RBD events. Using a custom-made Matlab script, photometry data were downsampled to 500Hz (average smooth window function), then aligned with the sleep scoring. For each 1-minute window, each channel of the photometry was normalized using the z-score function and then averaged over the 20s window corresponding to the sleep scoring. See: dx.doi.org/10.17504/protocols.io.6qpvr8xw3lmk/v1.

#### Data exclusion

All data were tested in with GraphPad Prism software (www.graphpad.com, version 9, GraphPad (RRID:SCR_000306)) for Rout method for outliers with a Q value of 5%.

#### Data sharing

Detailed protocols are available on https://www.protocols.io platform. Prism file including all analyses are available on the https://zenodo.org platform DOI: 10.5281/zenodo.10822458. All raw data and scripts are shared under the ASAP sharing depository.

### Statistics analyses

Statistical analyses were conducted using GraphPad Prism software (version 9). For comparisons involving two groups, paired- or unpaired-t-test analyses were applied. In cases involving three or more groups, a 1-way ANOVA analysis with a Tukey post hoc analysis was performed. At the exception of serum analyses where LSD post hoc test were used as well as the test where ANOVA display significant P-values but Tukey post hoc test show no significant effects (always indicated in figure legends). In the analyses of gut-injected animals, post hoc analyses were provided for the comparison of monomeric αSyn-injected animals with the αSyn-PFF-injected group, with additional statistical values available in the attached Prism files. For comparison of the surgery effect (i.e monomeric αSyn 2 weeks versus 7 months) a 2-way ANOVA time * group was conducted. For comparisons of sex differences, a 2-way ANOVA sex×group analysis was conducted, and the interaction effect was reported in the figure legend. In anatomical analyses of ROIs, a 1-way ANOVA or 2-way ANOVA analysis was performed on each ROI and presented as imagesc – Jet, while post hoc analyses were conducted using Dunnett’s test by default. Outlier data were removed using the Prism Rout method with a 5% Q value.

Statistical significance was considered with a P-value less than 0.05, denoted as *, P < 0.05, **, P < 0.01, ***, P < 0.001, and ****, P < 0.0001.

## List of Supplementary Materials

Materials and Methods

Fig. S1 to S23

## Acknowledgments

We thank Dr. Anderson Camargo for his contribution to early data collection and Novogene for the analyses and data representation of microbiota. This research was funded by Aligning Science Across Parkinson’s [020608] through the Michael J. Fox Foundation for Parkinson’s Research (MJFF) (D.D., W.P., R.M, M.G.K, R.C, V.L.D, T.M.D., P.S.). Other funding includes the NIH/NIA R01AG085688 (V.L.D., T.M.D), the JPB Foundation (T.M.D.), the Robert J and Claire Pasarow Foundation (V.L.D., T.M.D.). T.M.D. is the Leonard and Madlyn Abramson Professor in Neurodegenerative Diseases.

## Author contributions

D.D and P.S conceptualized the project, D.D, G.M.S, P.W, and D.D.C, performed experiments. M.K., R.C., V.L.D. and T.M.D. provided reagents. D.D and P.W analyzed the data. D.D, X.Z, P.W, and PS wrote the original draft. All authors contributed to reviewing and editing the final version of the manuscript.

## Data and materials availability

For open access, the author has applied a CC-BY 4.0 public copyright license to the Author Accepted Manuscript (AAM) version arising from this submission.

---

**Figure S1.**
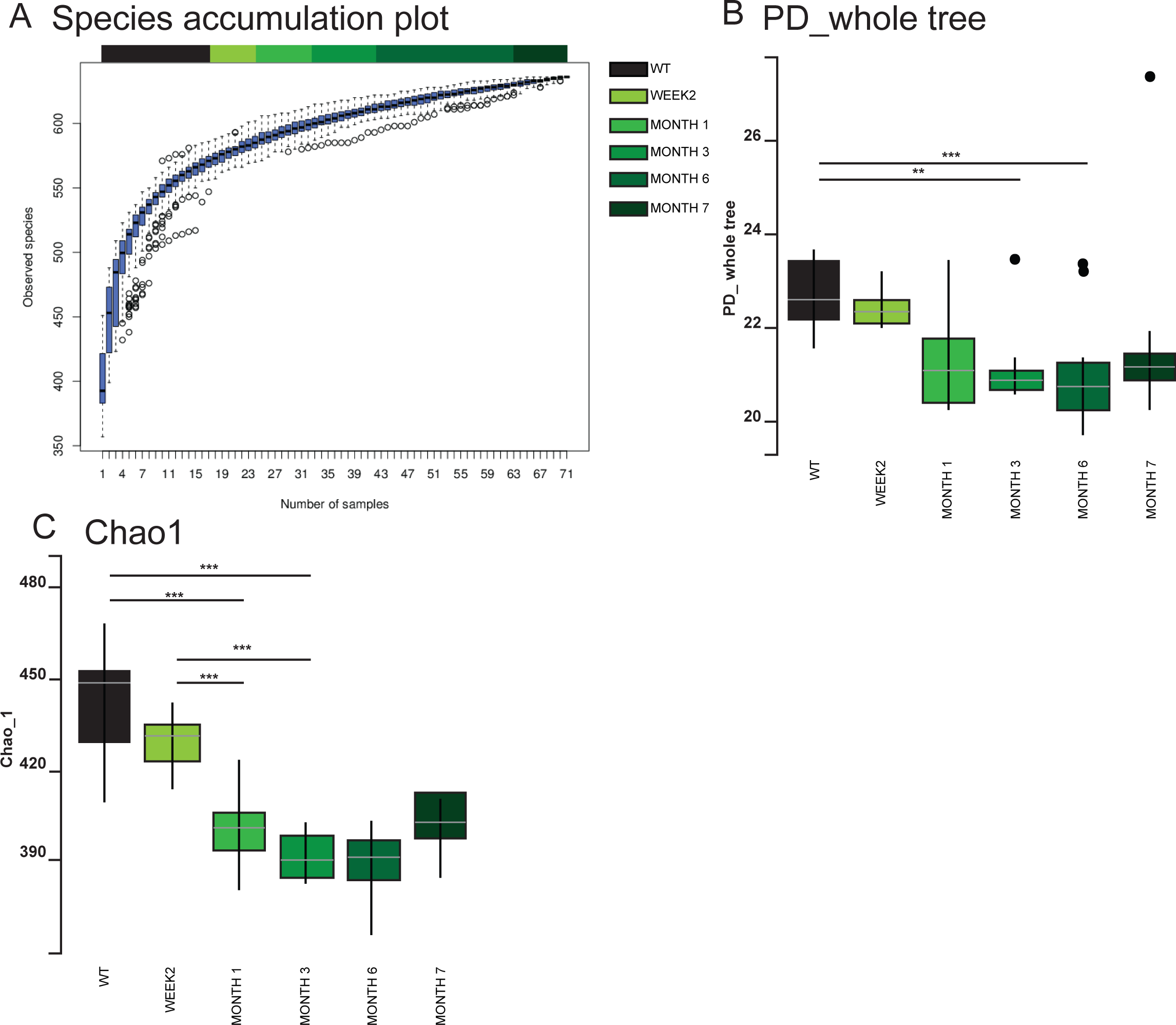

**Figure S2.**
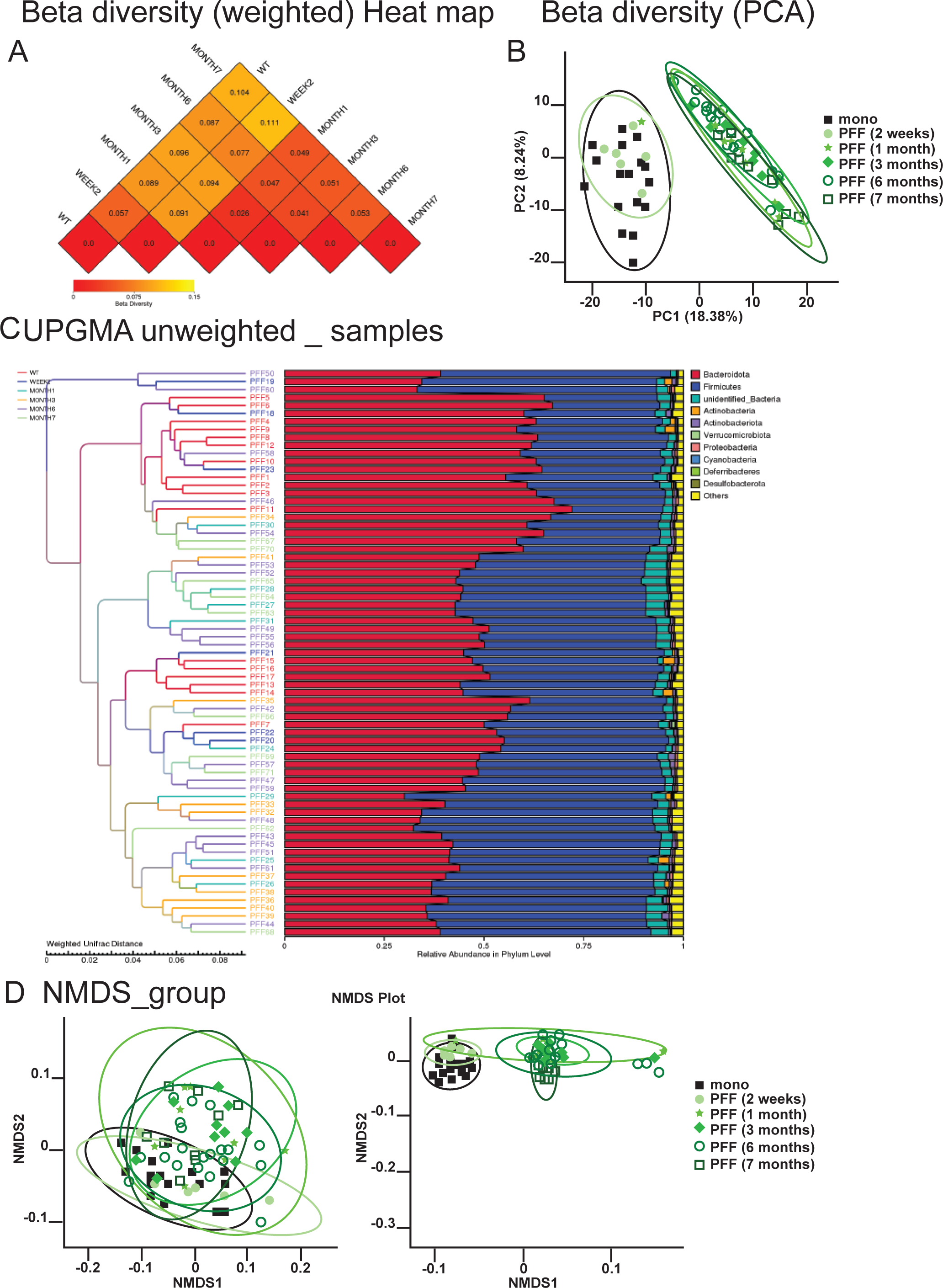

**Figure S3.**
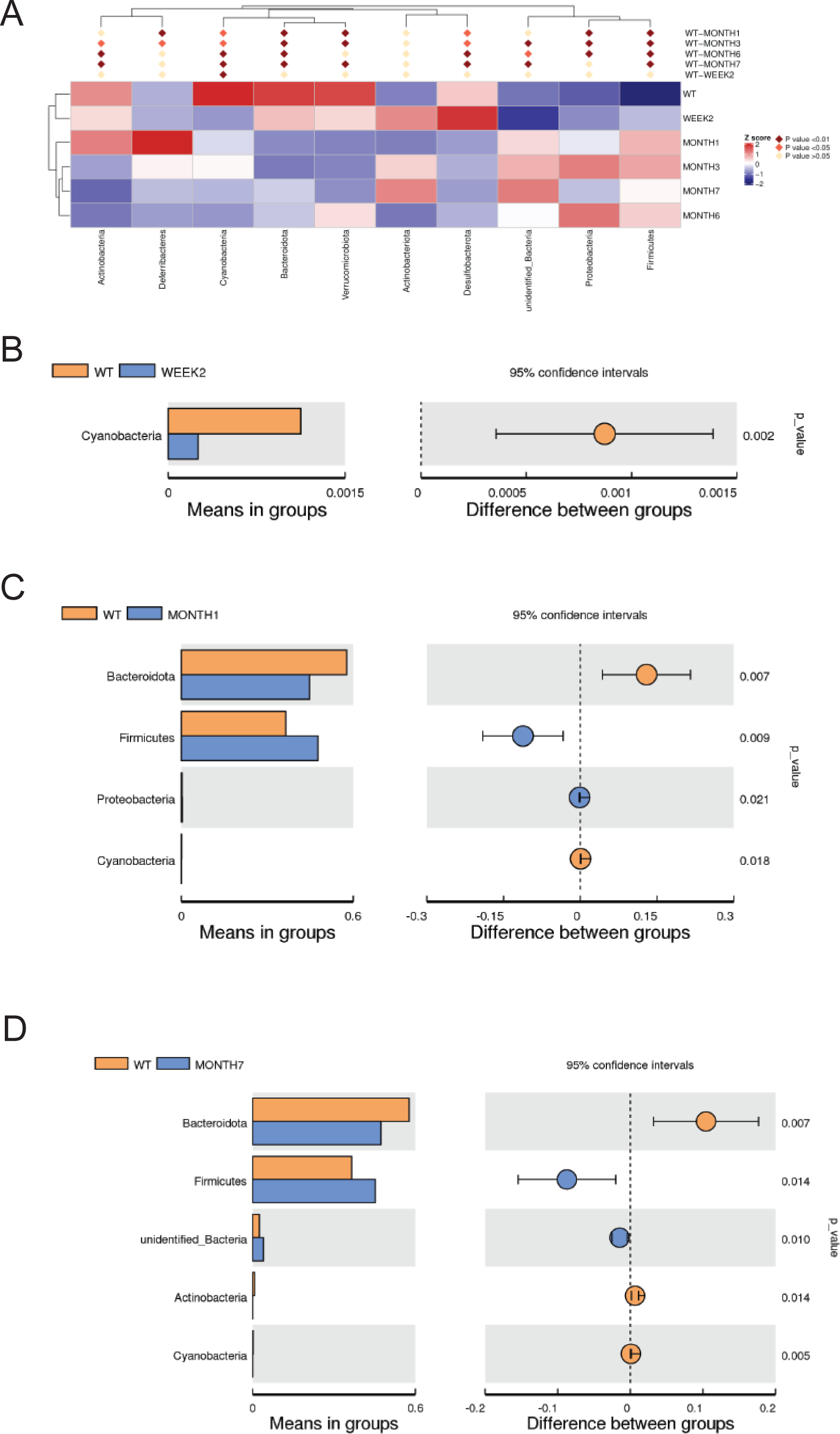

**Figure S4.**
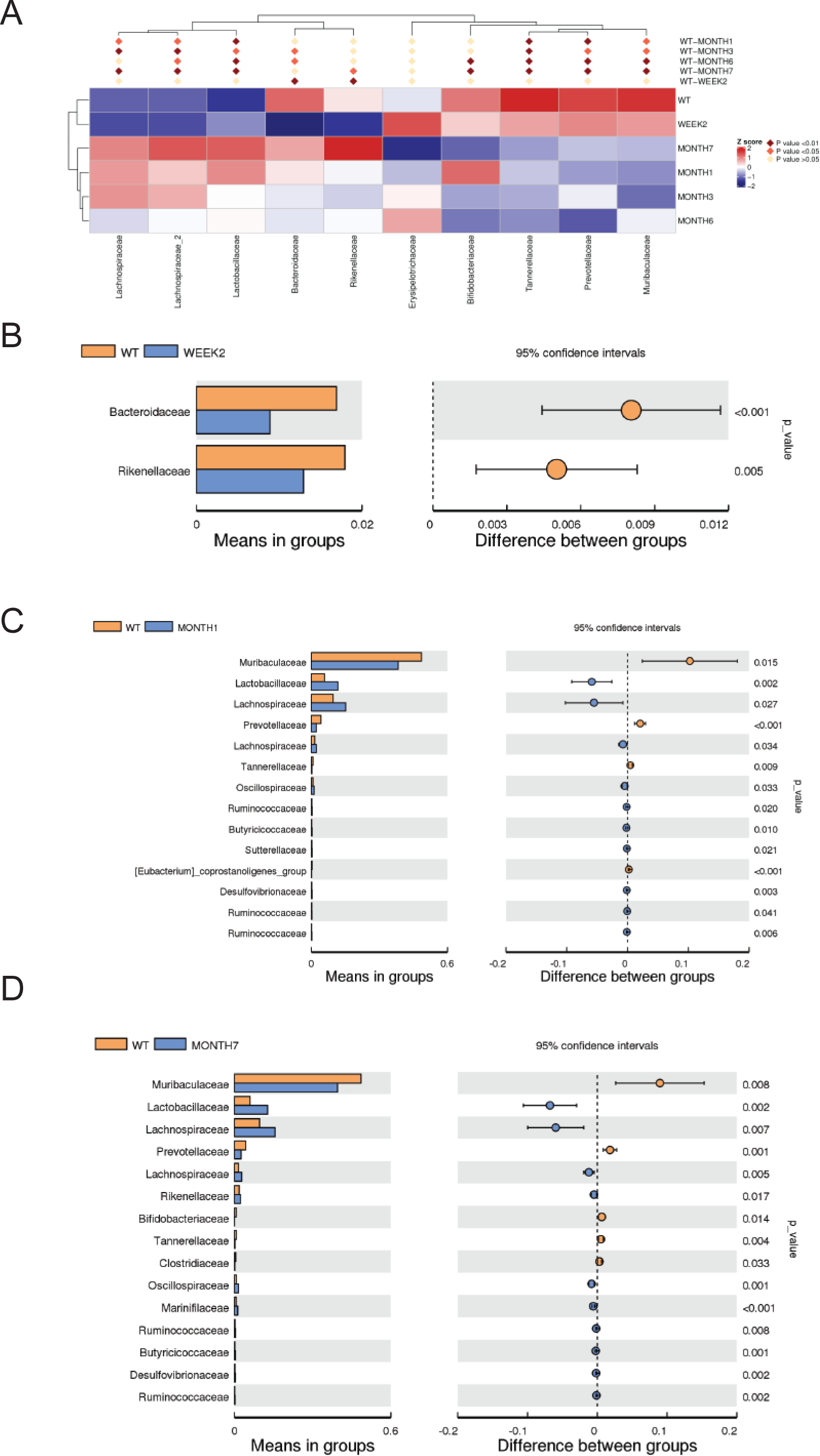

**Figure S5.**
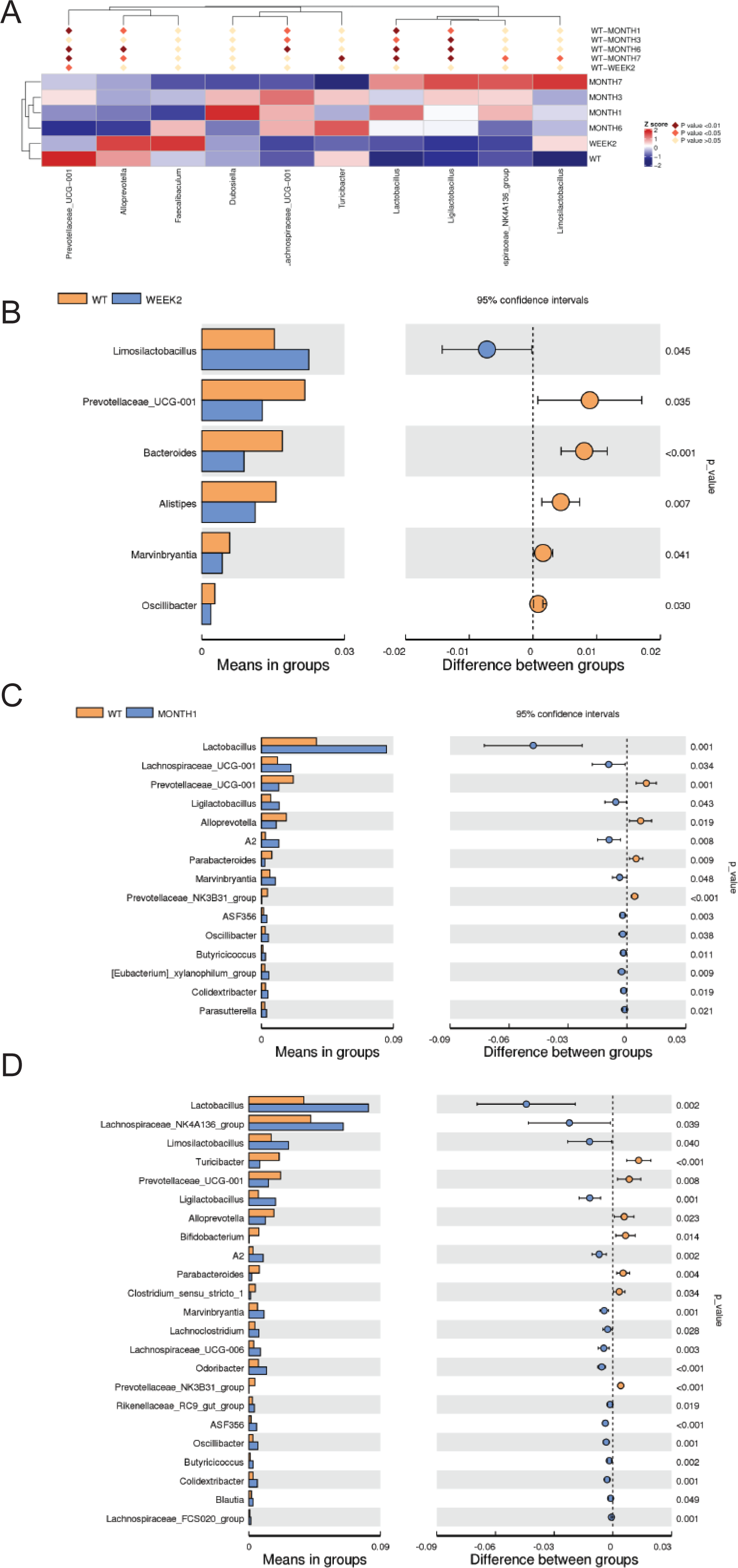

**Figure S6.**
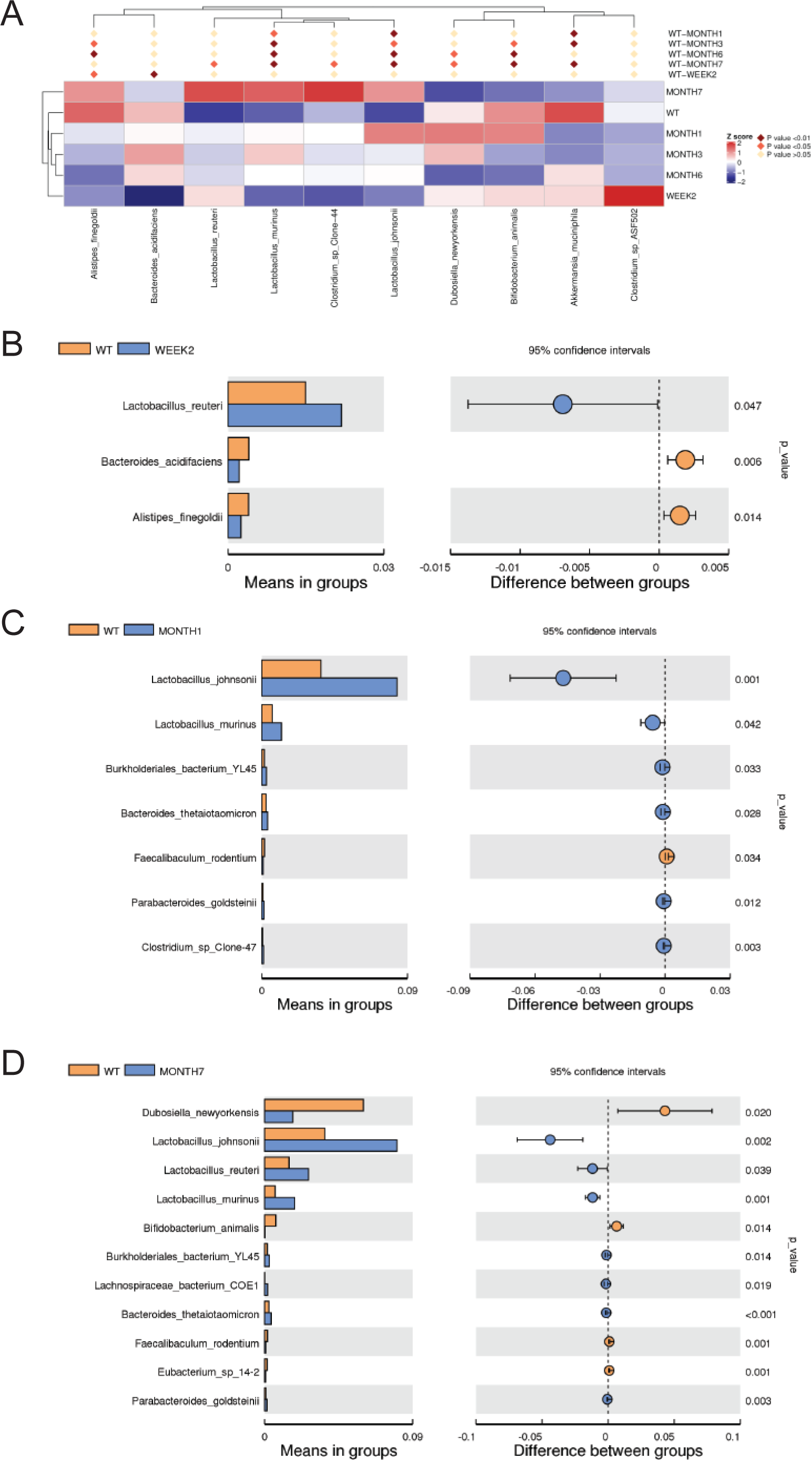

**Figure S7.**
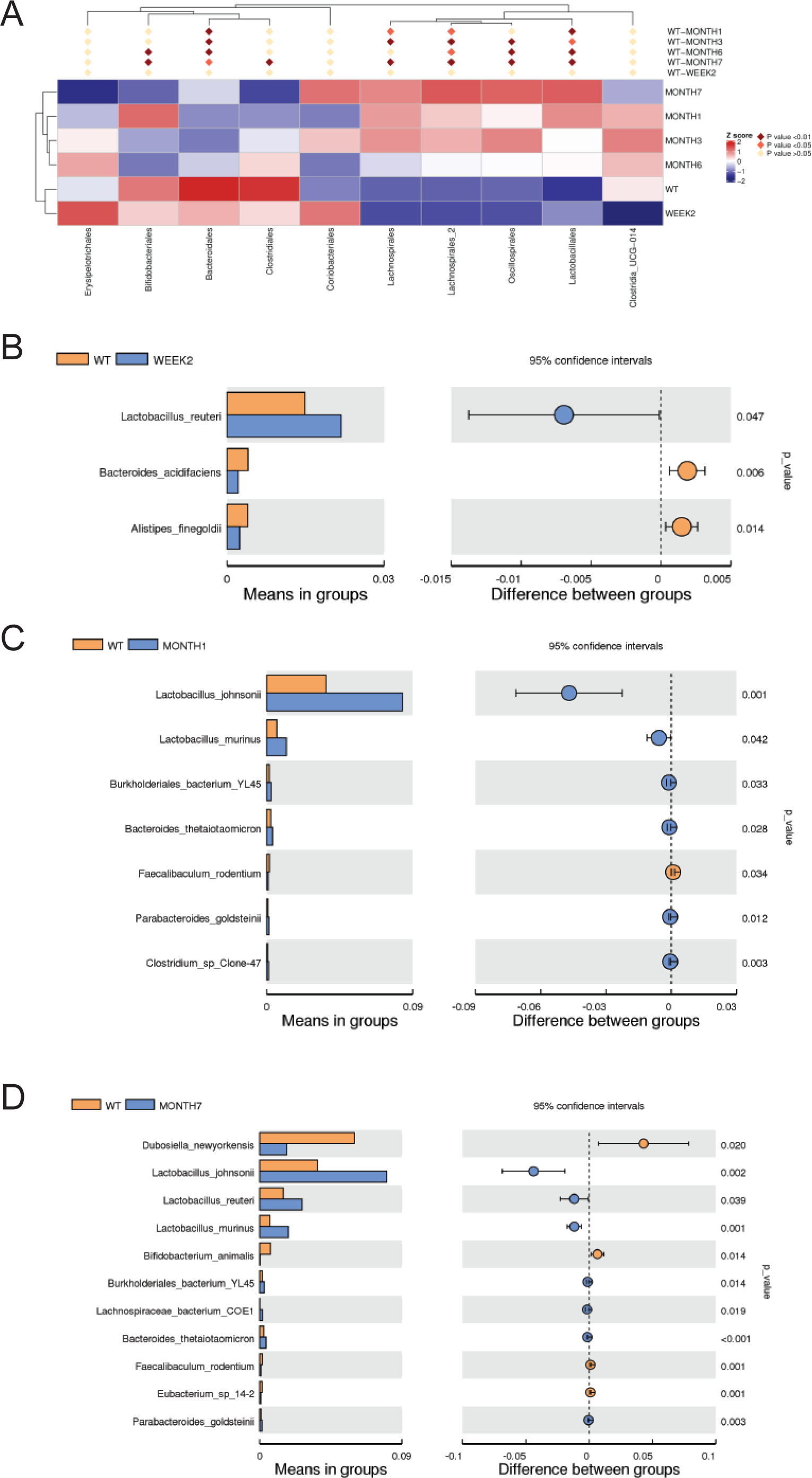

**Figure S8.**
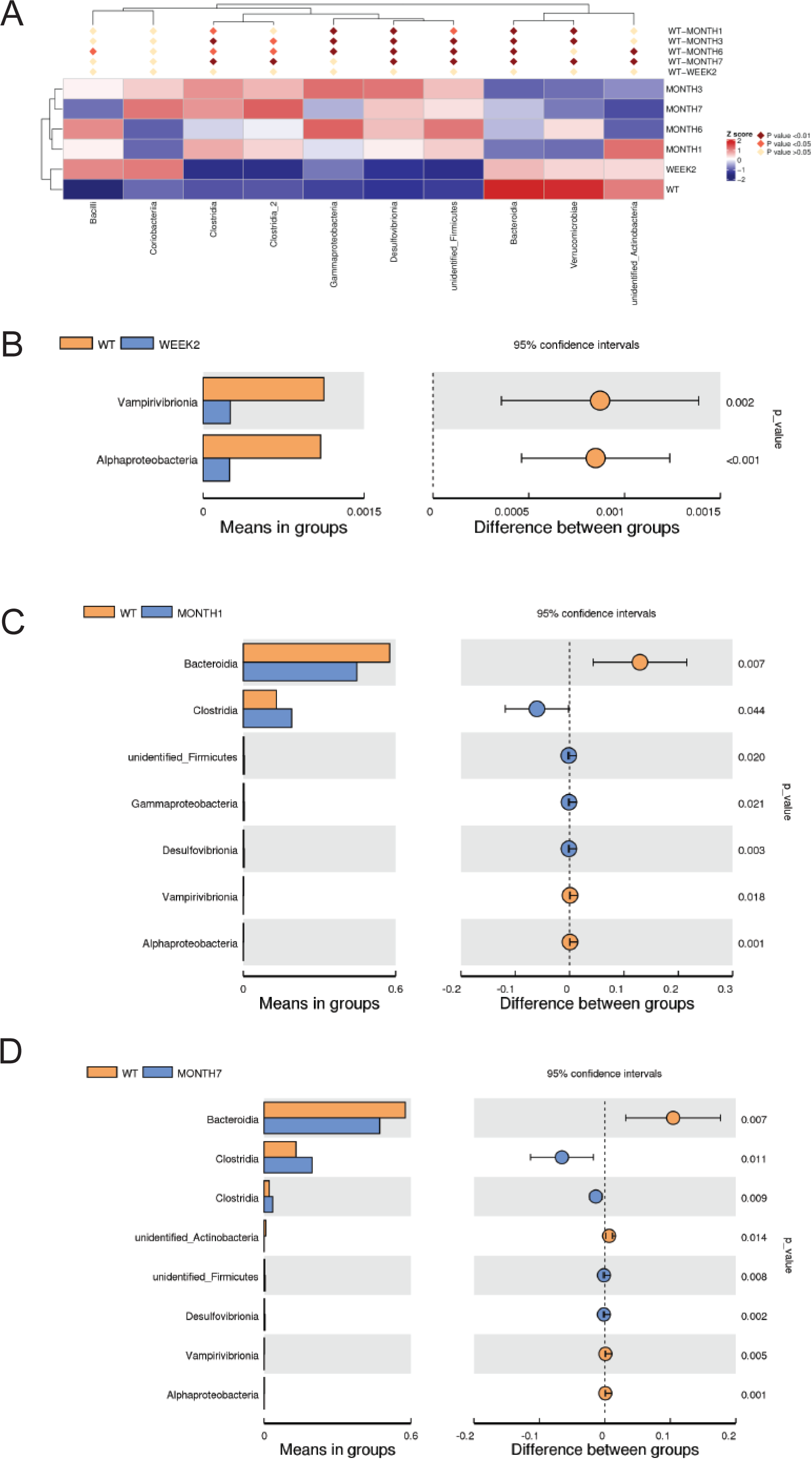

**Figure S9.**
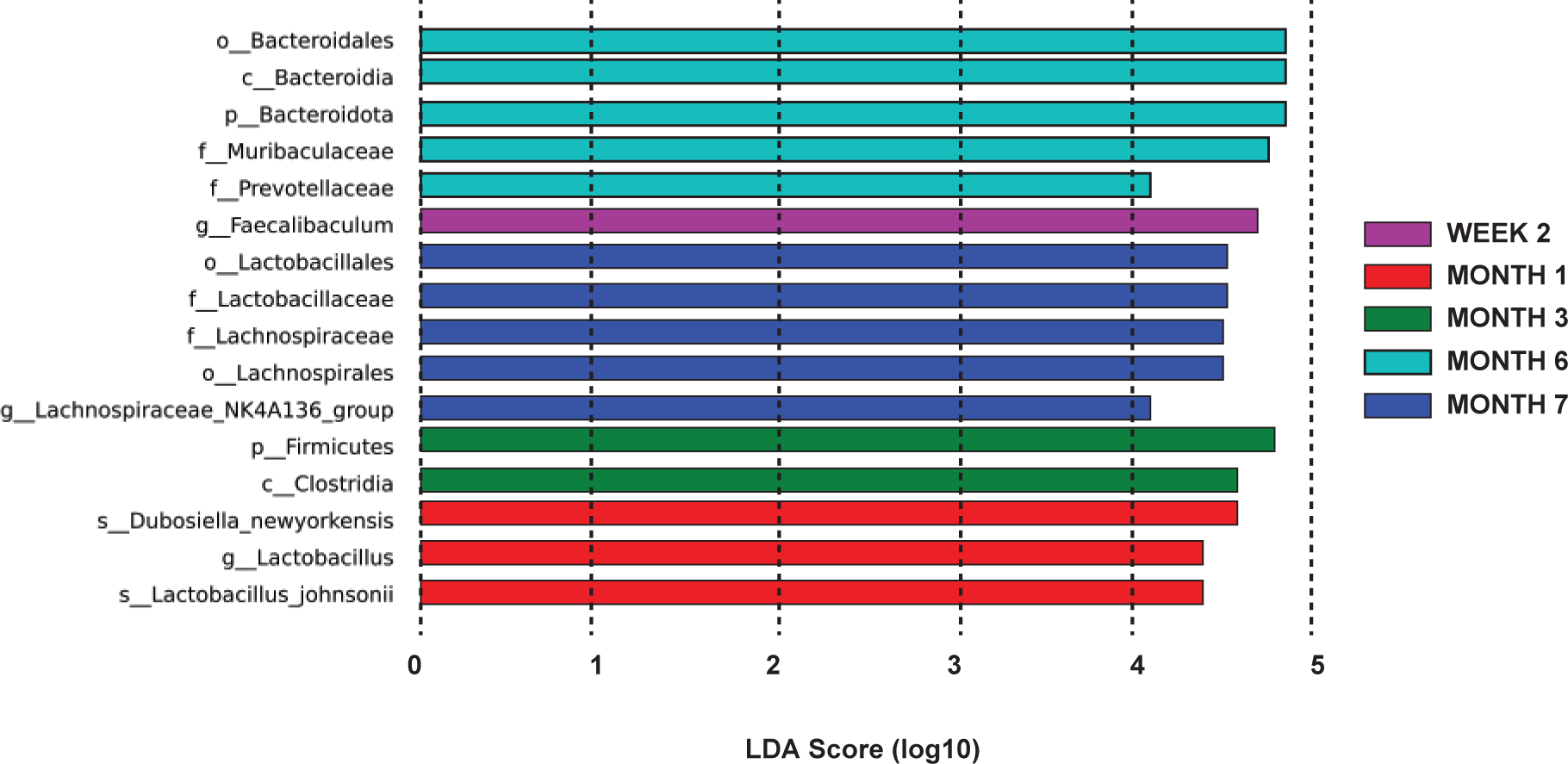

**Figure S10.**
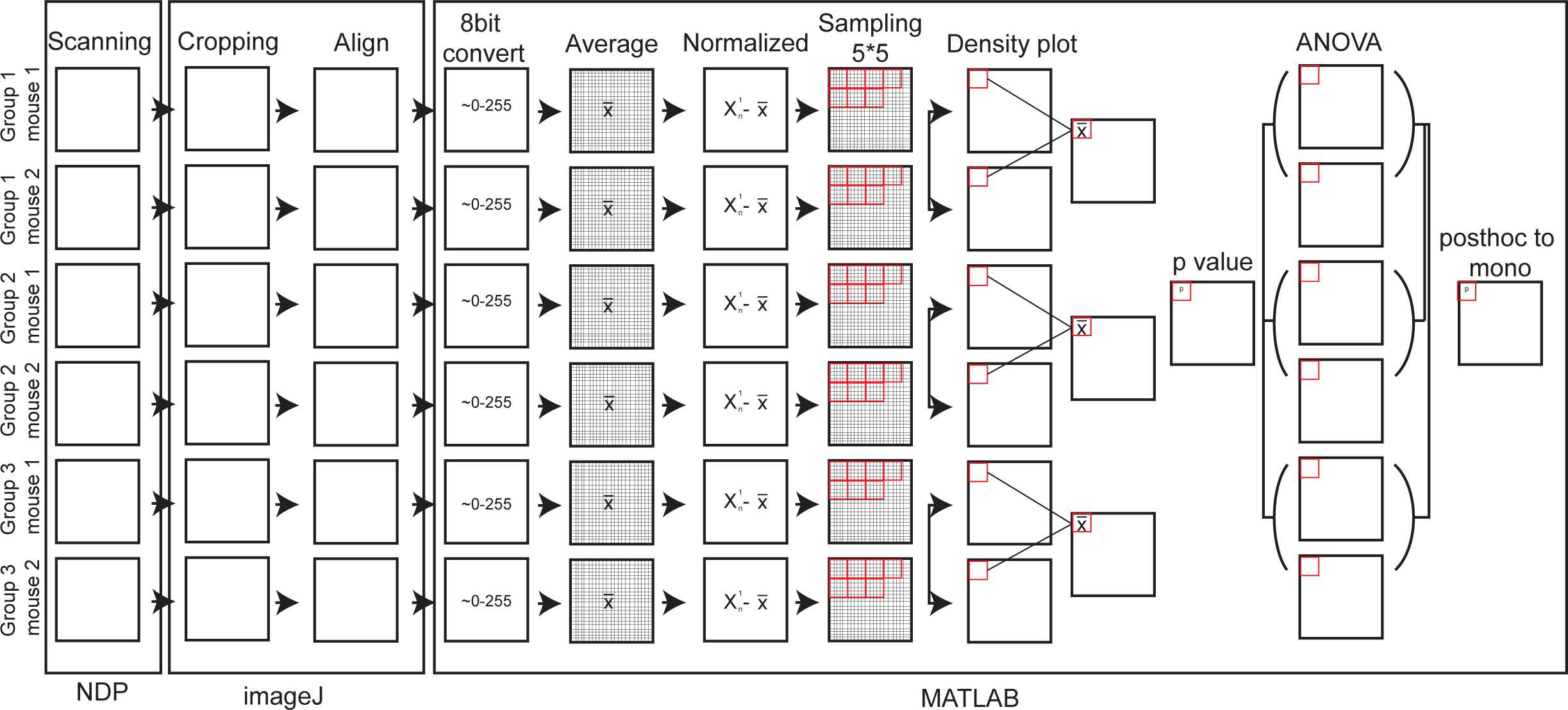

**Figure S11.**
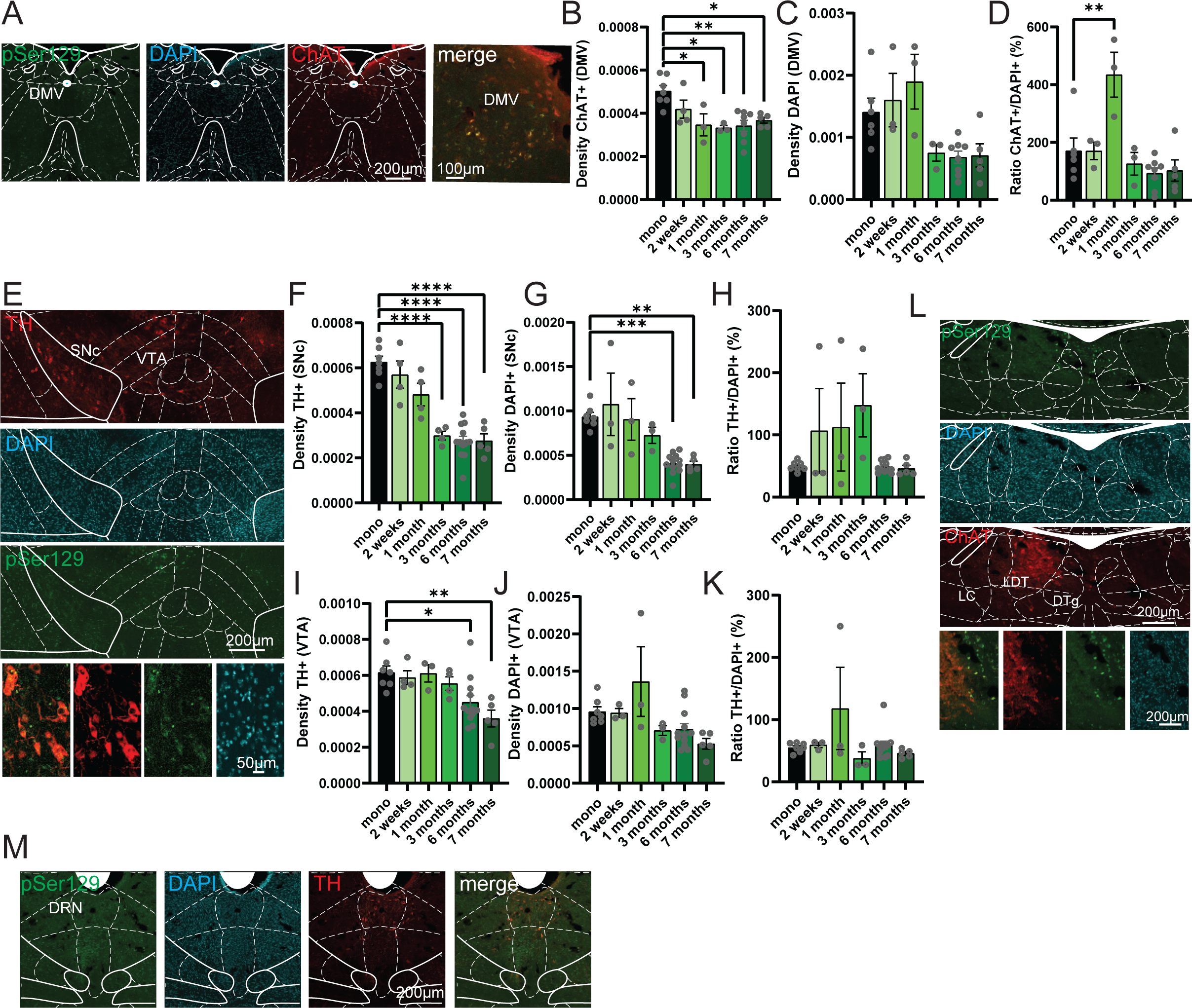

**Figure S12.**
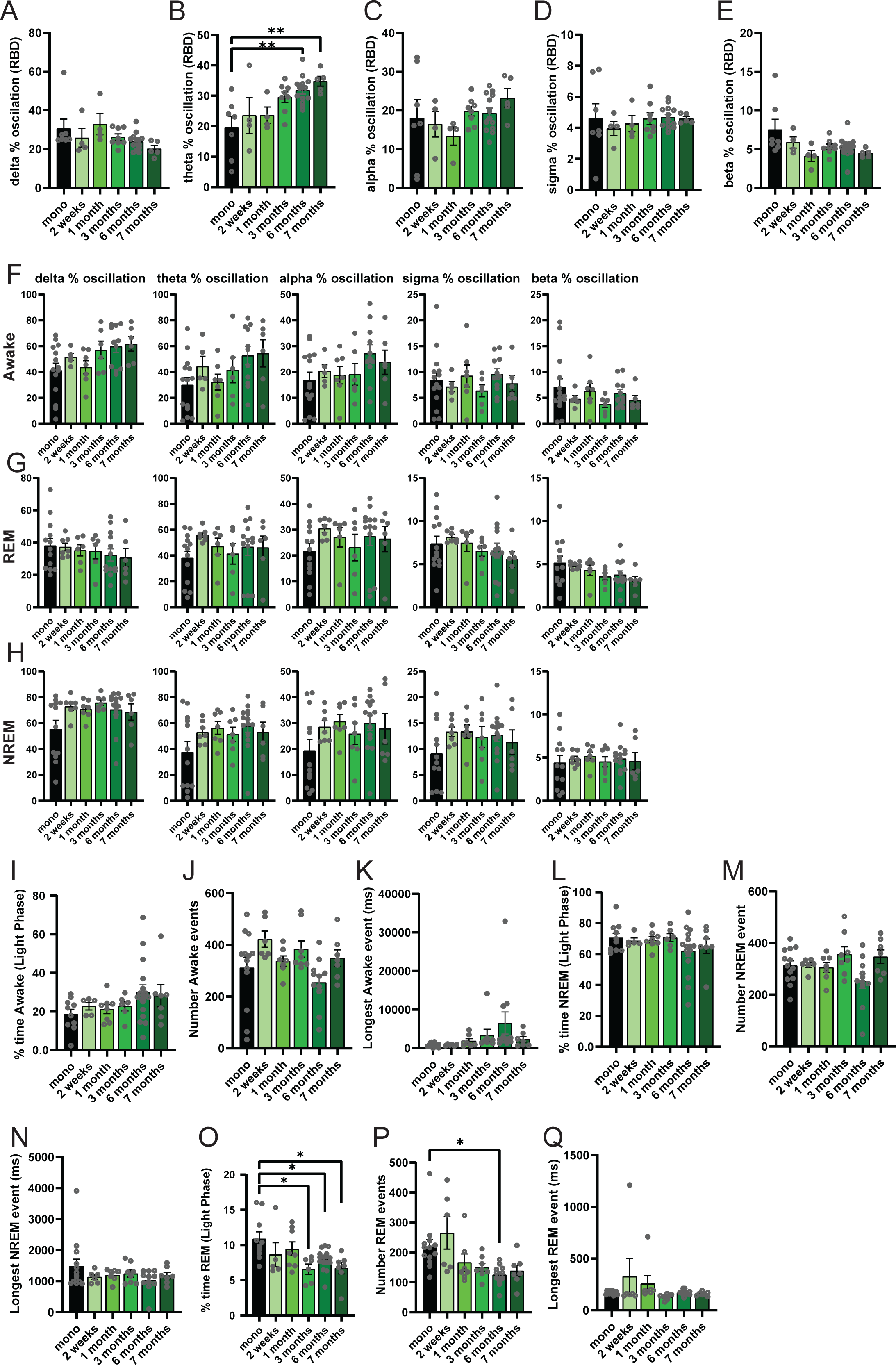

**Figure S13.**
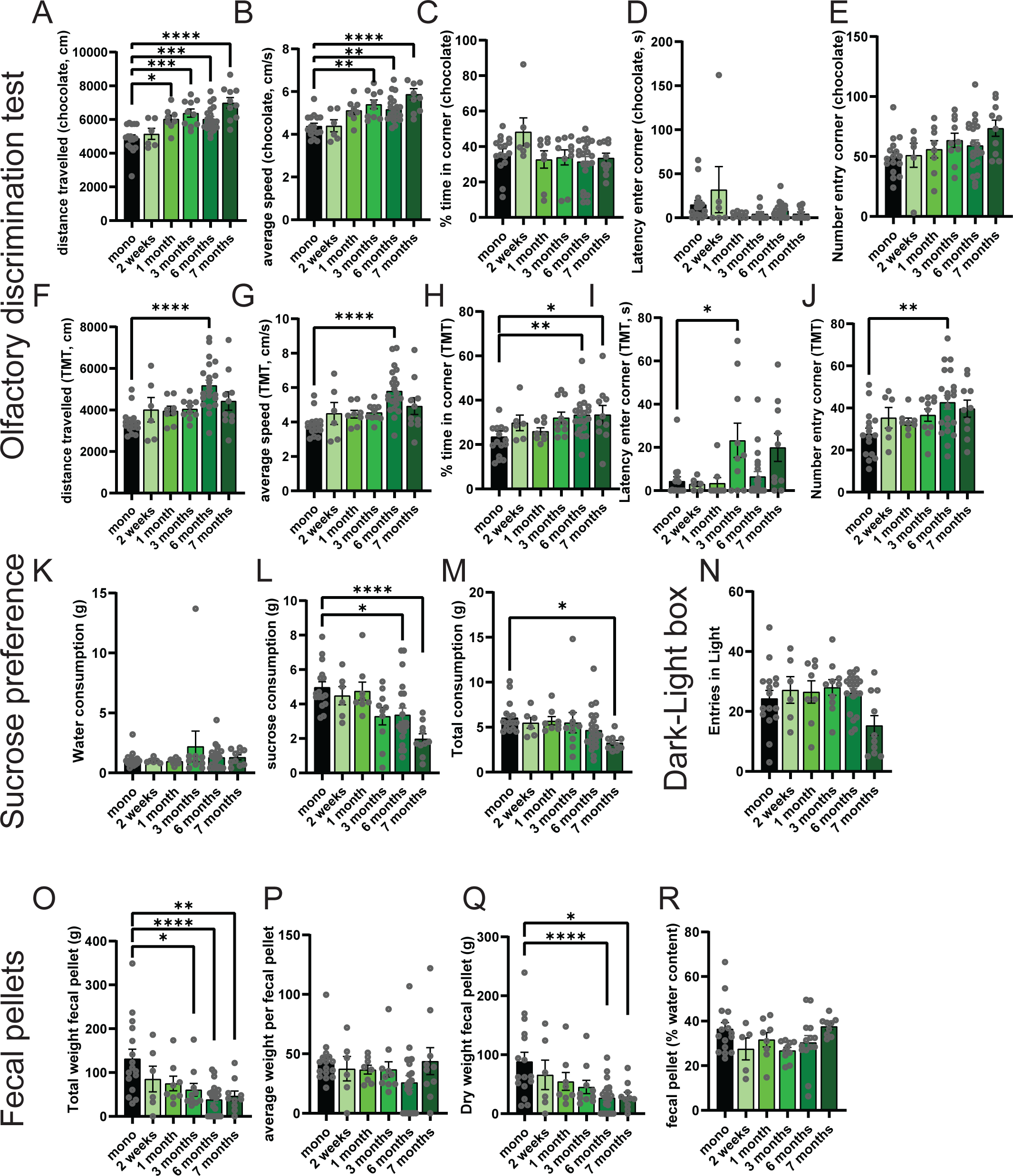

**Figure S14.**
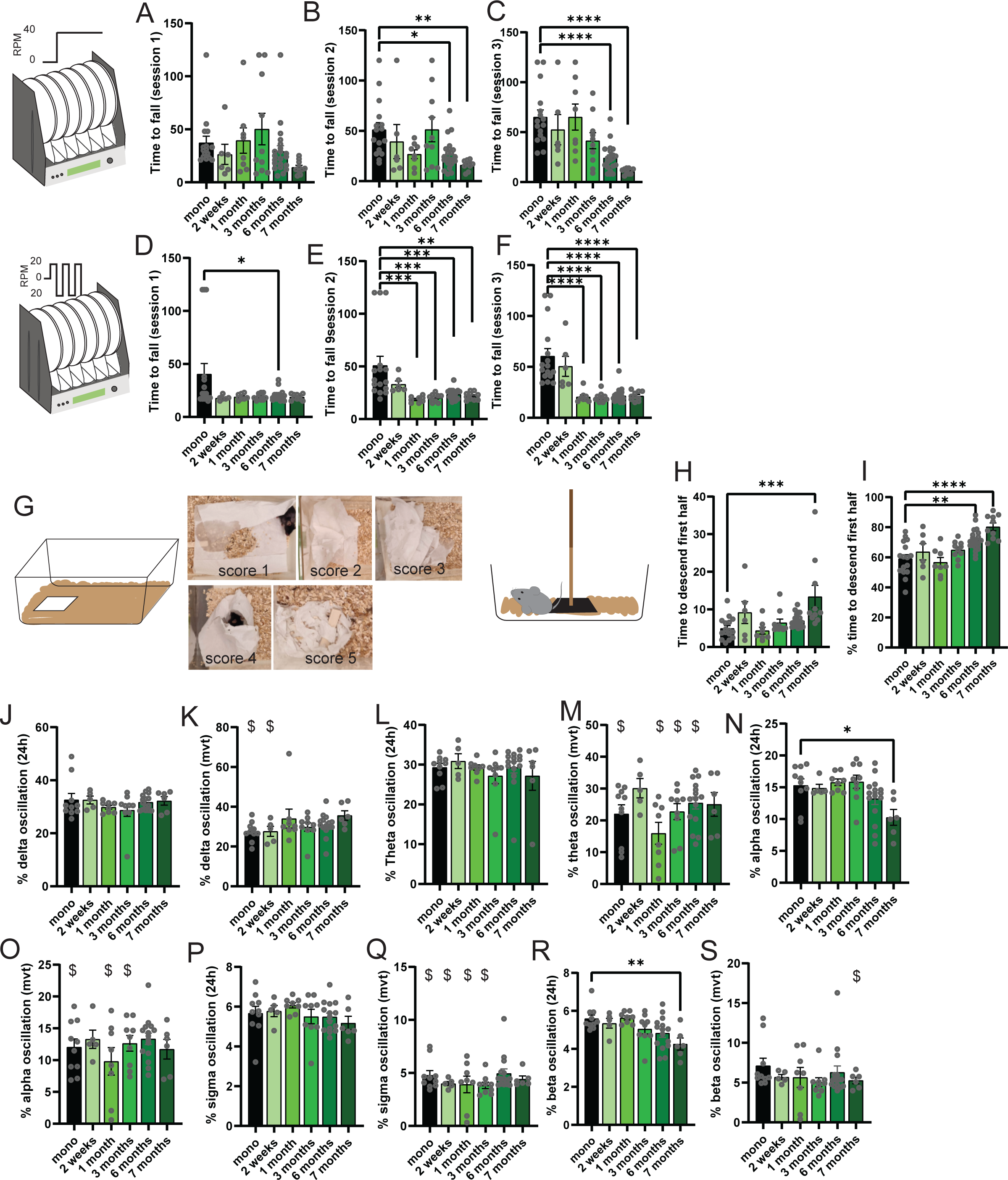

**Figure S15.**
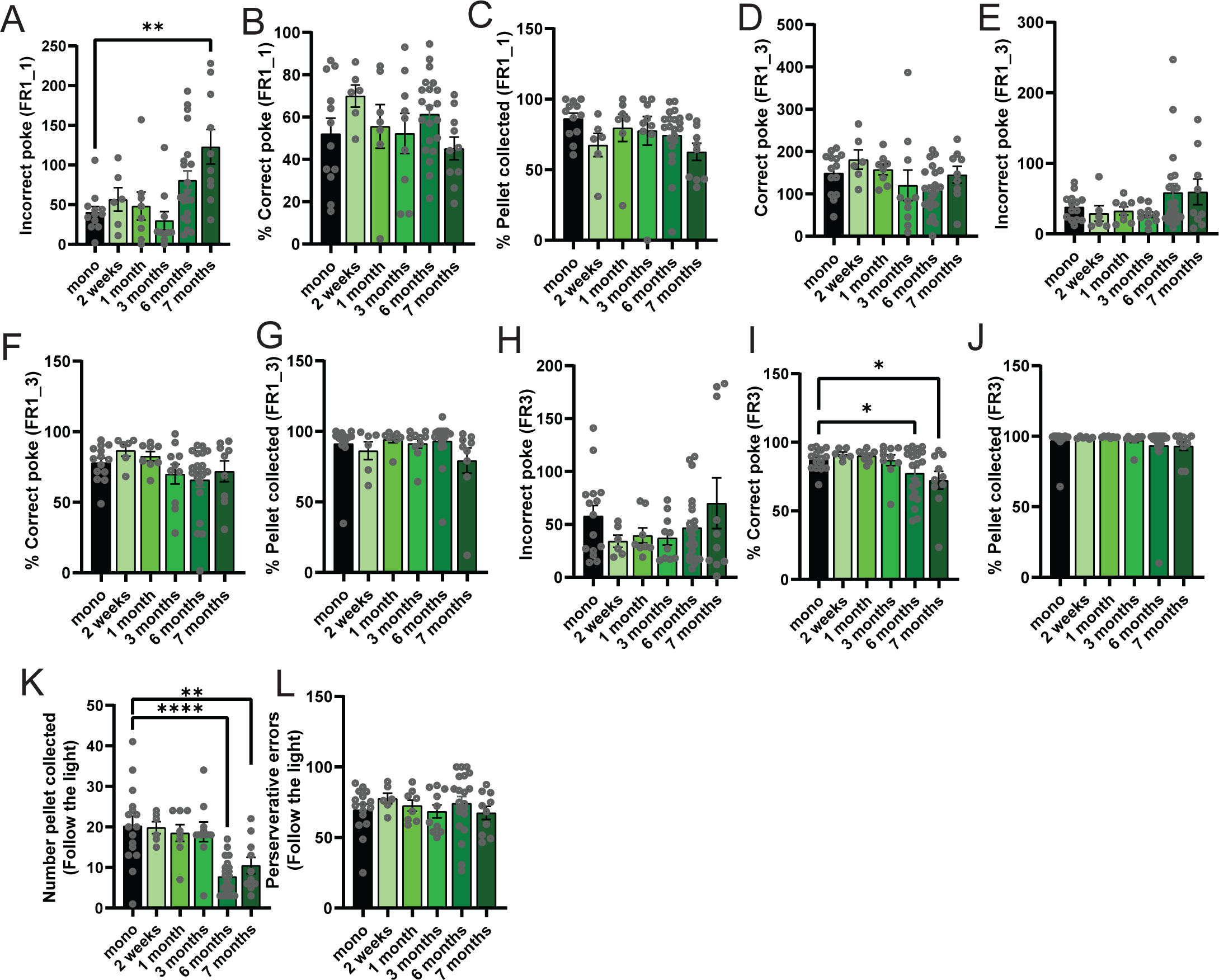

**Figure S16.**
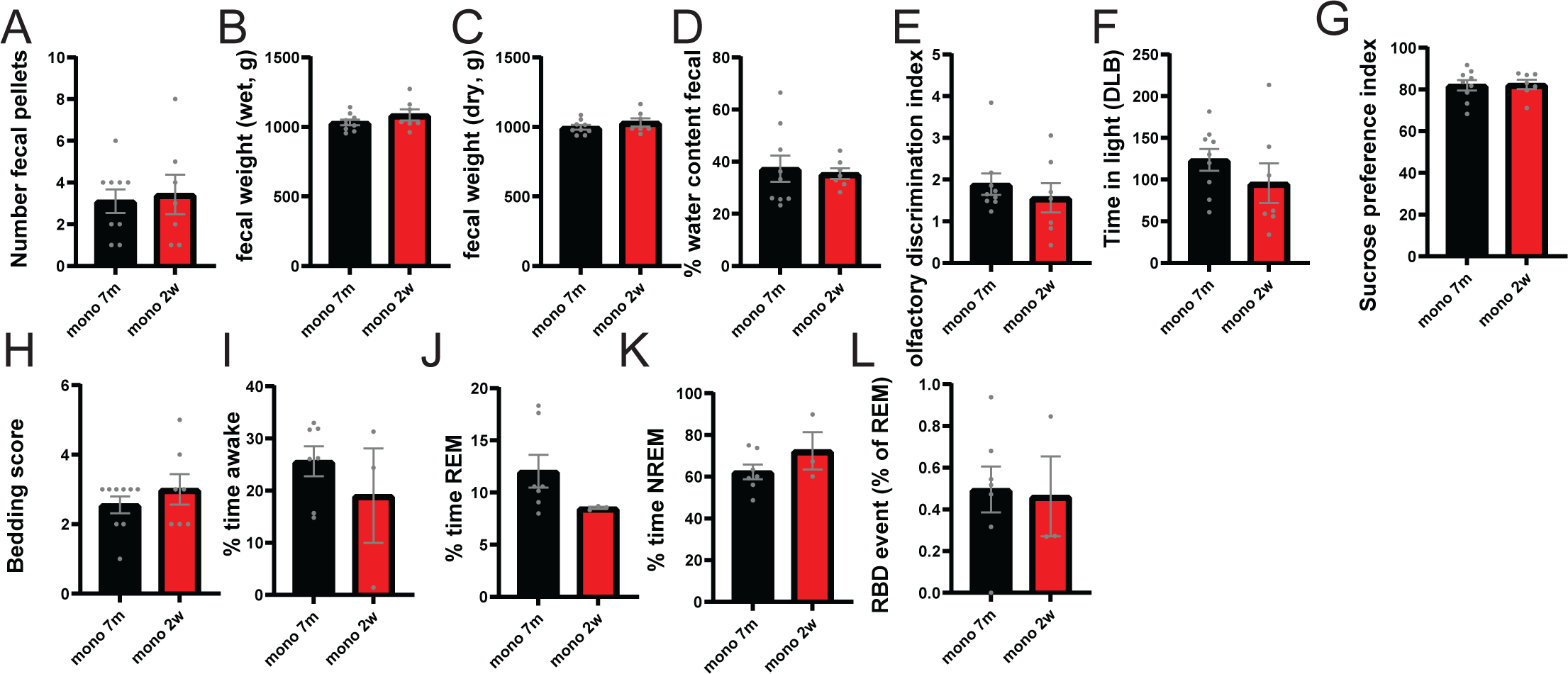

**Figure S17.**
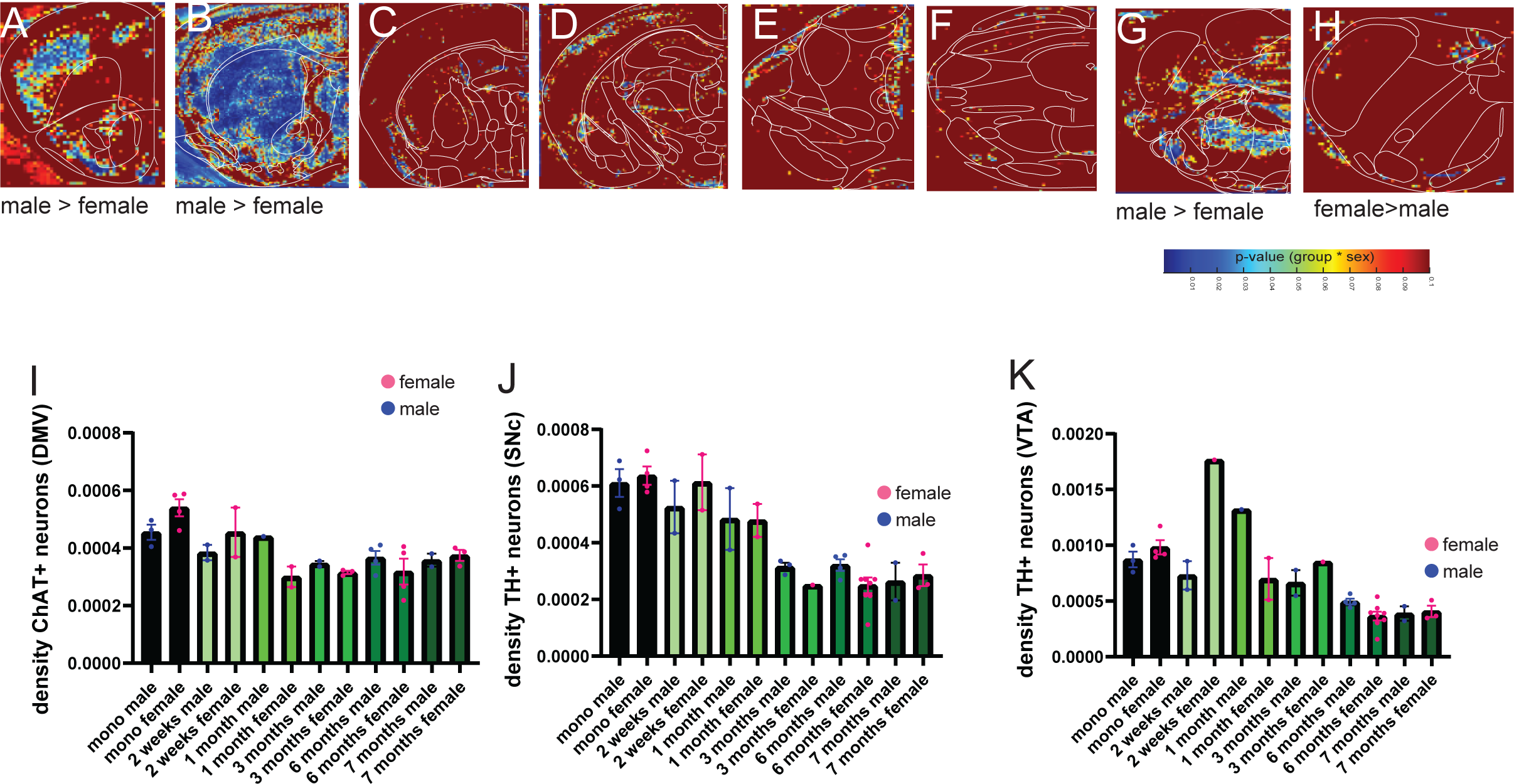

**Figure S18.**
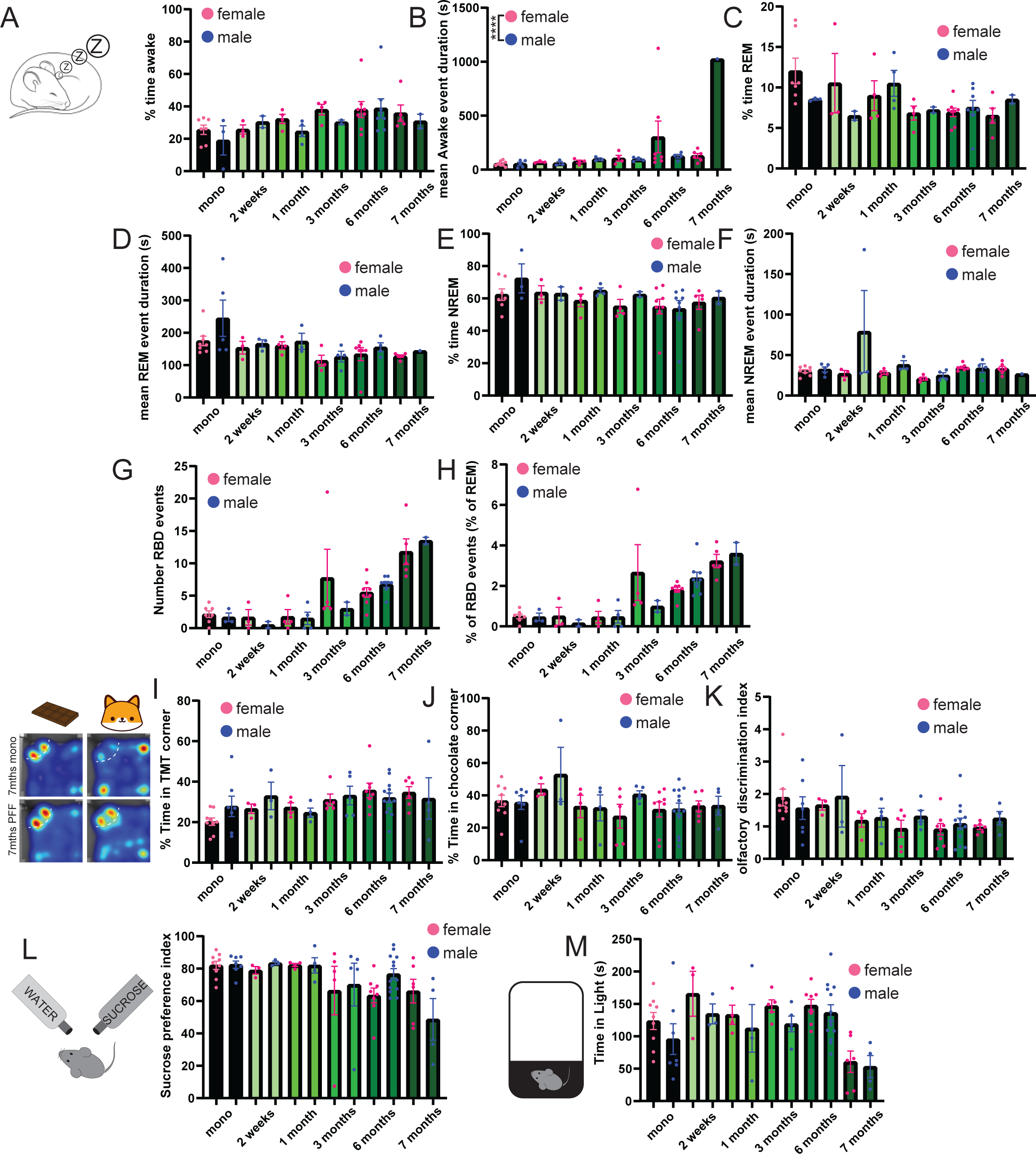

**Figure S19.**
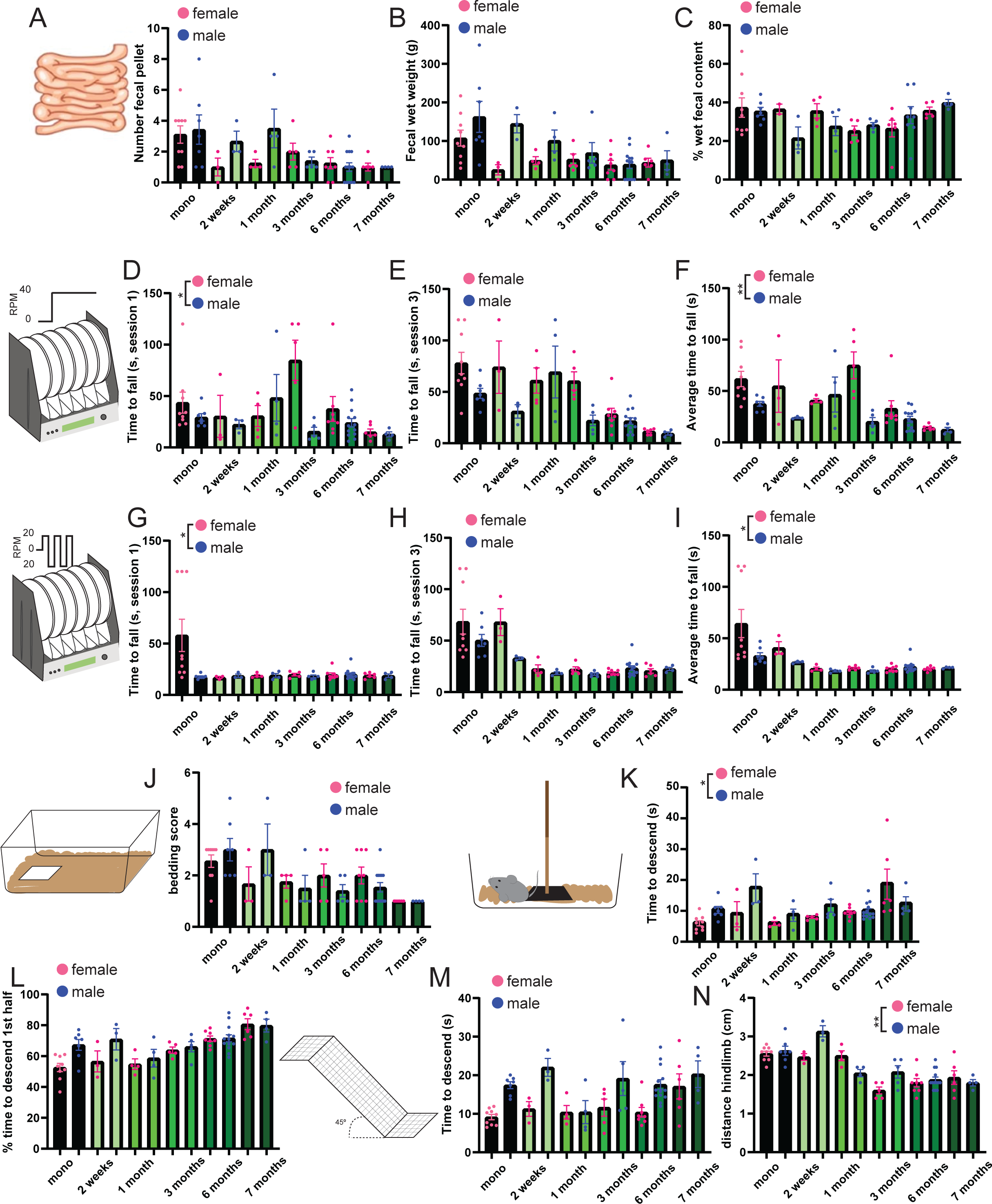

**Figure S20.**
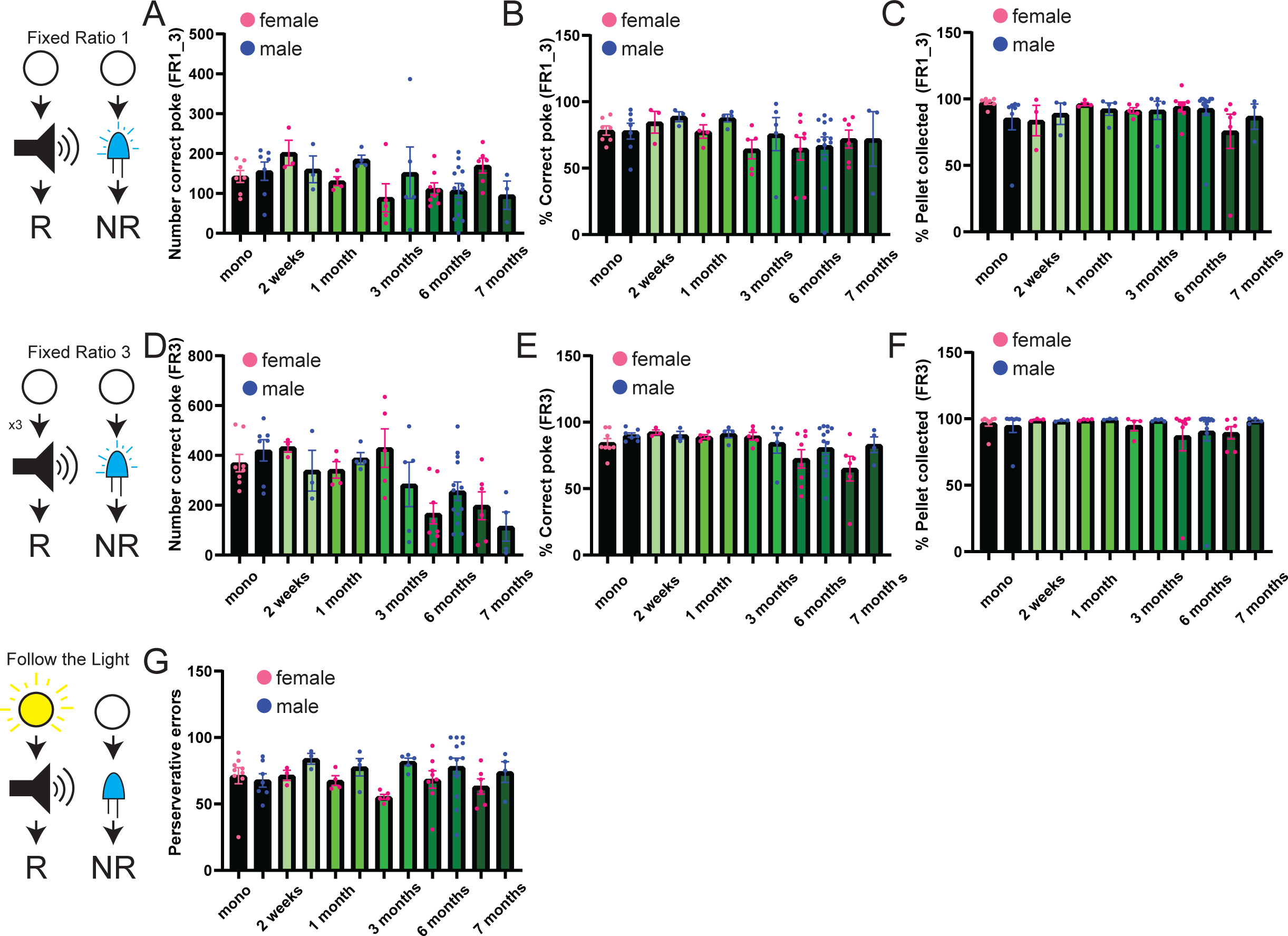

**Figure S21.**
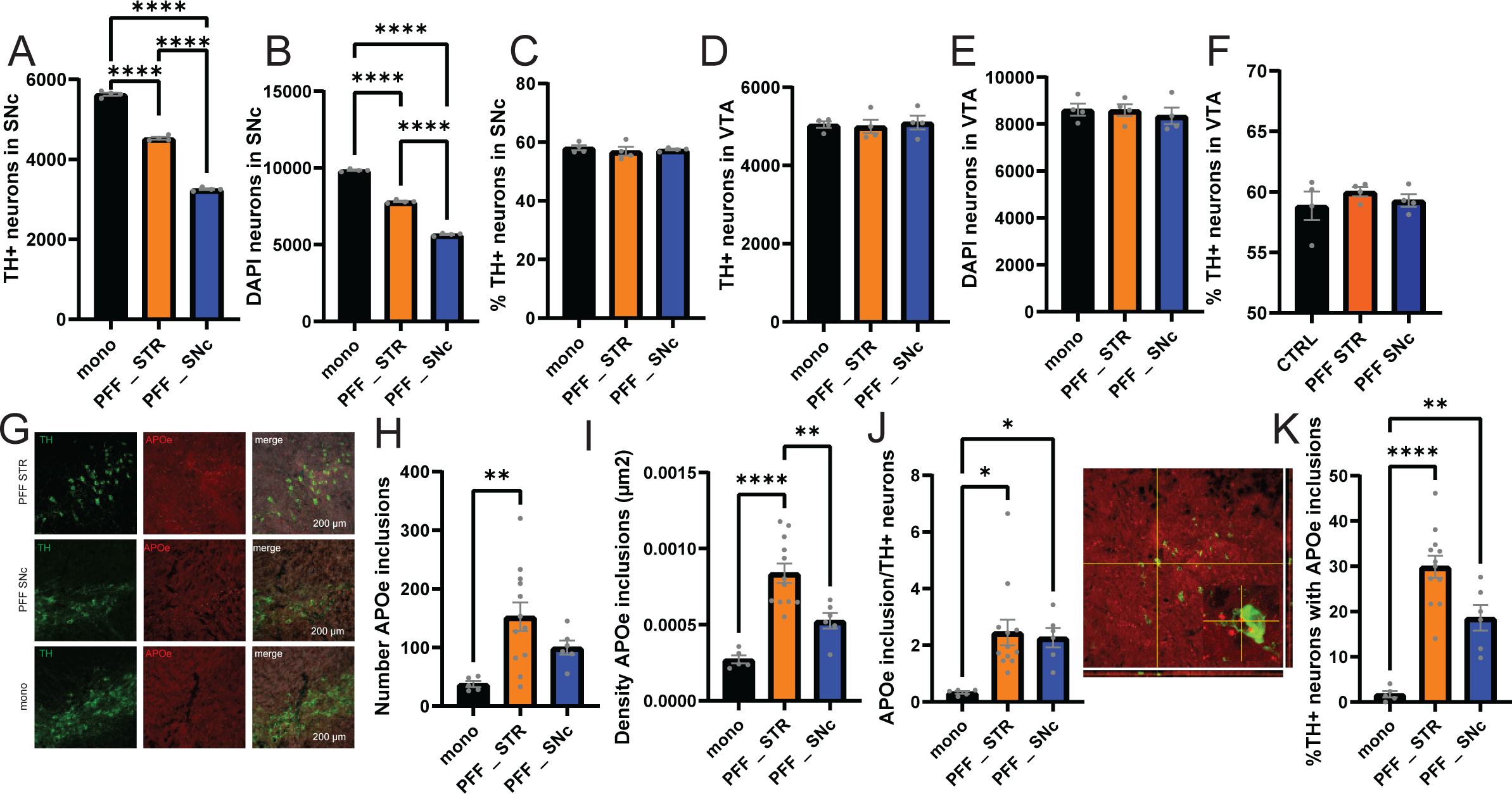

**Figure S22.**
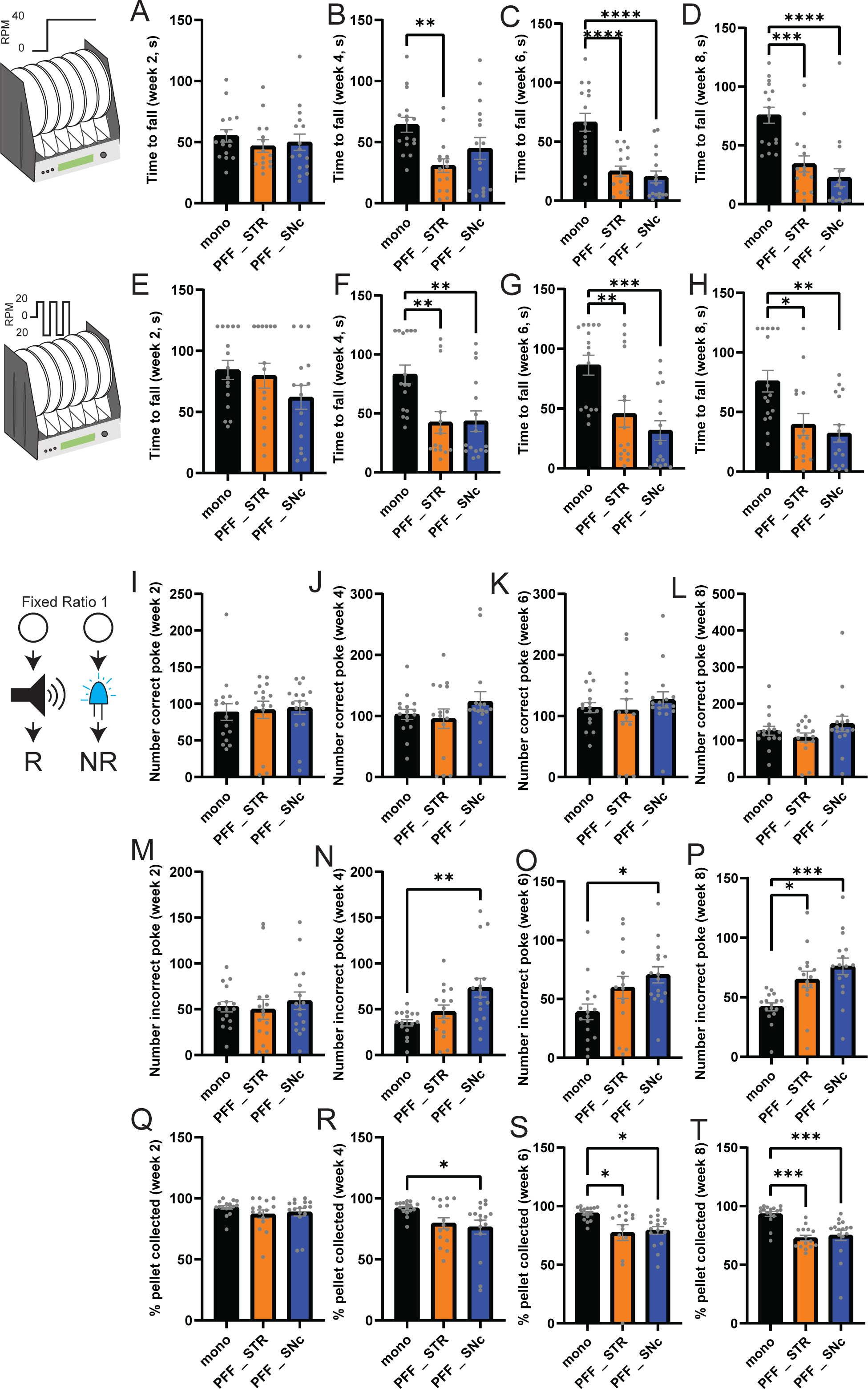

**Figure S23.**
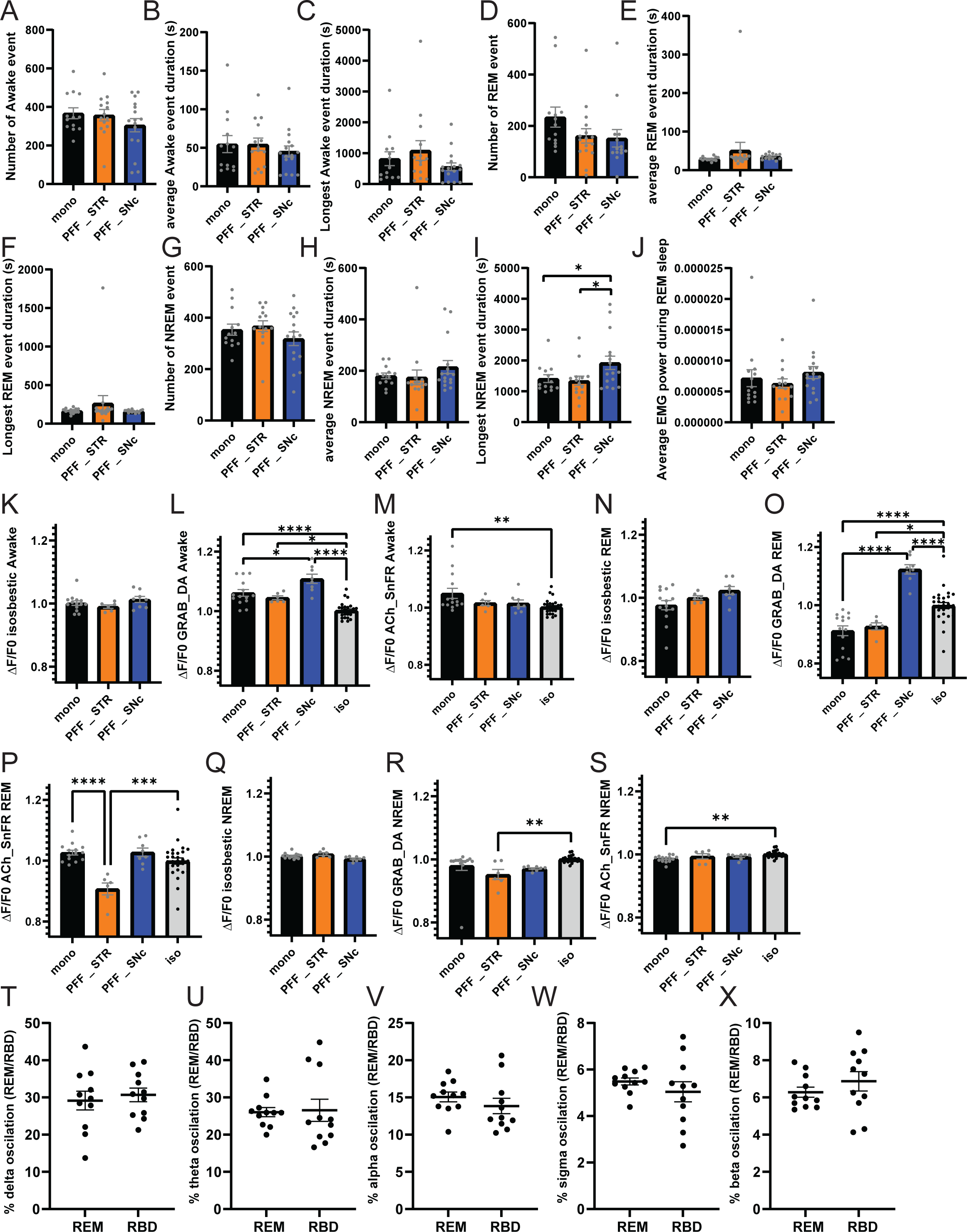

