## Supplementary materials for "Gut-Initiated Alpha Synuclein Fibrils Drive Parkinson’s Disease Phenotypes: Temporal Mapping of non-Motor Symptoms and REM Sleep Behavior Disorder"

### MATERIALS AND METHODS

#### Chemicals and antibodies

Unless stated otherwise, all chemicals were purchased from Sigma Aldrich (Merck KGaA, Germany) and were of an analytical grade. All solutions were prepared using Milli Q deionized water (Millipore, MA, US).

#### Animals

Wild-type C57Bl6 mice (breed in house from Jackson Lab, cat #000664; RRID: ISMR JAX:000664) were bred and maintained in the animal facility at the Karolinska Institute, Sweden, adhering to the guidelines set forth by the local ethical committee at Karolinska Institute (3218-2022, 24297-2022) and the European Communities Council Directive (86/609/EEC). The mice were housed in rooms with a 12-hour light cycle (lights ON at 07:00) and controlled temperature and humidity, 20°C and 53%, respectively. Throughout the entire experimental period, animals had unrestricted access to food and water. To minimize within-experiment variability, experimental animals and their respective controls were sourced from the same litters, ensuring identical age and housing conditions. Notably, both male and female mice were handled consistently across all experiments.

Animals were distributed as follow: 9 animals (6 females, 3 males) injected with monomeric αSyn for 7 months before behavior testing. 7 animals (4 females, 4 males) injected with monomeric αSyn for 2 weeks before behavior testing. 6 animals (3 females, 3 males) injected with PFF αSyn for 2 weeks before behavior testing. 8 animals (4 females, 4 males) injected with PFF αSyn for 1 month before behavior testing. 10 animals (5 females, 5 males) injected with PFF αSyn for 3 months before behavior testing. 21 animals (8 females, 13 males) injected with PFF αSyn for 6 months before behavior testing. 10 animals (6 females, 4 males) injected with PFF αSyn for 7 months before behavior testing.

**αSyn purification and preparation of the αSyn PFF**

αSyn plasmid (full length Prk172 Vector) was transformed into E. Coli bacteria genetically modified to prevent LPS-mediated immune response. Starter culture was maintained at -80ºC until used to generate αSyn monomer. When needed, the starter culture was transfer to LB medium with ampicillin overnight at 37°c before to be resuspended in high-salt buffer and EDTA solution. Next, bacterial cell were broken using high-pressure homogenizer and boiled for 15 minutes to precipitate other proteins. Following cooling on ice and spin at 6,000g for 20 minutes, the supernatant was dialyzed and filtered through Amicon Ultra centrifuge filter (100kDa). Each fraction was then checked using SDS-Page and Coomassie staining to collect αSyn bands (~15Kda). Next the fractions were concentrated and stored at -80°c until needed to generate fibrils.

When required for fibrils generations, aliquot was centrifuge at 12,000g at 4°c and the supernatant removed and confirmed using BCA assay. Monomeric protein was then adjusted to 5mg/ml and shaken for 7 days at 37°c with 1,000 RPM. Following that, fibrils were validated with Thioflavin T assay/Sedimentation assay prior to be kept at -80°c until needed for injections.

When required for injection αSyn PFF were diluted to 2mg/ml by adding sterile PBS and sonicated at 20% amplitude, for a total of 60 pulses (0.5s on/off cycle) with short break every 10 pulses. For all αSyn PFF preparation, a small aliquot was proceeded for transmission electron microscopy to validate αSyn PFF quality, while the remaining was used for injection. During injection, sonicated αSyn PFF were conserved for a maximum of 6 hours to avoid re-aggregation. To avoid within subject variability, all projects were done at the same time using the same lot of monomeric αSyn and αSyn PFF. See dx.doi.org/10.17504/protocols.io.dm6gp3nm5vzp/v1.

**Stereotaxic surgery**

Animals were injected bilaterally with virus and αSyn PFF for behavior and electrophysiology experiments, or unilaterally for anatomy mapping. During surgery, adult mice (~2 months old) were anesthetized with isoflurane (~2% in O2) and then placed on a stereotaxic apparatus (Kopf Instruments, US). Following shaving the skull, and opening a small incision on the scalp, a small drill hole was performed above the injected structures. Viral vector or αSyn PFF were injected using a Hamilton Syringe (NanoNeurons #7001) connected to a nanoinjector (WPI, US). Overexpression of hSNCA was obtained with injection of AAV6-CAG-a-Synuclein, with a titer of 2.3.10^11 viral particules per µl. All injections were performed for a period of 10 minutes followed by 5 minutes of diffusion time. Following injection, the syringe was slowly withdrawn, and scalp sutured. A 2week period was used for the animals to recover from surgery.

Stereotaxic injections were aiming to inject in the Substantia Nigra pars compacta (SNc, 250nl, AP: -3.6mm, ML1.5mm, DV: 4.2mm) and striatum (Str, 500nl, AP: +0.5mm, ML: ±1 and 2mm, DV: 2.5 and 3.5mm). All coordinates were defined from the Bregma and the surface of the brain. For injection in the SNc or Str, staining for tyrosine hydroxylase (TH) was used to define appropriate targeting of the structure.

**Intra-muscularis layer injection in stomach and proximal-duodenum**

At 2 months of age, mice were anesthetized with 2% isoflurane (in O2) and positioned supine with limbs extended. After shaving the abdominal area, a transversal incision was made on the right side, just below the thoracic cage. Subsequently, a second incision, smaller but similar to the first, was made through the abdominal muscles after dissociation from the skin.

Using small, blunt forceps, the liver was gently elevated, exposing the stomach, which was carefully drawn out of the abdominal cavity. After identifying the curvature of the stomach and the pyloric sphincter, a 10 µl Hamilton syringe loaded with 10 µl of αSyn PFF at a concentration of 2.5 µg/µl was inserted along the pyloric canal. In a single penetration, two discrete injections (2.5 µl each) were administered along the lesser curvature of the stomach within the muscularis layer.

A second penetration, perpendicular to the antrum, targeted the muscularis layers of the duodenum adjacent to the sphincter, where two injections (2.5 µl each) were delivered while slowly withdrawing the needle. Following injection, the syringe was withdrawn gradually to prevent any backflow. Abdominal muscles were sutured using a lock-stitch technique to minimize surgical complications. The skin was closed with simple interrupted stitches and surgical glue (Vetbond, 3M). Post-surgery, animals were transferred to clean cages with temperature monitoring for recovery. Over the subsequent 3 days, all animals received painkiller and antibiotic treatments (ketoprofen, Baytril).

**Tissue processing**

After behavioral and physiological experiments, all animals underwent transcardial perfusion. Prior to perfusion with phosphate-buffered saline (PBS), intracardiac blood was aspirated using a sterile syringe and promptly transferred to ice, maintaining it at room temperature for a maximum of 30 minutes. The entire blood volume (approximately 0.5 to 1 mL) was subsequently centrifuged at 1,300g for 15 minutes, and the resulting serum was expeditiously collected and stored at -80°C.

Preceding blood collection, a small skin sample from the thoracic region was procured and promptly preserved at -80°C.

The perfusion process commenced with the infusion of 20 mL of PBS 1X, succeeded by 20 mL of 4% paraformaldehyde (PFA), and finally, 5 mL of PBS 1X to mitigate auto fluorescent artifacts associated with PFA. Post-perfusion, various tissues, including the brain, spinal cord, kidney, liver, lungs, eyes, heart, stomach, intestine, and additional skin from the thoracic region, were harvested, and immersed in 4% PFA for post-fixation, a duration not exceeding 4-6 hours. After post-fixation, all tissues were individually segregated and transferred to a 30% sucrose solution for a period of 2-3 days. Thereafter, the tissues were transitioned to a cryoprotective solution (comprising 50% PBS 10X, 30% ethylene glycol, and 20% glycerol) for storage at -20°C until further processing.

**Serum analyses**

Plasma Orexin A was measured by ELISA (Novus Biologicals, NBP2-80231, MO, US) according to manufacturer instructions. Briefly, samples were centrifuged for 15 min at 1000×g at 4°C. 50µL of standards' working solutions and the samples were added to provided 96 well plate in duplicates. Immediately after 50µL of biotinylated detection antibody working solution was added to each well, covered with the provided plate sealer and incubated for 45 min at 37°C. Next, the solution from each well was decanted, 350 uL of wash buffer added to each well and allowed to soak for 1min. Then, the solution from each well was decanted and pat dry against clean absorbent paper. The wash step was repeated 3 times. 100µL of HRP conjugate working solution was added to each well, plate was covered with the sealer and incubated for 30 min at 37°C. Next, the was step was performed as described above and 90 µL of substrate reagent was added to each well, followed by incubation for 15 min at 37°C protected from light. 50 µL of stop solution was added to each well and the optical density was measured using a micro-plate reader with absorbance set to 450 nm. See dx.doi.org/10.17504/protocols.io.kxygx365zg8j/v1.

Plasma Melatonin was measured by ELISA Kit (Enzo Life Sciences, ENZ-KIT150-0001, NY, US) according to manufacturer instructions. Briefly, prior to ELISA the melatonin was phase extracted from solution. 200ul of plasma was mixed with an equal volume of cold Ethyl Acetate and vortexed gently. Layers were allowed to separate on ice for 3 min, vortex again and incubated on ice for 2 min. Next, samples were spinned at 1000g for 10 min at 4°C and the organic layer transferred to a new tube. The samples were dried and resuspended in 220µl of 1X stabilizer. 100µL of standards' working solutions and the samples were added to provided 96 well plate in duplicates. Immediately after 50µL of melatonin tracer was added to each well (except blanks) followed by 50µl of 1X melatonin antibody (except blanks). The plates were covered with the provided plate sealer and incubated for 1 h at 37°C with 500 rpm shaking. Next, the solution from each well was decanted and 400 uL of wash solution added to each well. Then, the solution from each well was decanted and pat dry against clean absorbent paper. The wash step was repeated 3 times. 200µL of melatonin conjugate solution was added to each well (except blanks), plate was covered with the sealer and incubated for 30 min at room temperature. Next, the was step was performed as described above and 200 µL of substrate reagent was added to each well, followed by incubation for 30 min at 37°C protected from light. 50 µL of stop solution was added to each well and the optical density was measured using a micro-plate reader with absorbance set to 450 nm. See dx.doi.org/10.17504/protocols.io.81wgbx35ylpk/v1.

Total and PS129 αSyn levels were measured using western-blotting (WB) as described previously (Paslawski and Svenningsson, 2023). Briefly, plasma samples were diluted 10-fold in MQ water before proceeding to further steps. The sample was mixed in a 1:1 ratio with 20 mM TCEP, to reduce disulfide bridges, and then 1:1 with a denaturing loading buffer (50 mM Tris-HCl, 70 mM Tris, 1% lithium dodecyl sulphate (LDS), 5% glycerol, 0.25mM EDTA, 0.11mM SERVA Blue G-250, 0.0875mM Phenol Red, pH-8.5) and boiled at 95°C for 5 min. Next, samples were separated using a 4-12% bis-tris acrylamide gel using MES running buffer (50mM Tris, 50mM 2-(N-morpholino) ethanesulfonic acid (MES), 0.1% SDS, 1mM EDTA, pH-7.3). 20 µL of the sample was used. After Sodium dodecyl sulphate polyacrylamide gel electrophoresis, gels were incubated in a transfer buffer (25mM Tris, 192mM glycine, 30% methanol) and 0.45µm pore size Immobilon-P™ PVDF membranes (Millipore, MA, US) were pre-wetted in methanol. Gels were assembled with membranes and the transfer was performed using the Trans-Blot Turbo Transfer System (BioRad, CA, US) according to manufacturer protocols. Next, membranes were dried and shortly fixed in 100% methanol. Between each incubation period, membranes were washed three times in Tris Buffered Saline (TBS, 20mM Tris, 150mM NaCl, pH-7.6) containing 0.1%(v/v) Tween20 (TBS-T). Membranes were blocked during 1h incubation in 5% skim milk at RT. Next, membranes were incubated overnight at 4°C with a primary antibody diluted 1:1000(v/v) in 1% skim milk. Antibodies used were, Anti-Total-Alpha-Synuclein Polyclonal Antibody (Thermo Fisher Scientific, PA5-143581, MA, US) and Anti-Alpha-synuclein (phospho S129) antibody [EP1536Y] (Abcam, ab51253, UK). Membranes were thereafter incubated for 2h with appropriate HRP-conjugated secondary antibody (Dako/Agilent, CA, US) diluted 1:10000(v/v) in 1% skim milk at RT. The signal was developed by a Clarity™ Western ECL Substrate (Bio-Rad, CA, US) and levels of visualized proteins were estimated by measuring band intensities with ImageJ and normalized to the amount of total protein in the sample, measured by BCA assay according to manufacturer instructions (Thermo Fisher Scientific, MA, US). A fluorescent scan was used to visualize the protein ladder. See dx.doi.org/10.17504/protocols.io.n92ldmk78l5b/v1.

**Microbiota analyses**

DNA extraction – Fecal collection from mice was meticulously conducted in sterile cages for each mouse, ensuring expeditious processing at a standardized time of day (morning, 09:00 to 11:00) to minimize potential microbial contamination and circumvent diurnal variations. All animals were allowed a minimum of 15 minutes in the cage, and collection continued until multiple pellets were obtained. Pellets in contact with urine were excluded from further analyses.

Immediately following extraction, the pellets were transferred to sterile Eppendorf tubes positioned on dry ice. Upon completion of the experiments, Eppendorf tubes were subsequently transferred to -80°C for storage until further processing. DNA extraction was carried out using the QiAamp PowerFecal Pro DNA kit (Qiagen, Sweden) and following the recommended protocol. Approximately 25 mg of feces were transferred to the bead beating tube under a fume hood. The feces were then homogenized thoroughly using both recommended mechanical and chemical methods. The resulting lysate was transferred to the inhibitor solution, and total DNA was captured using the provided silica membrane before undergoing washing and elution for subsequent processing.

Amplicon Generation - 16S rRNA/18SrRNA/ITS genes of distinct regions (16SV4/16SV3/16SV3- V4/16SV4-V5, 18SV4/18SV9, ITS1/ITS2, ArcV4) were amplified used specific primer (e.g. 16SV4: 515F- 806R, 18SV4: 528F-706R, 18SV9: 1380F- 1510R, et. al) with the barcode. All PCR reactions were carried out with 15 µL of Phusion® High - Fidelity PCR

Master Mix (NewEngland Biolabs); 0.2 μM of forward and reverse primers, and about 10 ng template DNA. Thermal cycling consisted of initial denaturation at 98C for 1 min, followed by 30 cycles of denaturation at 98C for 10 s, annealing at 50C for 30 s, and elongation at 72C for 30 s and72C for 5 min.

Quality control- PCR amplification of targeted regions was performed by using specific primers connecting with barcodes. The PCR products of proper size were selected through 2% agarose gel electrophoresis. The same amount of PCR products from each sample was pooled, end-repaired, A-tailed, and further ligated with Illumina adapters. Libraries were sequenced on a paired-end Illumina platform to generate 250bp paired-end raw reads.

OT Production - Sequences analysis was performed by Uparse software (Uparse v7.0. 100, https://drive5.com/uparse/). Sequences with ≥ 97% similarity were assigned to the same OTUs. Representative sequence for each OUT was screened for further annotation. For each representative sequence, the Silva Database (http://www.arb- silva.de/) was used based on Mothur algorithm to annotate taxonomic information. Phylogenetic Relationship Construction In order to study phylogenetic relationship of different OTUs, and the difference of the dominant species in different samples (groups), multiple sequence alignment was conducted using the MUSCLE software (http://www.drive5.com/muscle, Version 3.8.31). Data Normalization OTUs abundance information was normalized using a standard of sequence number corresponding to the sample with the least sequences. Subsequent analysis of alphadiversity and beta diversity were all performed basing on this output normalized data. Relative abundance Top 10 taxa of each sample at each taxonomic ranks (Phylum, Class, Order, Family, Genus, and Species) were selected to plot the distribution histogram of relative abundance in Perl through SVG function. Heatmap The abundance information of top 35 taxa of each sample at each taxonomic ranks were used to draw the heatmap, which visually display different abundance and taxa clustering. This was achieved in R through the pheatmap() function. Ternary plot Ternary plot for top 10 taxa at each taxonomic ranks can be used to show the abundance difference among three samples. It was performed in R with vcd() function. Venn and Flower diagram Venn and Flower diagrams visually display the common and unique information between different samples or groups. Venn and Flower diagrams were produced in R with VennDiagram() function and in perl with SVG function, respectively. Phylogenetic tree Phylogenetic tree, also called evolutionary tree, can describe the evolutionary relationship between different species. One hundred genera with the highest abundance in the samples were selected and performed sequence alignment to draw the phylogenetic tree in perl with SVG function.

Alpha-Diversity – Alpha diversity is applied in analyzing complexity of species diversity for a sample through 6 indices, including Observed-species, Chao1, Shannon, Simpson, ACE, Good-coverage. All these indices in our samples were calculated with QIIME (Version 1.9.1) and displayed with R software (Version 4.0.3, https://www.r-project.org/, RRID:SCR_000432). Two indices were selected to identify Community richness: Chao -the Chao1 estimator; CE -the ACE estimator; Two indices were used to identify Community diversity: Shannon - the Shannon index; Simpson – the Simpson index; One indice to characterized Sequencing depth: Coverage - the Good's coverage. Species accumulation boxplot In order to evaluate the richness of microbial community and sample size. Species accumulation boxplot can be used to visualize, which performed with vegan package in R software. Rank abundance curve Rank abundance curve can reflect the richness and evenness of species with samples through observing the width and shape of the curves.

Beta-Diversity - Beta diversity analysis was used to evaluate differences of samples in species complexity, Beta diversity on both weighted and unweighted unifrac were calculated by QIIMEsoftware (Version 1.9.1). Cluster analysis was preceded by principal component analysis (PCA), which was applied to reduce the dimension of the original variables using the ade4 package and ggplot2 package in R software (Version 4.0.3, https://www.r-project.org/, RRID:SCR_000432). Principal Coordinate Analysis (PCoA) was performed to get principal coordinates and visualize from complex, multidimensional data. A distance matrix of weighted or unweighted unifrac among samples obtained before was transformed to a new set of orthogonal axes, by which the maximum variation factor is demonstrated by first principal coordinate, and the second maximum one by the second principal coordinate, and so on. PCoA analysis was displayed by ade4 package and ggplot2 package in R software (Version 4.0.3).

Community difference analyses - A series of statistical analyses, which include Anosim, Adonis, Multi-response permutation procedure (MRPP), Simper, T-test, MetaStat and LEfSe, were performed to reveal the community structure differentiation. Anosim, Adonis and MRPP analysis are non-parametric tests that analyze the difference between high-dimensional data group. They can test whether the differences between groups are significantly greater than the differences within the group, which can determine whether the grouping is meaningful. All of them were performed with vegan package and ggplot2 package within R. Simper can reveal the contribution of each species to the differentiation between groups. Top 10 species were selected and presented on the graph. It was performed in R with Vegan package and ggplot2 package.

**Behavior testing**

For all behavioral experiments, animal cages were transferred to the behavioral antechamber one hour prior to testing for acclimatization. Male and female subjects were tested separately, with a minimum of 10 minutes allocated as a buffer time between transitions of sexes. Following each trial, the apparatus underwent thorough cleaning using a 50% ethanol solution to prevent any potential cross-contamination between animals from different cages.

Consistency in testing conditions was maintained by conducting all experiments at the same time of day, with a permissible variation of within 4 hours, either in the morning or afternoon. This approach was adopted to mitigate variability associated with diurnal cycles.

**Olfactory discrimination test**

Animals were transferred to a 30 x 30 cm grey open field chamber equipped with overhead housing lights and a camera recording system positioned above the apparatus. The camera was connected to behavior tracking software (EthoVision XT , RRID:SCR_000441) for precise data analysis. To prevent day-to-day habituation, no buffer habituation sessions were implemented.

On the first day of the experiment, mice were introduced to the apparatus for a duration of 20 minutes, accompanied by a 5 cm plexiglass circle cage filled with chocolate-flavored pellets (05684, Bio-Serv). On the subsequent day, a different apparatus was utilized, and the chocolate-flavored pellets were replaced with cotton pads infused with 1 ml of a 1% solution of 2,3,5-trimethyl-3-thiazoline (TMT, Sigma Aldrich) in water.

For each experimental session, parameters such as the distance traveled, time spent in the center, average speed, and time spent in the corner associated with the olfactory stimulus were extracted from the tracking software. The olfactory discrimination index was calculated as the ratio of time spent in the corner linked with chocolate flavor to the time spent in the corner associated with TMT. A ratio of 1 was considered indicative of no significant preference for either odor. See dx.doi.org/10.17504/protocols.io.36wgq3wwxlk5/v1.

**Sucrose preference test**

Individually housed mice in single cages were provided access to two novel bottles, distinct from those used for husbandry. One bottle contained a 1% sucrose solution, while the other contained regular water. After a 24-hour period, each bottle was weighed, and the weights were compared to the initial measurements to determine water and sucrose consumption. The ratio of water to sugar consumption was then expressed as the sucrose preference index. See dx.doi.org/10.17504/protocols.io.4r3l22wwxl1y/v1.

**Dark light box**

A 45 x 20 x 20cm custom box divided into 1/3 dark (covered with lid, <5lux) and 2/3 white (~100lux) was used for the experiment. Animals were released in the dark compartment before positioning the lid and left to explore for 6 minutes. All experiments are video recorded, and manual scoring was done for the time in the light box as well as the number of time the mice explored the light box. See: **dx.doi.org/10.17504/protocols.io.ewov19jz7lr2/v1**.

**Rotarod 40 RPM**

The animals were positioned on an apparatus (Ugo Basile, cat # 47650, https://ugobasile.com/products/categories/motory-coordination/rotarod-for-mice-and-rats) with the receptacle platform in the upward position. The session was repeated three times on three consecutive days at a consistent time of day (0900 to 1200). Mice were allowed to acclimate for a few seconds before initiating the rotation protocol. Up to five mice were tested simultaneously if they originated from the same cage.

Upon commencing the protocol, the speed was increased to 40 rotations per minute (40 RPM) and maintained at this speed for 120 seconds. For each mouse, the time to fall (determined by the receptacle) or the time for the mouse's body to complete three full rotations (head down) around the beams was recorded. See dx.doi.org/10.17504/protocols.io.3byl4qo5zvo5/v2.

**Rotarod 20 RPM – docking**

The animals were positioned on an apparatus (Ugo Basile, cat # 47650, https://ugobasile.com/products/categories/motory-coordination/rotarod-for-mice-and-rats) with the receptacle platform in the upward position. The session was repeated three times on three consecutive days at a consistent time of day (13:00 to 16:00). Mice were allowed to acclimate for a few seconds before initiating the rotation protocol. Up to five mice were tested simultaneously if they originated from the same cage.

Upon commencing the protocol, the speed was increased to 20 rotations per minute (20 RPM) and maintained at this speed for five rotations before stopping. Subsequently, the platform rotated in the opposite direction for five rotations at 20 RPM (docking). For each mouse, the time to fall (determined by the receptacle) or the time for the mouse's body to complete three full rotations (head down) around the beams was noted. See dx.doi.org/10.17504/protocols.io.3byl4qo5zvo5/v2.

**Bedding test**

Mice were individually housed for a minimum of 48 hours before testing. On the test day at 09:00, the mice were transferred to a clean cage with ad libitum access to food and water, featuring a 5x5 cm cotton pad positioned on the floor. After 24 and 48 hours, a picture was taken of the apparatus, and a bedding quality score (on a scale of 0 to 5 points) was assigned based on specific parameters: 0) did not touch the pad, 1) unfolded the pad, 2) folded the pad in a corner, 3) folded the pad in a corner with bite marks, 4) initiated shredding the pad and created a nest, and 5) the pad is shredded, and a proper nest is visible. See dx.doi.org/10.17504/protocols.io.n2bvj3ko5lk5/v1.

**Descending pole test**

Mice were positioned on top of a wooden pole (50 cm long, ~1 cm diameter) leading to a clean cage. It's noteworthy that the mice were not subjected to training to prevent contamination of the learning experience. The mice were placed with their heads up, and the duration from the moment the head faced downward to the time the forelimb touched the bedding was measured using recorded video sessions.

**Descending inclined platform test**

At the initiation of the test, mice were positioned on the horizontal section of a 60 cm long, 45° inclined platform featuring a 1 cm grid. Subsequently, mice were gently prompted to the starting point of the inclined path and allowed to move down freely. The time taken to reach the clean cage positioned at the bottom was then measured utilizing video recorded sessions. Importantly, mice were not subjected to training or habituation on the platform to prevent any contamination of learning effects. For all animals, 5 different frames were randomly selected for the period the mice descend the platform. Each frame was then used to determine the distance between hindlimb express as number of grid frame. Next, the number of grid frames were converted to cm. See DOI: dx.doi.org/10.17504/protocols.io.14egn3ko6l5d/v2.

**Operant behavior**

Operant behavior testing was conducted in individually housed mice without access to regular food. In each animal cage, a FED3.0 operant box was positioned ( https://open-ephys.org/fed3/fed3, Matikainen-Ankney *et al.*, 2019). To prevent wood bedding from entering the nose poke, the amount of bedding provided to each animal was deliberately reduced. The

FED3.0 operant box was defined with two nose pokes using infrared beams to determine nose entry, and a food pellet dispenser was positioned in the middle.

Initially, mice were exposed to the FED3.0 for 24 hours with 6g of chow pellets placed on the floor. In the following two days, mice were exposed to free-feeding sessions where 20mg sugar pellets (BioServ, #f07595, https://www.bio-serv.com/product/DPP_Sucrose.html) were randomly delivered to habituate the mice. The next three days involved a switch to a Fixed Ratio 1 (FR1) protocol starting at 09:00. Each correct poke was paired with an auditory cue and the delivery of a single sugar pellet, while an incorrect poke was paired with a visual cue (Blue LED positioned below the pellet dispenser). When a pellet was delivered, both correct and incorrect nose pokes became inactive to prevent multiple pellet deliveries.

Following three consecutive sessions of FR1, the protocol was increased to Fixed Ratio 3 (FR3) for one day. In this protocol, each time the mice produced three correct nose pokes, a 20mg pellet was delivered with the same auditory cue. Any incorrect poke was paired with a visual cue to indicate errors. Similar to FR1, when a pellet was delivered, both correct and incorrect nose pokes were made inactive to avoid multiple pellet deliveries.

Finally, on the last day, the protocol was switched to a "Follow the Light" protocol for 24 hours. In this protocol, correct and incorrect sides were alternated randomly using a fixed ratio 1 schedule. The poke associated with pellet delivery was indicated by a yellow LED turning ON in the nose poke. When a poke occurred on the correct side, a 20mg sugar pellet was delivered with an auditory cue, while an incorrect poke was paired with a visual cue. Similar to the previous protocols, when a pellet was delivered, both correct and incorrect nose pokes became inactive to prevent multiple pellet deliveries. The number of correct, incorrect pokes and pellets collected within 10s of delivery were then extracted for further analysis. See: dx.doi.org/10.17504/protocols.io.j8nlk8b31l5r/v1.

**Telemetry implantation**

Mice underwent deep anesthesia with 2% isoflurane in O2 before being secured in a stereotaxic frame (Kopf Instruments). The skin over the skull was incised and retracted to expose the skull. An electroencephalogram (EEG) electrode (Agnhtos.se, MCS1x2) was then screwed into the skull above the frontal cortex (AP: 1.5, ML: 1.0), and a reference electrode was secured into the skull above the cerebellum. Electromyogram (EMG) electrodes were positioned in the nuchal muscles on the same side with approximately 1mm spacing; these electrodes were then affixed using small heat-shrink tubing. Subsequently, a sizable chamber was created in the abdominal area to accommodate the wireless telemetry recording device (F20-EET, DSI, USA). The cavity was thoroughly flushed with saline to prevent complications, and the skull electrodes were secured using dental cement (Paladur). Finally, the skin was sutured and covered with surgical glue. For all animals, EEG was recorded on channel 2, while EMG was recorded on channel 1 of the wireless telemetry device. https://www.protocols.io/view/procedure-for-eeg-surgery-kxygx3dxog8j/v1

**Telemetry recording**

Following a recovery period of at least 1 week post-surgery, animals were transferred to the sleep-recording room and allowed to acclimate for a minimum of 48 hours. During the recordings, animals were group-housed with 1-2 cage mates throughout the entire procedure, enabling the characterization of sleep independently of any social stress factors. On the day of the recording, the telemetry device was activated using a strong magnet, and the cages were placed on the telemetry receiver (DSI, https://www.datasci.com/products/software/neuroscore). A total of 16 animals were recorded simultaneously.

EEG, EMG, and movement activity were recorded via telemetry using Neuroscore v3.0 software (DSI) at a frequency of 500Hz for EEG/EMG and 1Hz for activity. Mice were recorded for 24 hours without interruption in a soundproof room, with ad libitum access to food and water.

**Sleep data analysis**

Sleep data files were imported into Neuroscore V3.0 software and visually inspected. Data with interruptions, noise, or issues related to battery status were excluded from further analyses. Subsequently, the DSI-provided "sleep scoring 2" script was utilized for analysis, incorporating EEG, EMG, and activity channels for scoring (Tools – Rodent scoring 2 visual Tuning).

For our analyses, specific power levels were considered indicative of different sleep stages:

- Delta power levels with a delta ratio of 0.5 were associated with a maximum probability of REM/Wake, and a NREM delta ratio of 1 represented a maximum probability of NREM.

- Theta power levels with a theta to delta ratio of 1.3 indicated a maximum probability of Wake/NREM, and a theta to delta ratio of 3 indicated a maximum probability of REM.

- EMG power levels with an EMG ratio of 1.1 were considered representative of a maximum probability of REM to NREM, and an EMG ratio of 2.4 indicated a maximum probability of Wake.

- An activity count of 0.1 was defined for the active wake phase. To avoid separating different wake stages, the scoring rules combined active wake and wake based on the activity channel.

Artifact detection thresholds were set on a per-recording basis, with most recordings having thresholds set to 0.5mV for both EEG and EMG.

Stage transition probabilities based on EEG, EMG, and activity were defined as follows:

- Wake to NREM: 90%
- Wake to REM: 90%
- NREM to Wake: 80%
- NREM to REM: 90%
- REM to Wake: 70%
- REM to NREM: 80%

Delta oscillation was determined within a power band of 0.5 to 4Hz, while Theta oscillation was within the 6 to 9Hz range. The contributing factors of Delta, Theta/Delta, and EMG were considered equally important for all recordings. Sleep scoring was then obtained within 20s windows, and any recording with more than 1% artifacts was excluded from further analyses. See dx.doi.org/10.17504/protocols.io.yxmvm3rrbl3p/v1.

**REM sleep behavior disorder scoring and Periodogram analyses**

Following the scoring of sleep stages, EMG was converted into the root mean square (RMS) using the signal grid function. For all recordings, a signal grid was generated, including the timestamps (20s windows) of the GMT time, the RMS of the EMG, the sleep scoring, and the EEG. Additionally, a periodogram Power Band (PB) for each 20s window was obtained within the 0.3 to 80 Hz band range. The analysis employed a 10/1024 FFT order, an overlap of 50%, and a Hamming spectral function applied to the window. PB values were expressed as a relative percentage of the power band value. Delta oscillation was determined within the 0.5 to 4Hz window, Theta within the 4 to 8Hz window, Alpha within the 8 to 12Hz window, Sigma within the

12 to 16Hz window, Beta within the 16 to 24Hz window, Low Gamma within the 24 to 49 Hz window, and High Gamma within the 51 to 80Hz window.

The signal grid was then exported to Excel for further analyses. In Excel, the RMS-EMG was first converted to the z-score over the entire recording. A COUNTIF function was then used to count the number of REM events (scored as P), awake events (scored as W), or NREM events (scored as S) for the entire recording, as well as during the dark/light phases. Additionally, a COUNTIF function was used to count the number of REM events with a z-score EMG higher than 2 standard deviations (SD), which were further defined as REM Sleep Behavior Disorder (RBD) or REM events without atonia. For each animal, the number of RBD events, as well as the ratio of RBD events to total REM events, was further extracted. Furthermore, for each sleep, awake, and REM event, the average PB value for each band-pass frequency was extracted and subjected to further comparisons. See dx.doi.org/10.17504/protocols.io.81wgbxjdolpk/v1.

**Fiber Photometry during sleep**

A subset of mice from each group received a combination of two Adenoviruses (AAV) at a ratio of 1:1 for imaging dopamine release (AAV9-hSyn-GRAB-rDA1h, 250nl, addgene #140557) or acetylcholine release (AAV1-CAG-iAChSnFr, 250nl, addgene #137955) in the dorsomedial striatum (coordinates from the Bregma: AP 0.5mm, ML 1.1, DV 3.0mm) in the same injection. After a 2-week interval, mice were implanted with a sleep recording telemetry device and an optic fiber (0.50NA, 400µm, 4mm long, Thorlabs) above the virus injection site. The ceramic ferrule holding the optic fiber was secured in place with dental cement. Following recovery, animals were acclimated to the sleep recording room for 48 hours. During acclimation, mice were daily connected to the fiber photometry setup for habituation.

The fiber photometry setup enabled excitation of the isosbestic channel (405nm), GCAMP (460-490), and RCAMP (555-570), with fluorescence collection at 500-540 and 580-680 (Built-in detector Gen 2 6-port Fluorescence mini cube, Doric lenses). Excitation was controlled using the Fiber Photometry console and analyzed through manufacturer software (Doric Neuroscience Studio v2.0). All three excitation wavelengths were used in a lock-in protocol with non-overlapping frequencies, and the excitation LED power was set at 10% below the level that provided clear calcium events. This adjustment aimed to prevent arbitrary calcium release that could reduce signal variation in large windows (20s). The setup allowed simultaneous collection of isosbestic, Ach-SnFr, and rGRAB-DA signals at 1017Hz. A TTL input synchronized photometry and sleep signals. For each animal, a minimum of 30 minutes of data were acquired and repeated multiple times to cover multiple sleep cycles. Recordings were limited to 30 minutes to prevent the excitation light from affecting the mice's sleep schedule and to minimize interference from the patch cord on their ability to sleep.

Following recording, sleep scoring was processed as described above, extracting timestamps of Wake, REM, NREM, and RBD events. Using a custom-made Matlab script, photometry data were downsampled to 500Hz (average smooth window function), then aligned with the sleep scoring. For each 1-minute window, each channel of the photometry was normalized using the z-score function and then averaged over the 20s window corresponding to the sleep scoring. For each animal, photometry data were then averaged for specific sleep stages and compared across groups. See: dx.doi.org/10.17504/protocols.io.6qpvr8xw3lmk/v1.

**Data exclusion**

All data were tested in with GraphPad Prism software (www.graphpad.com, version 9, GraphPad (RRID:SCR_000306)) for Rout method for outliers with a Q value of 5%. For the experiment with the operant boxes, animals were excluded if they do not proceed to a correct poke over the entire 24h period, however, the animals were still tested for the next protocol. Only animals that do not execute correct poke for 2 consecutive sessions were excluded. For sleep experiments, data with more than 1% artifacts detected were automatically excluded. For sleep electrophysiology experiments, 20s events that were considered as artifact by the software were automatically discarded from analyses, but the animals were maintained for further analyses. For RBD events, animals that display no REM events during the entire recording were automatically discarded from further analyses. For virus experiments (AAV-hSNCA or photometry), posthoc histology of the virus location and expression was used for discarding animals with misplaced injections. For photometry experiments, all animals with a misplaced optic fiber, or a signal that show no variation compared to the isosbestic were discarded from further analyses.

**Data sharing**

Detailed protocols are available on https://www.protocols.io platform. Prism file including all analyses are available on the https://zenodo.org platform. All raw data and scripts are shared under the ASAP sharing depository.

**Statistics analyses**

Statistical analyses were conducted using GraphPad Prism software (version 9). For comparisons involving two groups, paired- or unpaired- t-test analyses were applied. In cases involving three or more groups, a 1-way ANOVA analysis with a Tukey post hoc analysis was performed. At the exception of serum analyses where LSD post hoc test were used as well as the test where ANOVA display significant P-values but Tukey post hoc test show no significant effects (always indicated in figure legends). In the analyses of gut-injected animals, post hoc analyses were provided for the comparison of monomeric αSyn-injected animals with the αSyn PFF-injected group, with additional statistical values available in the attached Prism files. For comparison of the surgery effect (i.e monomeric αSyn 2 weeks versus 7 months) a 2-way ANOVA time * group was conducted. For comparisons of sex differences, a 2-way ANOVA sex×group analysis was conducted, and the interaction effect was reported in the figure legend. In anatomical analyses of ROIs, a 1-way ANOVA or 2-way ANOVA analysis was performed on each ROI and presented as imagesc – Jet, while post hoc analyses were conducted using Dunnett's test by default. Outlier data were removed using the Prism Rout method with a 5% Q value.

Statistical significance was considered with a P-value less than 0.05, denoted as *, $P < 0.05$, **, $P < 0.01$, ***, $P < 0.001$, and ****, $P < 0.0001$.

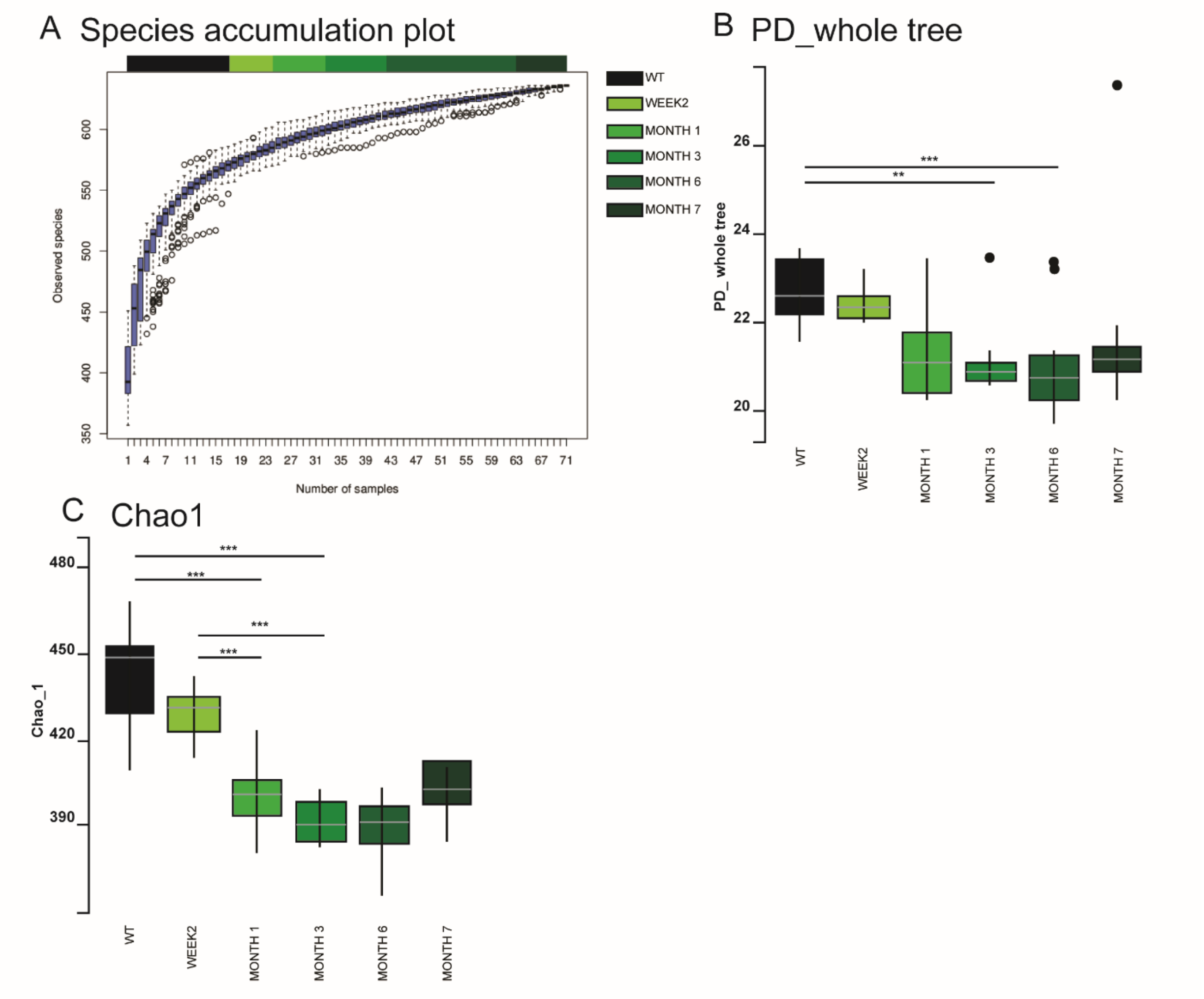


**Fig. S1. Related to Figure 2**

Species accumulation plot (**A**), PD whole tree (**B**) and Chao1 analyses (**C**) of alpha diversity found in microbiota of mice injected with monomeric αSyn or αSyn PFF in the stomach. Box plots are representing the median, lower and upper quartile as well as the minimum and maximum value. All individual point represents individual animals. ** P<0.01, *** P<0.001. Statistical data represent post hoc analyses compared to monomeric αSyn group following 1-way ANOVA.

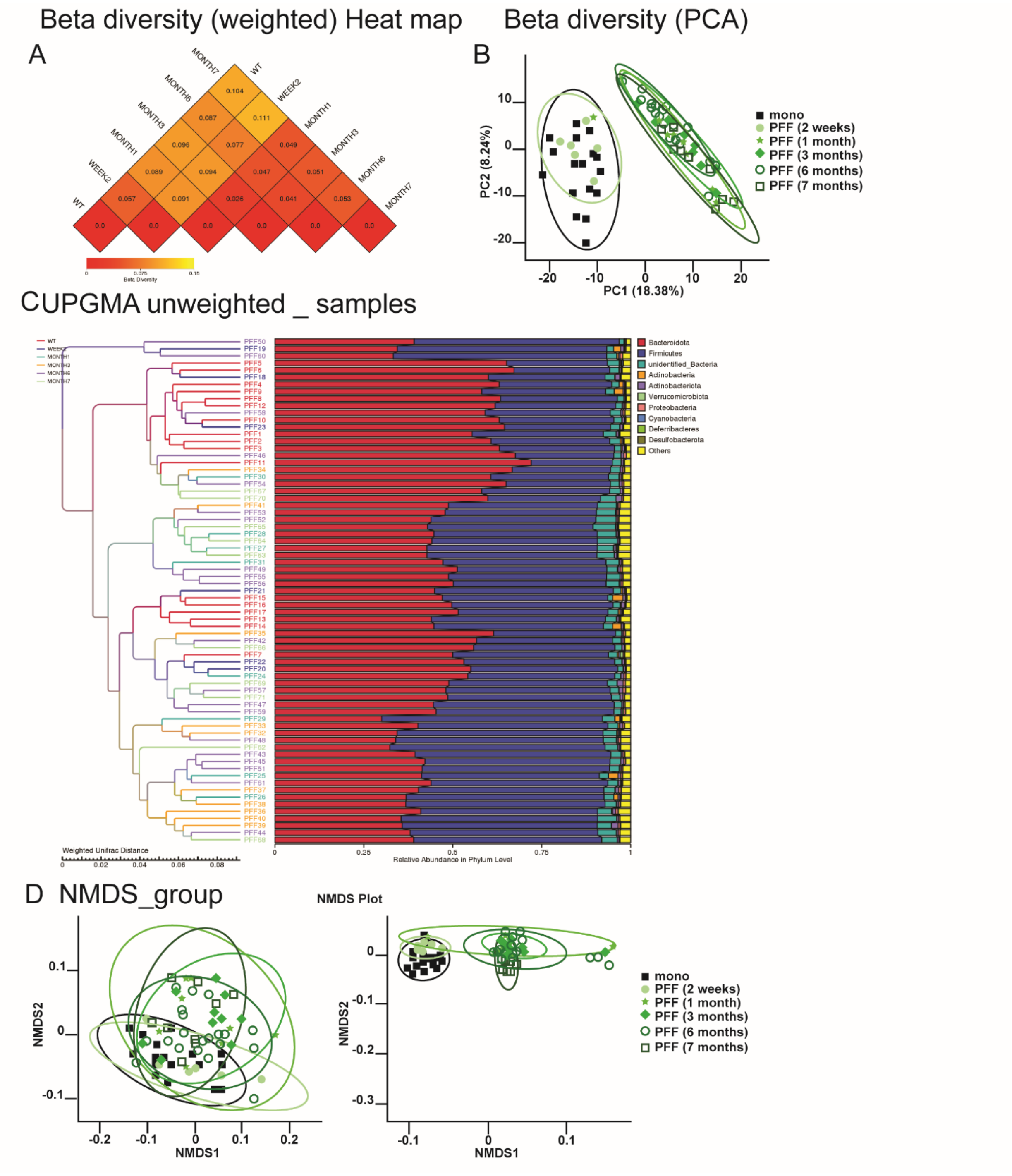


**Fig. S2. Related to Figure 2**

Beta diversity heat map (**A**), personal component analyses (PCA, **B**), UPGMA unweighted samples (**C**) and NMDS analyses (**D**) of beta diversity found in microbiota of mice injected with monomeric αSyn or αSyn PFF in the stomach. All individual point represents individual animals.

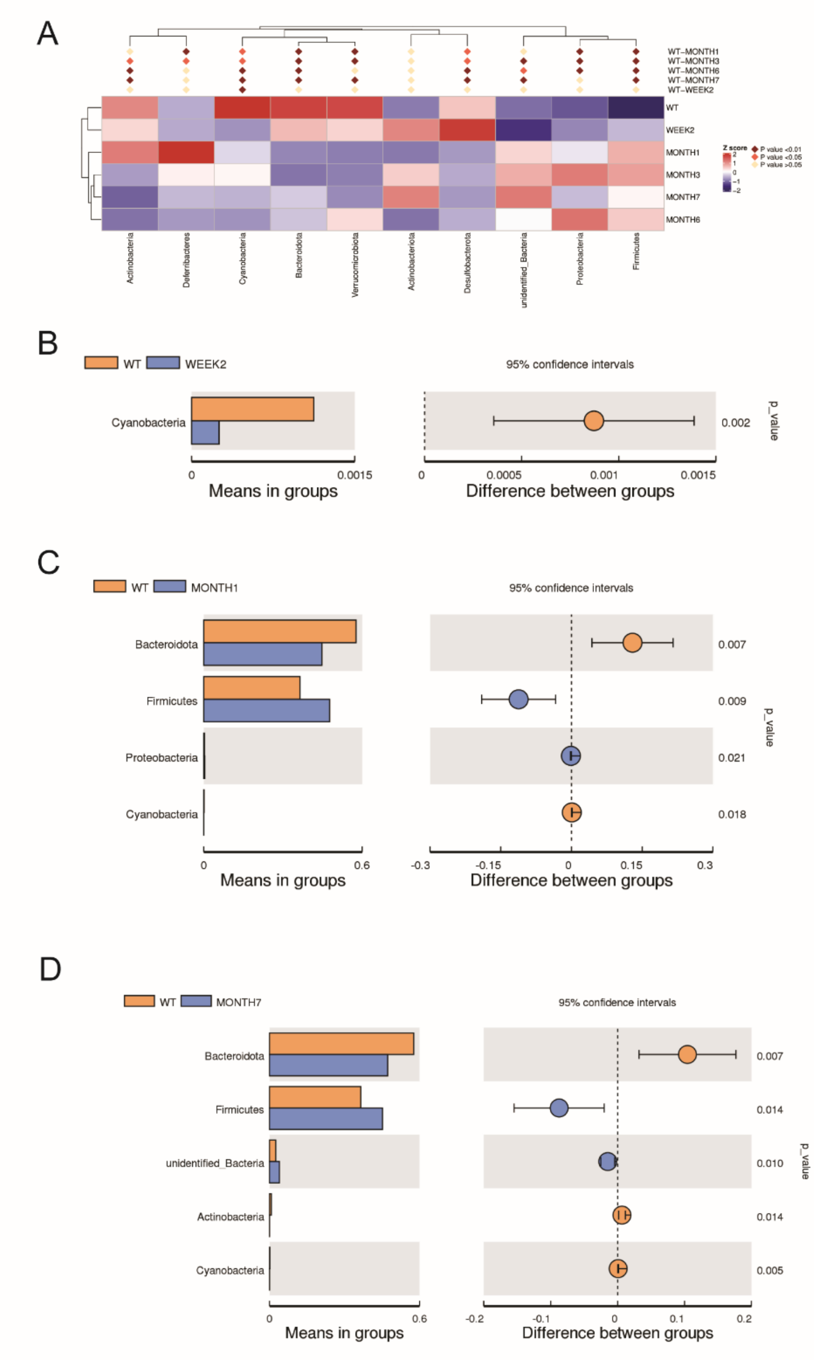


**Fig. S3. Related to Figure 2**

(**A**) Cluster distribution of phylum found in the microbiota of mice injected with monomeric αSyn or αSyn PFF in the stomach. (**B-D**) Comparison of the mean in groups (left) and difference between group (right) of significantly different Phylum between WT (monomeric αSyn) and mice injected with αSyn PFF for 2 weeks (**B**), 1 month (**C**) and 7 months (**D**). Box plots are representing the mean and standard deviation. Statistical data represent post hoc analyses compared to monomeric αSyn group following unpaired t-test.

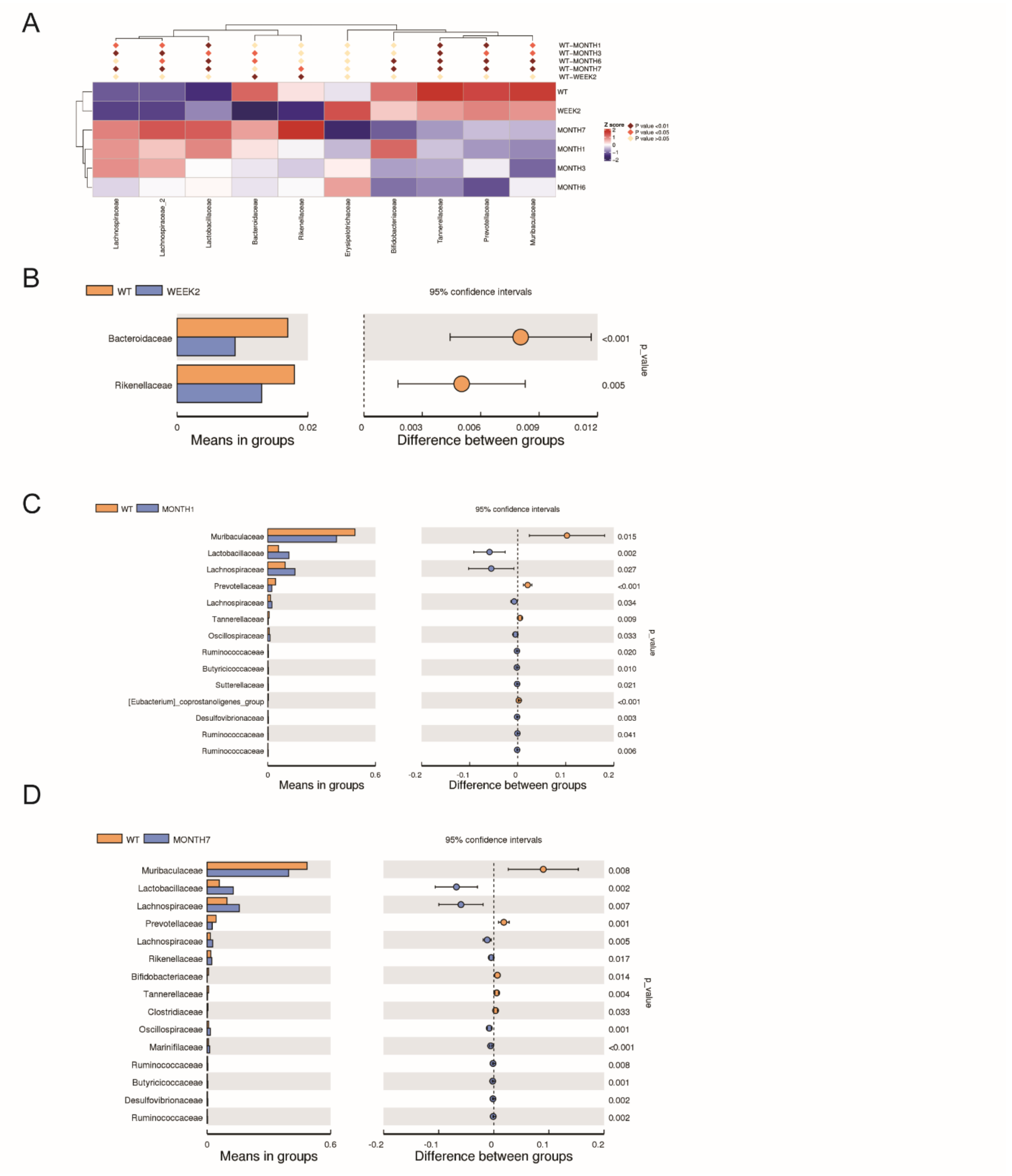


**Fig. S4. Related to Figure 2**

(**A**) Cluster distribution of family found in the microbiota of mice injected with monomeric αSyn or αSyn PFF in the stomach. (**B-D**) Comparison of the mean in groups (left) and difference between group (right) of significantly different family between WT (monomeric αSyn) and mice injected with αSyn PFF for 2 weeks (**B**), 1 month (**C**) and 7 months (**D**). Box plots are representing the mean and standard deviation. Statistical data represent post hoc analyses compared to monomeric αSyn group following unpaired t-test.

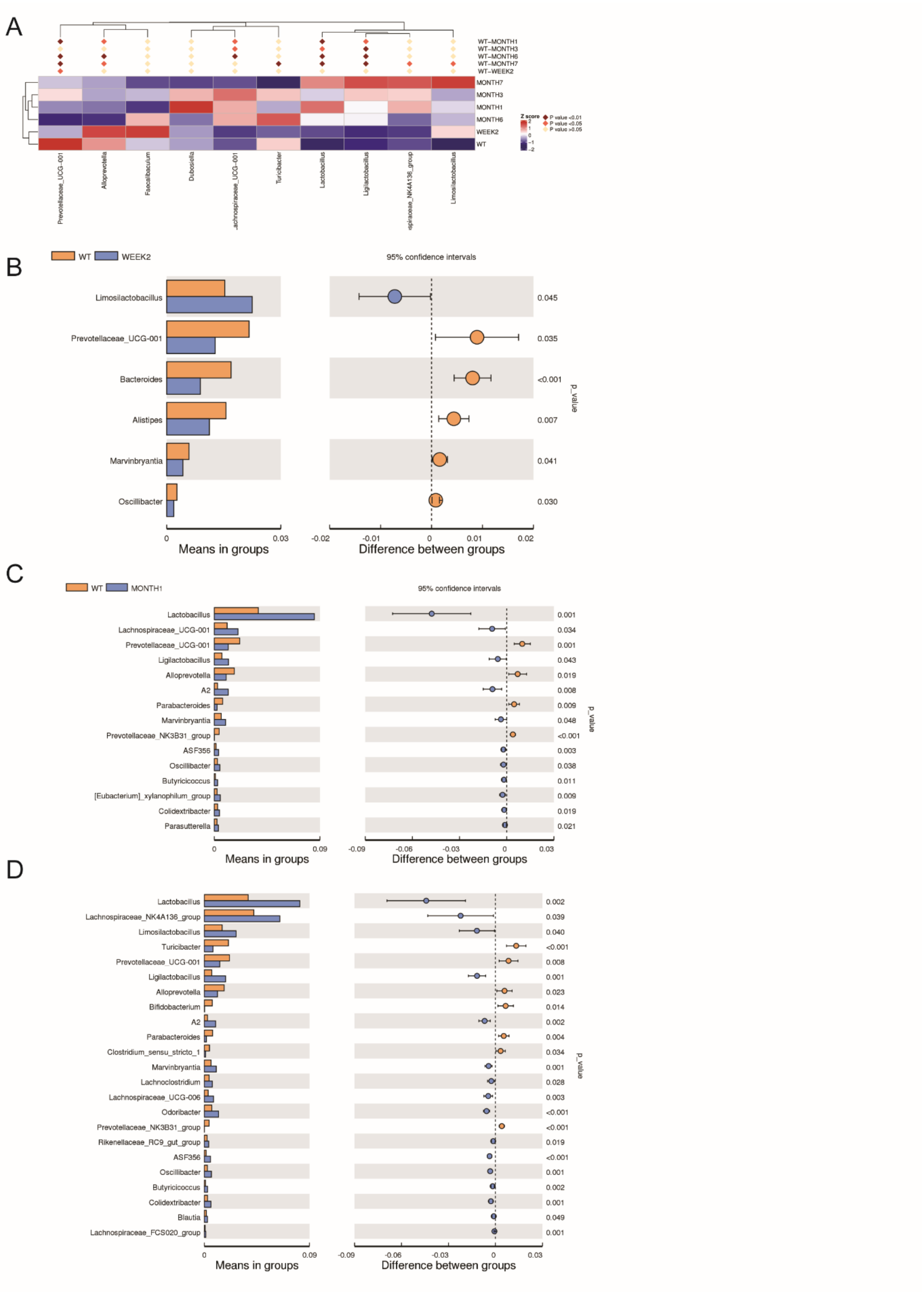

A
WT−MONTH1
WT−MONTH3
WT−MONTH6
WT−MONTH7
WT−WEEK2
MONTH7
MONTH3
MONTH1
MONTH6
WEEK2
WT
Z score
2
1
0
−1
−2
P value <0.01
P value <0.05
P value >0.05
Prevotellaceae_UCG-001
Alloprevotella
Faecalibaculum
Dubosiella
achnospiraceae_UCG-001
Turicibacter
Lactobacillus
Ligilactobacillus
spiraceae_NK4A136_group
Limosilactobacillus
B
WT
WEEK2
95% confidence intervals
Limosilactobacillus
Prevotellaceae_UCG-001
Bacteroides
Alistipes
Marvinbryantia
Oscillibacter
0.045
0.035
<0.001
0.007
0.041
0.030
p_value
0
0.03
Means in groups
-0.02
-0.01
0
0.01
0.02
Difference between groups
C
WT
MONTH1
95% confidence intervals
Lactobacillus
Lachnospiraceae_UCG-001
Prevotellaceae_UCG-001
Ligilactobacillus
Alloprevotella
A2
Parabacteroides
Marvinbryantia
Prevotellaceae_NK3B31_group
ASF356
Oscillibacter
Butyricicoccus
[Eubacterium]_xylanophilum_group
Colidextribacter
Parasutterella
0.001
0.034
0.001
0.043
0.019
0.008
0.009
0.048
<0.001
0.003
0.038
0.011
0.009
0.019
0.021
p_value
0
0.09
Means in groups
-0.09
-0.06
-0.03
0
0.03
Difference between groups
D
Lactobacillus
Lachnospiraceae_NK4A136_group
Limosilactobacillus
Turicibacter
Prevotellaceae_UCG-001
Ligilactobacillus
Alloprevotella
Bifidobacterium
A2
Parabacteroides
Clostridium_sensu_stricto_1
Marvinbryantia
Lachnoclostridium
Lachnospiraceae_UCG-006
Odoribacter
Prevotellaceae_NK3B31_group
Rikenellaceae_RC9_gut_group
ASF356
Oscillibacter
Butyricicoccus
Colidextribacter
Blautia
Lachnospiraceae_FCS020_group
0.002
0.039
0.040
<0.001
0.008
0.001
0.023
0.014
0.002
0.004
0.034
0.001
0.028
0.003
<0.001
<0.001
0.019
<0.001
0.001
0.002
0.001
0.049
0.001
p_value
0
0.09
Means in groups
-0.09
-0.06
-0.03
0
0.03
Difference between groups

**Fig. S5. Related to Figure 2**

(**A**) Cluster distribution of genus found in the microbiota of mice injected with monomeric αSyn or αSyn PFF in the stomach. (**B-D**) Comparison of the mean in groups (left) and difference between group (right) of significantly different genus between WT (monomeric αSyn) and mice injected with αSyn PFF for 2 weeks (**B**), 1 month (**C**) and 7 months (**D**). Box plots are representing the mean and standard deviation. Statistical data represent post hoc analyses compared to monomeric αSyn group following unpaired t-test.

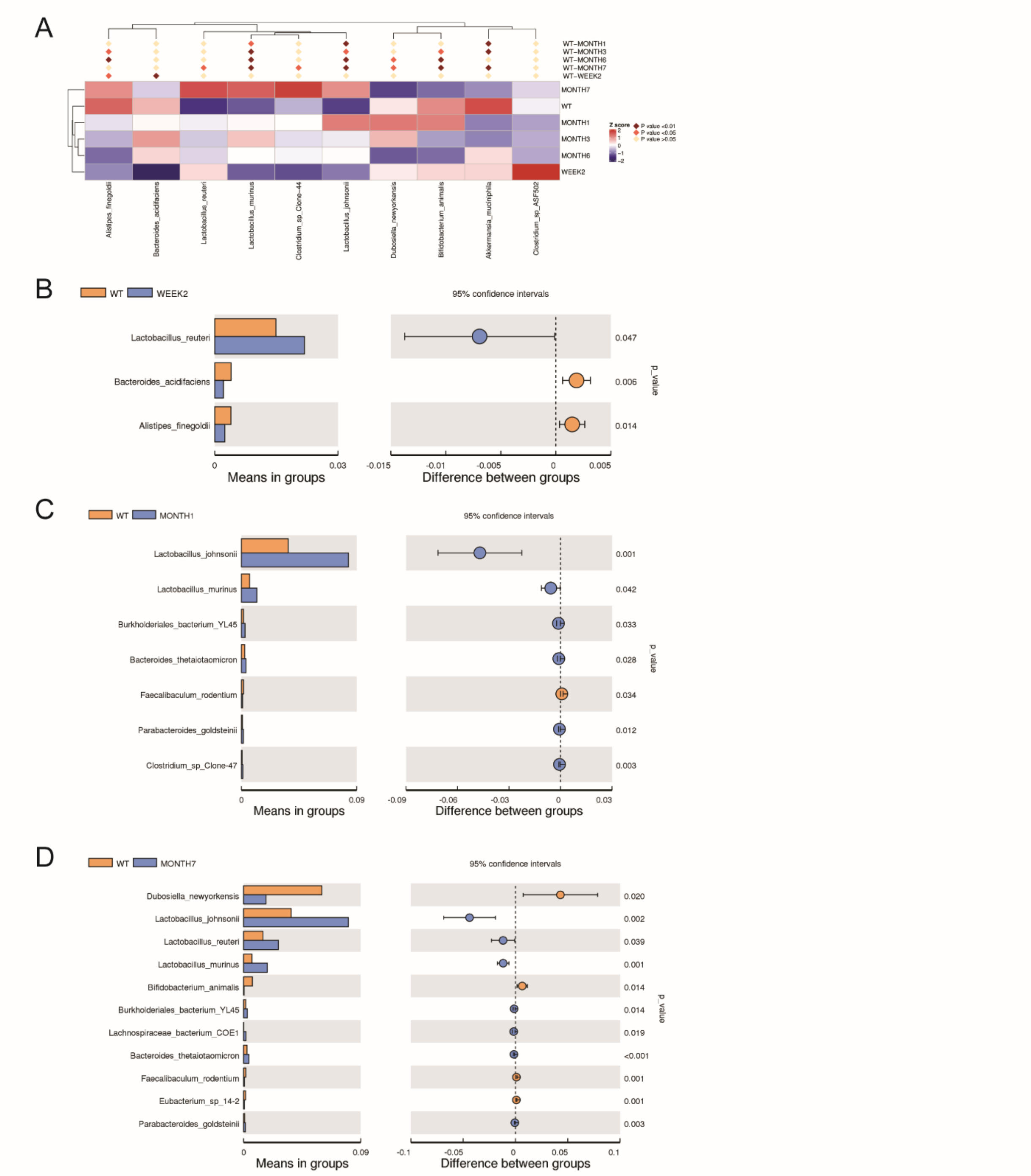


**Fig. S6. Related to Figure 2**

(**A**) Cluster distribution of species found in the microbiota of mice injected with monomeric αSyn or αSyn PFF in the stomach. (**B-D**) Comparison of the mean in groups (left) and difference between group (right) of significantly different species between WT (monomeric αSyn) and mice injected with αSyn PFF for 2 weeks (**B**), 1 month (**C**) and 7 months (**D**). Box plots are representing the mean and standard deviation. Statistical data represent post hoc analyses compared to monomeric αSyn group following unpaired t-test.

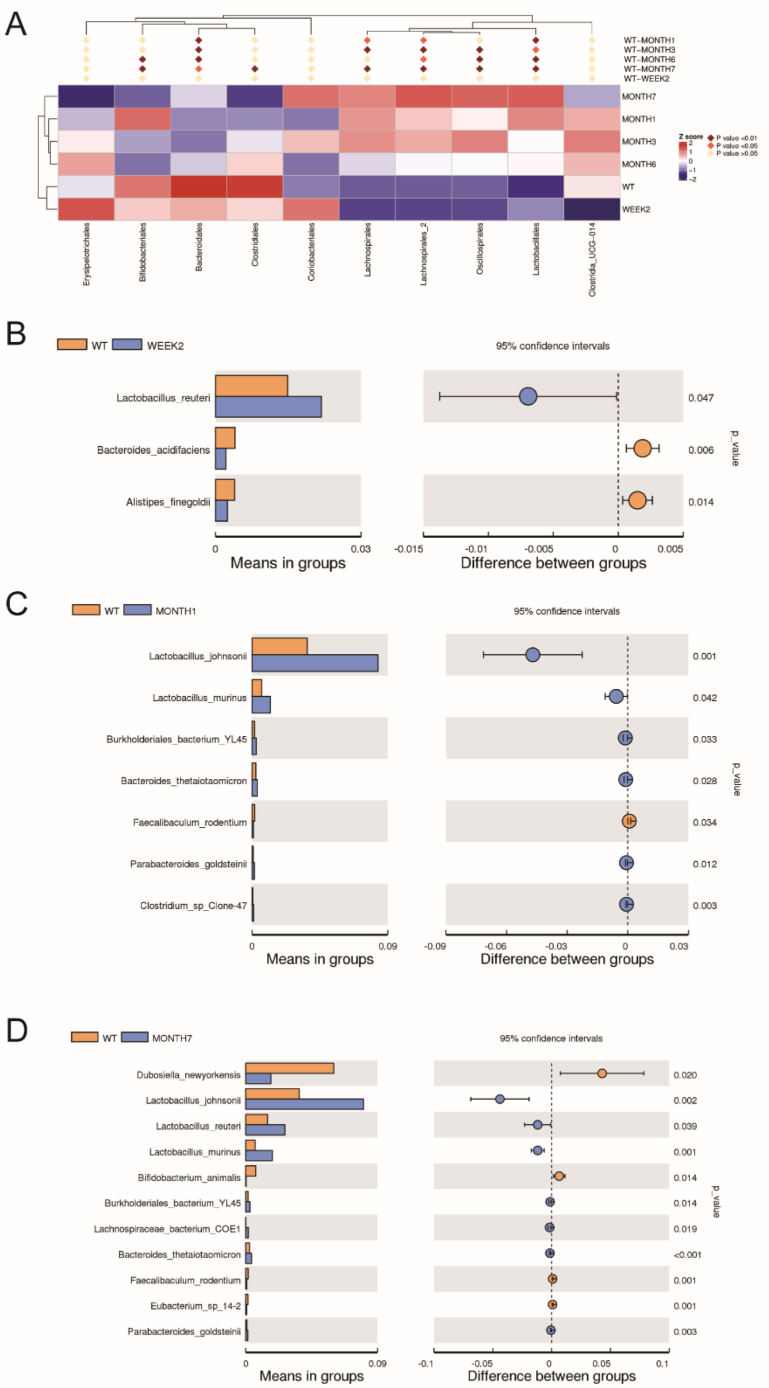

**Fig. S7. Related to Figure 2**

(**A**) Cluster distribution of orders found in the microbiota of mice injected with monomeric αSyn or αSyn PFF in the stomach. (**B-D**) Comparison of the mean in groups (left) and difference between group (right) of significantly different orders between WT (monomeric αSyn) and mice injected with αSyn PFF for 2 weeks (**B**), 1 month (**C**) and 7 months (**D**). Box plots are representing the mean and standard deviation. Statistical data represent post hoc analyses compared to monomeric αSyn group following unpaired t-test.

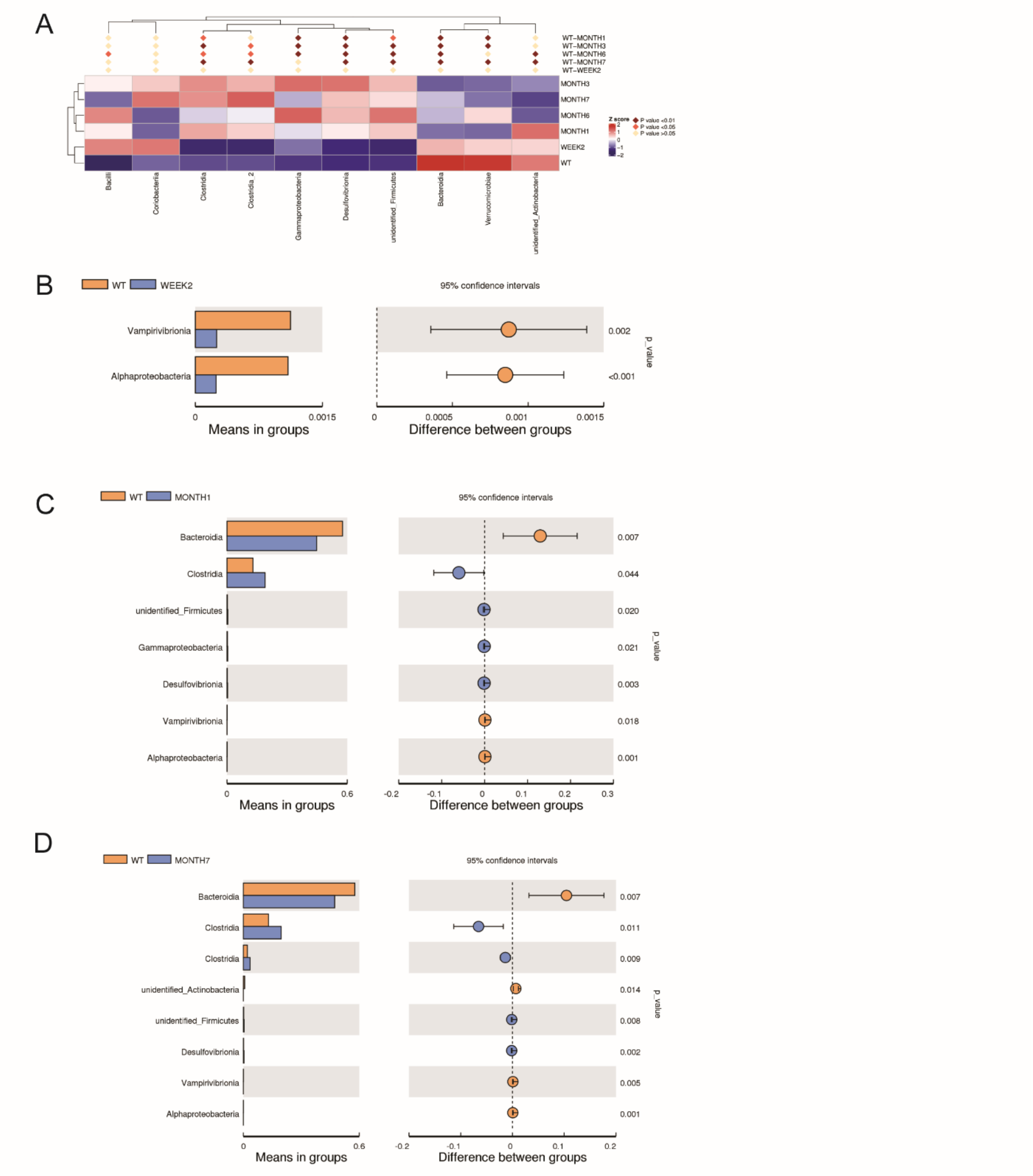


**Fig. S8. Related to Figure 2**

(**A**) Cluster distribution of class found in the microbiota of mice injected with monomeric αSyn or αSyn PFF in the stomach. (**B-D**) Comparison of the mean in groups (left) and difference between group (right) of significantly different class between WT (monomeric αSyn) and mice injected with αSyn PFF for 2 weeks (**B**), 1 month (**C**) and 7 months (**D**). Box plots are representing the mean and standard deviation. Statistical data represent post hoc analyses compared to monomeric αSyn group following unpaired t-test.

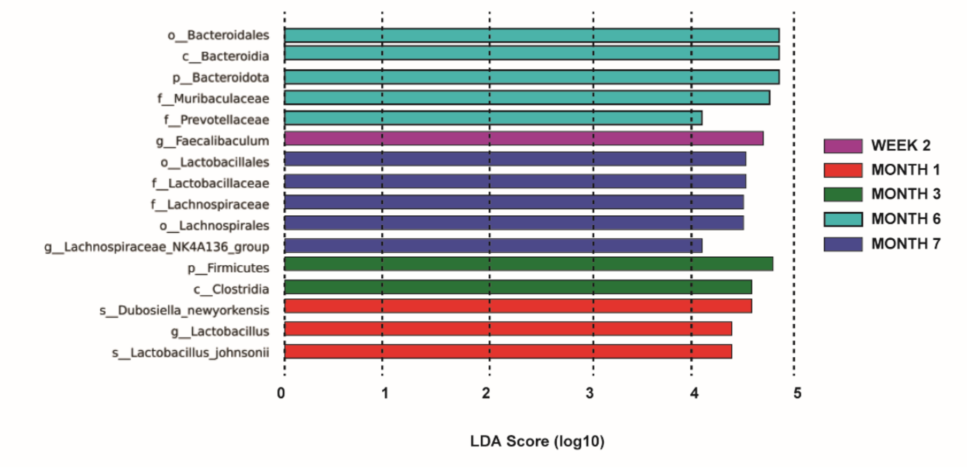


**Fig. S9. Related to Figure 2.**

Linear discriminant analysis of the effect size class found in the microbiota of mice injected with monomeric αSyn or αSyn PFF in the stomach.

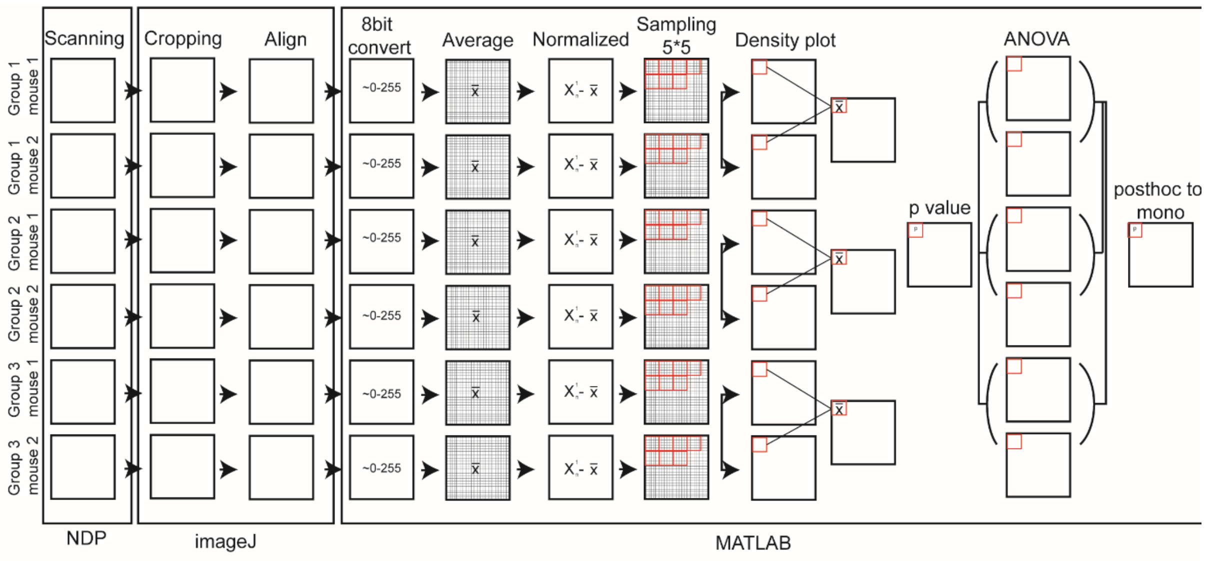

**Fig. S10. Related to Figure 3.**

Graphical illustration of the analyses of single and group images obtained for pSer129-DAB staining.

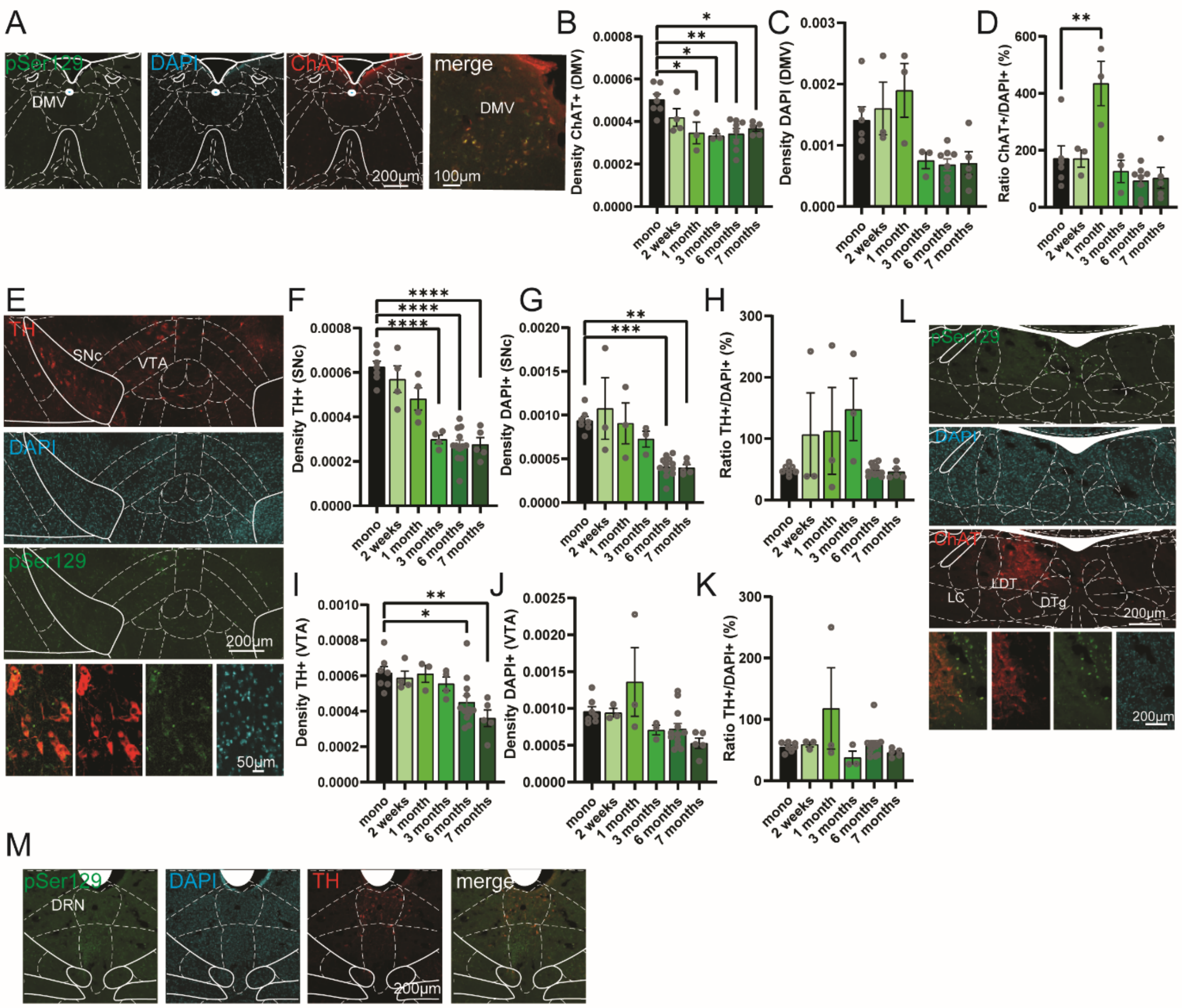


**Fig. S11. Related to Figure 3.**

(**A**) Confocal images of pSer129, DAPI and ChAT staining in the Dorsal Vagus Nerve of mice injected with αSyn PFF for 7 months. DMV: Dorsal Vagus nerve. (**B**) Quantification of the density of ChAT-positive neurons found in the DMV of mice injected with monomeric or a Syn PFF in the stomach ($F_{5,24}$=5.968, P=0.0010). (**C**) Quantification of the density of DAPI-positive neurons found in the DMV of mice injected with monomeric αSyn or αSyn PFF in the stomach ($F_{5,22}$=4.958, P=0.0034). (**D**) Quantification of the ratio between DAPI-positive and ChAT-positive neurons found in the DMV of mice injected with monomeric αSyn or αSyn PFF in the stomach ($F_{5,22}$=8.012, P=0.0002). (**E**) Confocal images of pSer129, DAPI and TH staining in the midbrain of mice injected with αSyn PFF for 7 months. SNc: Substantia nigra pars compacta. VTA: Ventral tegmental area. (**F**) Quantification of the density of TH-positive neurons found in the SNc of mice injected with monomeric αSyn or αSyn PFF in the stomach ($F_{5,30}$=25.80, P<0.0001). (**G**) Quantification of the density of DAPI-positive neurons found in the SNc of mice injected with monomeric αSyn or αSyn PFF in the stomach ($F_{5,27}$=9.115, P<0.0001). (**H**) Quantification of the ratio between DAPI-positive and TH-positive neurons found in the SNc of mice injected with monomeric αSyn or αSyn PFF in the stomach ($F_{5,27}$=2.785, P=0.0374). (**I**) Quantification of the density of TH-positive neurons found in the VTA of mice injected with monomeric αSyn or αSyn PFF in the stomach ($F_{5,29}$=5.095, P=0.0018). (**J**) Quantification of the density of DAPI-positive neurons found in the VTA of mice injected with monomeric αSyn or αSyn PFF in the stomach ($F_{5,27}$=3.689, P=0.0113). (**K**) Quantification of the ratio between DAPI-positive and TH-positive neurons found in the VTA of mice injected with monomeric αSyn or αSyn PFF in the stomach ($F_{5,27}$=2.086, P=0.0982). (**L**) Confocal images of pSer129, DAPI and ChAT staining in the brainstem of mice injected with αSyn PFF for 7 months. LDT: Laterodorsal tegmental nucleus. LC: Locus coeruleus. DTg: Dorsal tegmentum. (**M**) Confocal images of pSer129, DAPI and TH staining in the brainstem of mice injected with αSyn PFF for 7 months. DRN: Dorsal Raphe nucleus. Data are expressed as mean±SEM. All individual point represents individual animals. * P<0.05, ** P<0.01, *** P<0.001, **** P<0.0001. Statistical data represent post hoc analyses compared to monomeric αSyn group following 1-way ANOVA.

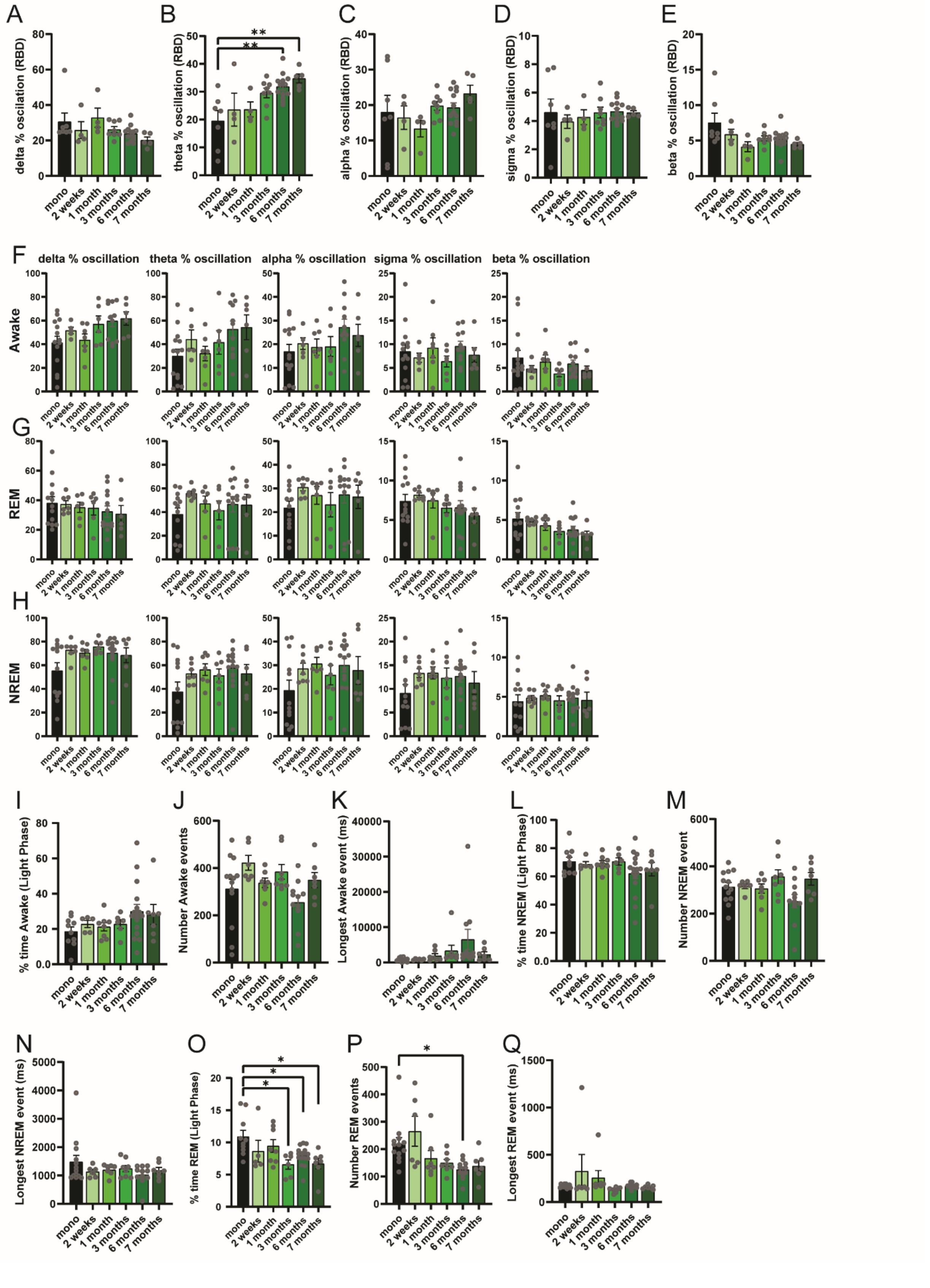

A
delta % oscillation (RBD)
B
theta % oscillation (RBD)
**
**
C
alpha % oscillation (RBD)
D
sigma % oscillation (RBD)
E
beta % oscillation (RBD)
F
delta % oscillation
theta % oscillation
alpha % oscillation
sigma % oscillation
beta % oscillation
Awake
G
REM
H
NREM
I
% time Awake (Light Phase)
J
Number Awake events
K
Longest Awake event (ms)
L
% time NREM (Light Phase)
M
Number NREM event
N
Longest NREM event (ms)
O
% time REM (Light Phase)
*
*
*
P
Number REM events
*
Q
Longest REM event (ms)
mono
2 weeks
1 month
3 months
6 months
7 months

**Fig. S12. Related to Figure 4.**

(**A**) Delta oscillation of the cortical periodogram express as percentage of all oscillation during RBD events $F_{5,35}$=1.905, P=0.1185. (**B**) Theta oscillation of the cortical periodogram express as percentage of all oscillation during RBD events $F_{5,35}$=4.988, P=0.0015. (**C**) Alpha oscillation of the cortical periodogram express as percentage of all oscillation during RBD events $F_{5,35}$=1.102, P=0.3767. (**D**) Sigma oscillation of the cortical periodogram express as percentage of all oscillation during RBD events $F_{5,35}$=0.2403, P=0.9419. (**E**) Beta oscillation of the cortical periodogram express as percentage of all oscillation during RBD events $F_{5,35}$=2.541, P=0.0460. (**F**) Cortical periodogram oscillation (delta, theta, alpha, sigma and beta) express as percentage of all oscillation during awake events ($F_{5,43}$=2.554, P=0.0414; $F_{5,43}$=2.046, P=0.0912; $F_{5,43}$=1.394, P=0.2455; $F_{5,43}$=0.5422, p=0.7432; $F_{5,43}$=0.8671, P=0.5110, respectively). (**G**) Cortical periodogram oscillation (delta, theta, alpha, sigma and beta) express as percentage of all oscillation during REM events ($F_{5,45}$=0.4386, P=0.8192; $F_{5,45}$=0.8534, P=0.5196; $F_{5,45}$=0.7783, P=0.5706; $F_{5,45}$=0.8458, P=0.5246; $F_{5,45}$=1.655, P=0.1651, respectively). (**H**) Cortical periodogram oscillation (delta, theta, alpha, sigma and beta) express as percentage of all oscillation during NREM events ($F_{5,47}$=2.224, P=0.0674; $F_{5,47}$=1.616, P=0.1743; $F_{5,47}$=1.428, P=0.2317; $F_{5,47}$=1.084, P=0.3818; $F_{5,47}$=0.1677, P=0.9732, respectively). (**I**) Percentage of time awake during the light phase (0700-1900) $F_{5,46}$=1.509, P=0.2057. (**J**) Total number of awake events over the 24h recordings. $F_{5,46}$=2.682, P=0.0329. (**K**) Duration of the longest awake events $F_{5,46}$=2.041, P=0.0904. (**L**) Percentage of time in NREM during the light phase (0700-1900) $F_{5,46}$=1.083, P=0.3825. (**M**) Total number of NREM events over the 24h recordings $F_{5,46}$=2.469, P=0.0461. (**N**) Duration of the longest NREM events $F_{5,46}$=1.116, P=0.3654. (**O**) Percentage of time in REM during the light phase (0700-1900) $F_{5,46}$=3.863, P=0.0053. (**P**) Total number of REM events over the 24h recordings $F_{5,44}$=5.251,

P=0.0007. (**Q**) Duration of the longest REM events $F_{5,46}$=1.494, P=0.2101. Data are expressed as mean±SEM. All individual point represents individual animals. * P<0.05, ** P<0.01. Statistical data represent post hoc analyses compared to monomeric αSyn group following 1-way ANOVA.

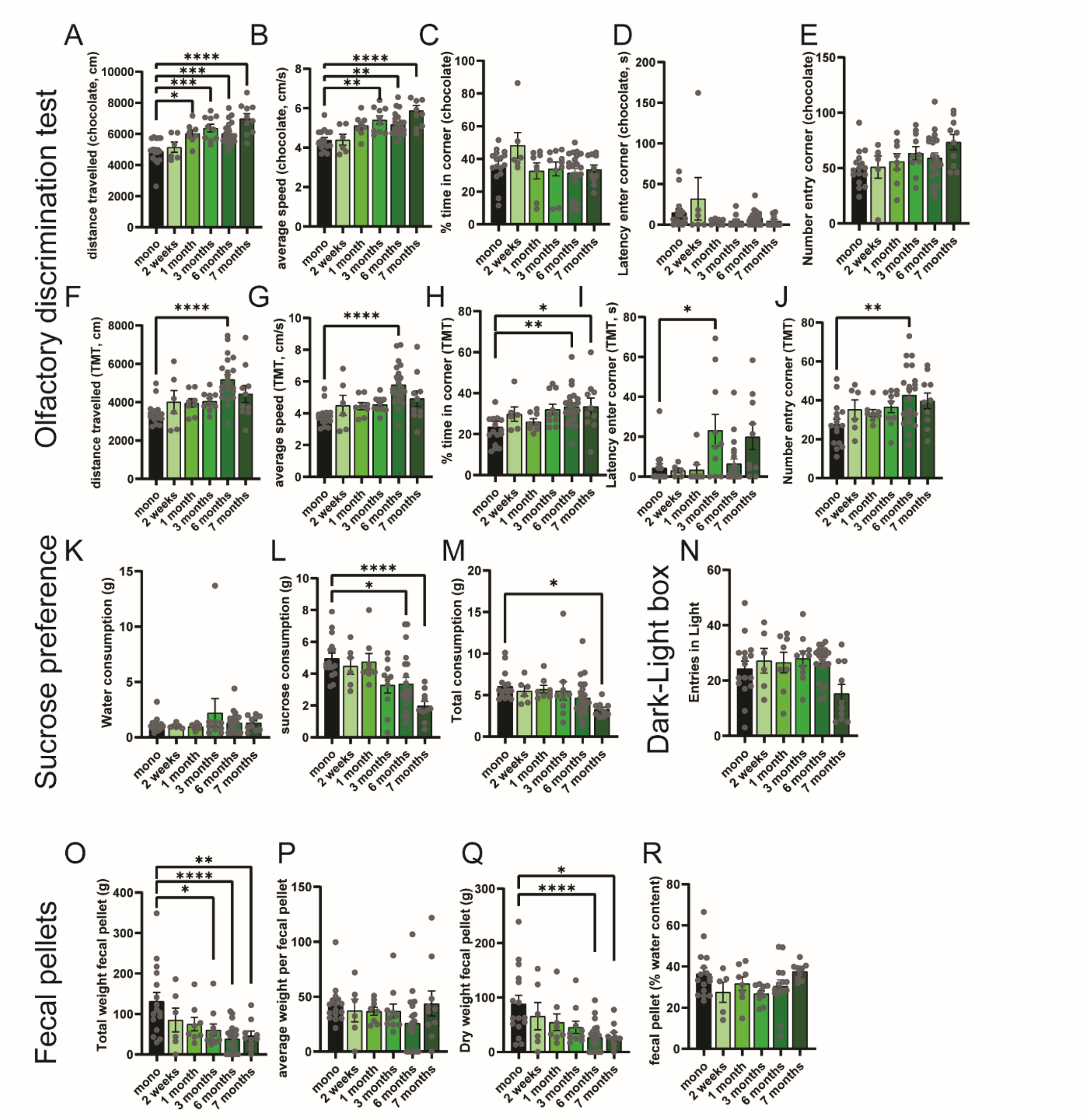


**Fig. S13. Related to Figure 4.**

(**A**) Distance travelled during the olfactory test paired with chocolate smell $F_{5,65}$=11.02, P<0.0001. (**B**) Average speed during the olfactory test paired with chocolate smell $F_{5,65}$=9.207, P<0.0001. (**C**) Percentage of time spent in the corner paired with chocolate smell $F_{5,65}$=1.741, P=0.1378. (**D**) Latency to first entry in the corner paired with chocolate smell $F_{5,65}$=1.963, P=0.0960. (**E**) Total number of entries in the corner paired with chocolate smell $F_{5,65}$=2.061, P=0.0817. (**F**) Distance travelled during the olfactory test paired with TMT smell $F_{5,65}$=6.266, P<0.0001. (**G**) Average speed during the olfactory test paired with TMT smell $F_{5,65}$=6.312, P<0.0001. (**H**) Percentage of time spent in the corner paired with TMT smell $F_{5,64}$=4.096, P=0.0027. (**I**) Latency to first entry in the corner paired with TMT smell $F_{5,65}$=4.098, P=0.0027. (**J**) Total number of entries in the corner paired with TMT smell $F_{5,65}$=3.070, P=0.0150. (**K**) Total water consumptions over the 24h test $F_{5,65}$=0.7661, P=0.5776. (**L**) Total sucrose consumptions over the 24h test $F_{5,65}$=6.397, P<0.0001. (**M**) Total consumptions over the 24h test $F_{5,65}$=2.550, P=0.0361. **(N)** Total number of entries in the light zone during the dark-light test $F_{5,65}$=2.601, P=0.0331. (**O**) Total weight of the fecal pellets produced in 20 min $F_{5,65}$=5.637, P=0.0002. (**P**) Average weight per individual pellets produced in 20min $F_{5,65}$=1.225, P=0.3077. (**Q**) Weight of the dry content of fecal pellet produced 20min $F_{5,65}$=4.366, P=0.0017. (**R**) Percentage of water content in fecal pellets produced in 20min $F_{5,65}$=2.150, P=0.0728. Data are expressed as mean±SEM. All individual point represents individual animals. * P<0.05, ** P<0.01, *** P<0.001, **** P<0.0001. Statistical data represent post hoc analyses compared to monomeric αSyn group following 1-way ANOVA.

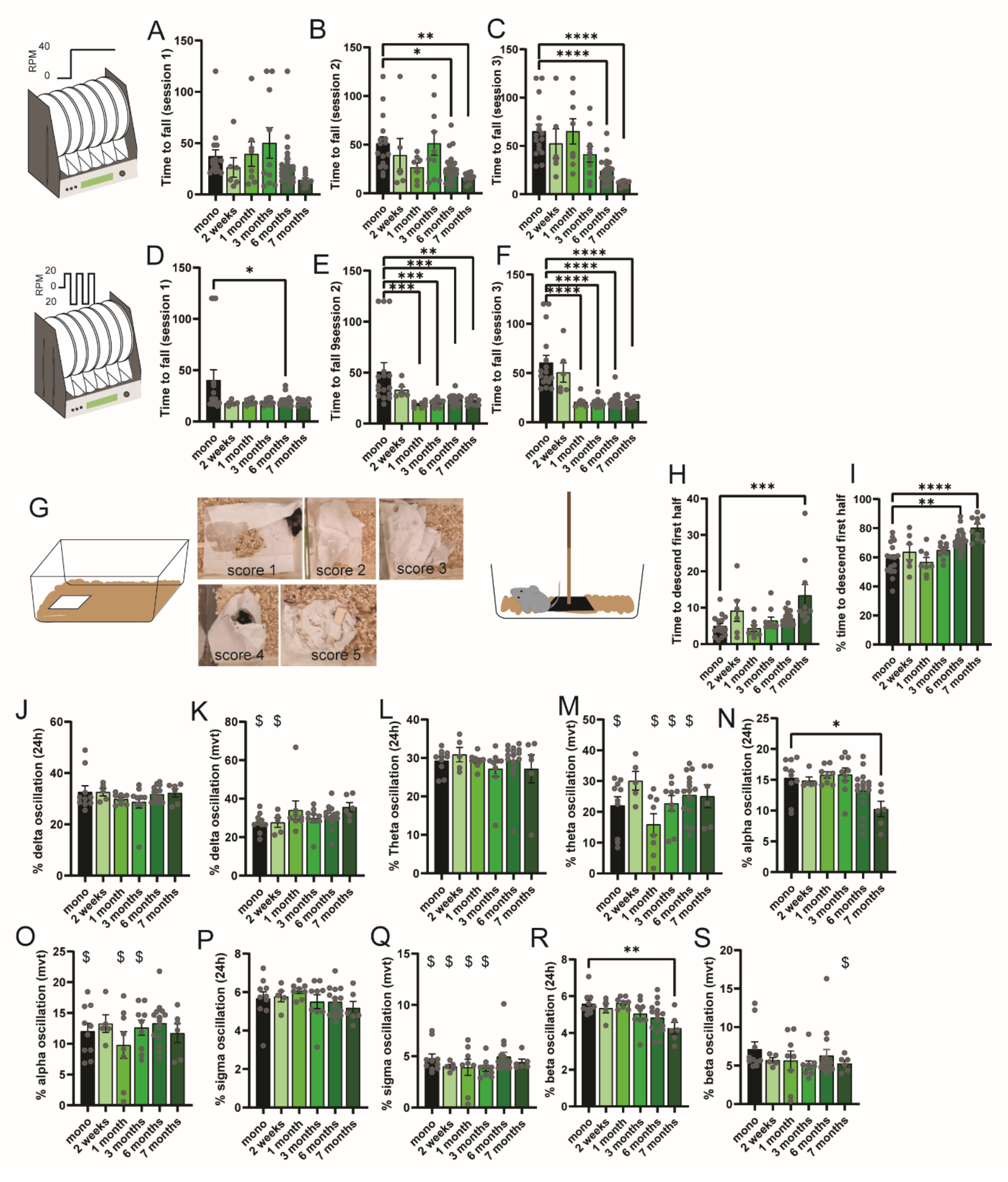


**Fig. S14. Related to Figure 5.**

(**A**) Time to fall on the constant speed 40 RPM rotarod schedule during first session $F_{5,65}$=1.948, P=0.0984. (**B**) Time to fall on the constant speed 40 RPM rotarod schedule during second session

$F_{5,65}$=4.106, P=0.0027. (**C**) Time to fall on the constant speed 40 RPM rotarod schedule during third session $F_{5,65}$=10.18, P<0.0001. (**D**) Time to fall on the docking speed 20 RPM rotarod schedule during first session $F_{5,65}$=2.946, P=0.0185. (**E**) Time to fall on the docking speed 20 RPM rotarod schedule during second session $F_{5,65}$=6.887, P<0.0001. (**F**) Time to fall on the docking speed 20 RPM rotarod schedule during third session $F_{5,65}$=15.49, P<0.0001. (**G**) Representative images of the scoring allocated during the bedding test. (**H**) Time to descend the first half of the pole during the pole test $F_{5,65}$=5.279, P=0.0004. (**I**) Percentage of time to descend the first half of the pole test compared to the entire pole test $F_{5,65}$=10.41, P<0.0001. (**J**) Delta oscillation of the cortical periodogram express as percentage of all oscillation during 24h recordings $F_{5,47}$=0.8781, P=0.5031. (**K**) Delta oscillation of the cortical periodogram express as percentage of all oscillation during activity events $F_{5,47}$=1.426, P=0.2322. (**L**) Theta oscillation of the cortical periodogram express as percentage of all oscillation during 24h recordings $F_{5,47}$=0.5092, P=0.7678. (**M**) Theta oscillation of the cortical periodogram express as percentage of all oscillation during activity events $F_{5,47}$=2.279, P=0.0618. (**N**) Alpha oscillation of the cortical periodogram express as percentage of all oscillation 24h recordings $F_{5,47}$=3.835, P=0.0054. (**O**) Alpha oscillation of the cortical periodogram express as percentage of all oscillation during activity events $F_{5,47}$=0.8813, P=0.5011. (**P**) Sigma oscillation of the cortical periodogram express as percentage of all oscillation during 24h recordings $F_{5,47}$=0.9291, P=0.4707. (**Q**) Sigma oscillation of the cortical periodogram express as percentage of all oscillation during activity events $F_{5,47}$=1.097, P=0.3745. (**R**) Beta oscillation of the cortical periodogram express as percentage of all oscillation during 24h recordings $F_{5,47}$=4.219, P=0.0030. (**S**) Beta oscillation of the cortical periodogram express as percentage of all oscillation during activity events $F_{5,47}$=0.7363, P=0.6000. Data are expressed as mean±SEM. All individual point represents individual animals. * P<0.05, ** P<0.01, ***

P<0.001, **** P<0.0001. Statistical data represent post hoc analyses compared to monomeric αSyn group following 1-way ANOVA. $ are used for P<0.05 of the paired t-test between group during 24h and activity events.

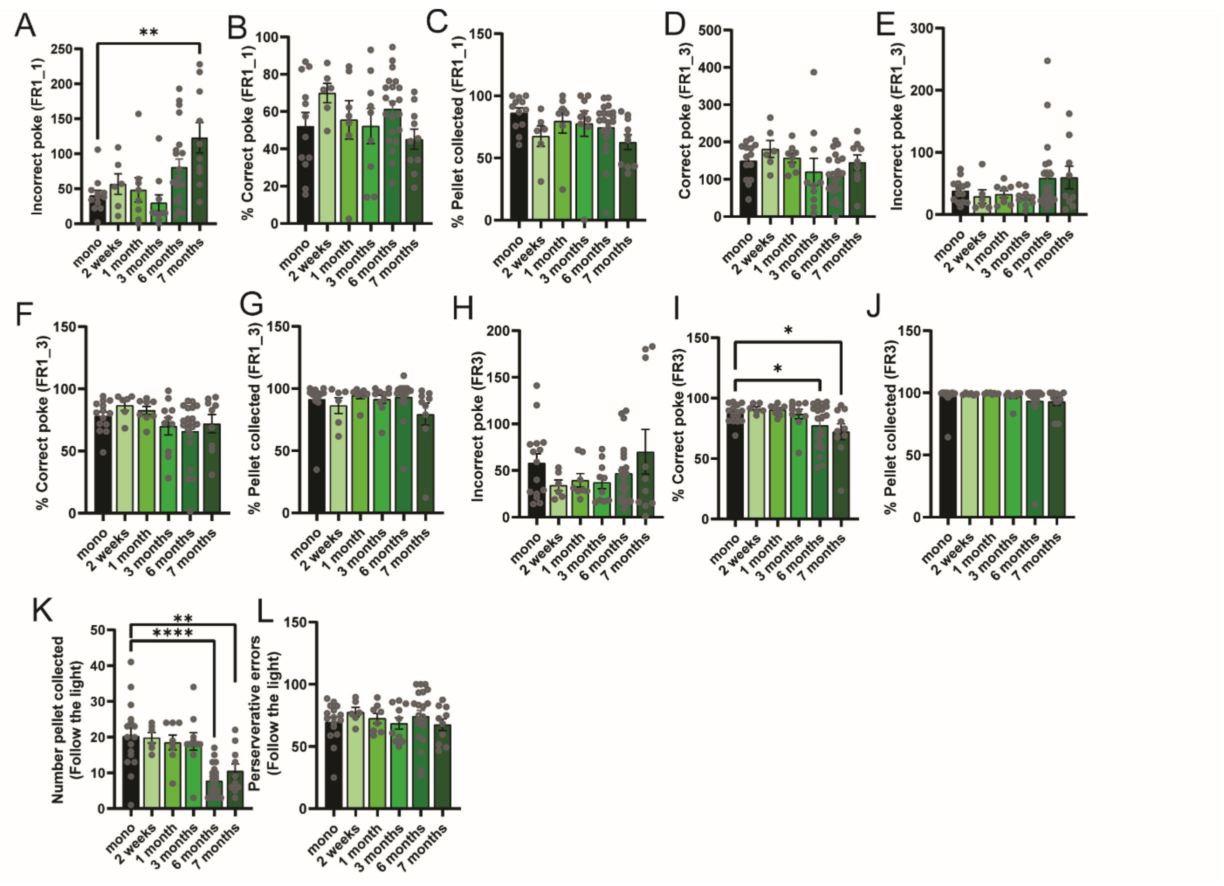


**Fig. S15. Related to Figure 5.**

(**A**) Number of incorrect nose poke in the first session of the fixed ratio 1 $F_{5,61}$=5.064, P=0.0006. (**B**) Percentage of correct poke in the first session of the fixed ratio 1 $F_{5,59}$=1.354, P=0.2546. (**C**) Percentage of pellets collected within 10s in the first session of the fixed ratio 1 $F_{5,59}$=1.518, P=0.1981. (**D**) Number of correct pokes in the third session of the fixed ratio 1 $F_{5,62}$=1.724, P=0.1423. (**E**) Number of incorrect pokes in the third session of the fixed ratio 1 $F_{5,62}$=1.465, P=0.2141. (**F**) Percentage of correct poke in the third session of the fixed ratio 1 $F_{5,62}$=1.826,

P=0.1209. (**G**) Percentage of pellets collected within 10s in the third session of the fixed ratio 1 $F_{5,62}$=1.137, P=0.3507. (**H**) Number of incorrect pokes in the first session of the fixed ratio 3 $F_{5,64}$=1.157, P=0.3401. (**I**) Percentage of correct poke in the first session of the fixed ratio 3 $F_{5,64}$=3.032, P=0.0162 (post hoc is LSD test). (**J**) Percentage of pellets collected within 10s in the first session of the fixed ratio 3 $F_{5,64}$=0.5086, P=0.7687. (**K**) Total number of pellets collected within 10s in the "follow the light" protocol $F_{5,65}$=9.238, P<0.0001. (**L**) Number of perseverative errors in the "follow the light" protocol $F_{5,65}$=0.5002, P=0.7750. Data are expressed as mean±SEM. All individual point represents individual animals. * P<0.05, ** P<0.01, **** P<0.0001. Statistical data represent post hoc analyses compared to monomeric αSyn group following 1-way ANOVA.

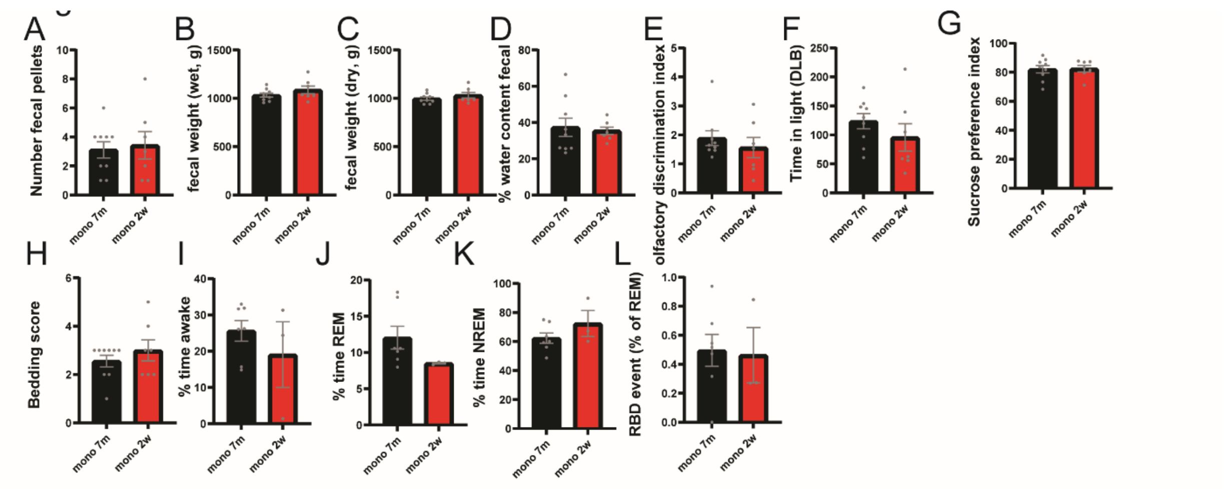


**Fig. S16. Related to Figure 3-5.**

(**A**) Number of fecal pellets in mice injected with monomeric αSyn for 7 months (black) or 2 weeks (red) (unpaired t-test $t_{14}$=0.3028, P=0.7665). (**B**) Total weight of fecal pellets in mice injected with monomeric αSyn for 7 months (black) or 2 weeks (red) (unpaired t-test $t_{14}$=1.310, P=0.2113). (**C**) Total dry fecal weight in mice injected with monomeric αSyn for 7 months (black) or 2 weeks (red) (unpaired t-test $t_{14}$=1.113, P=0.843). (**D**) Percentage of water content in fecal pellets in mice injected with monomeric αSyn for 7 months (black) or 2 weeks (red) (unpaired t-test $t_{14}$=0.3210,

P=0.7529). (**E**) Olfactory discrimination index in mice injected with monomeric αSyn for 7 months (black) or 2 weeks (red) (unpaired t-test $t_{14}$=0.7650, P=0.4570). (**F**) Time spent in the light area of the dark light box in mice injected with monomeric αSyn for 7 months (black) or 2 weeks (red) (unpaired t-test $t_{14}$=1.095, P=0.2919). (**G**) Sucrose preference index in mice injected with monomeric αSyn for 7 months (black) or 2 weeks (red) (unpaired t-test $t_{14}$=0.1247, P=0.9025). (**H**) Bedding score in mice injected with monomeric αSyn for 7 months (black) or 2 weeks (red) (unpaired t-test $t_{14}$=0.9439, P=0.3612). (**I**) Percentage of time awake in mice injected with monomeric αSyn for 7 months (black) or 2 weeks (red) (unpaired t-test $t_8$=0.9321, P=0.3786). (**J**) Percentage in REM in mice injected with monomeric αSyn for 7 months (black) or 2 weeks (red) (unpaired t-test $t_8$=1.414, P=0.1951). (**K**) Percentage in NREM in mice injected with monomeric αSyn for 7 months (black) or 2 weeks (red) (unpaired t-test $t_8$=1.298, P=0.2303). (**L**) Percentage of RBD events over the number of REM events in mice injected with monomeric αSyn for 7 months (black) or 2 weeks (red) (unpaired t-test $t_8$=0.1572, P=0.8790). Data are expressed as mean±SEM. All individual point represents individual animals. Statistical data represent unpaired t-test analyses.

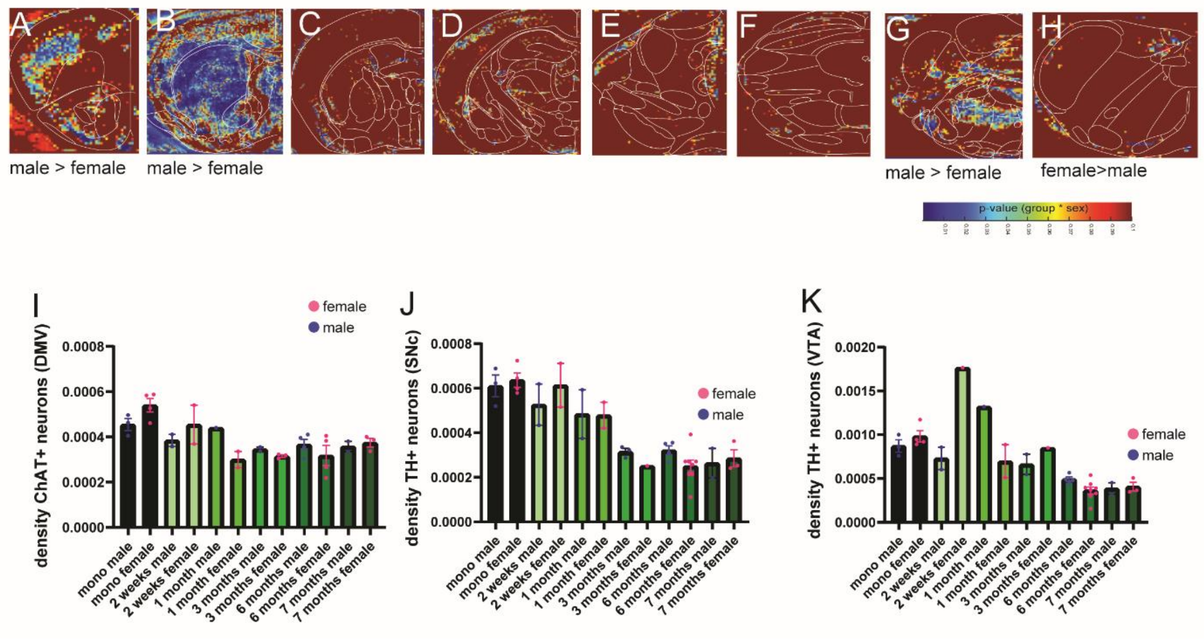


**Fig. S17. Related to Figure 3-5.**

2-way ANOVA comparison of the intensity of pSer129-DAB staining in ROIs between male and female (sex x group effects) in the frontal cortex (**A**), the striatum (**B**), the forebrain (**C**), the thalamus (**D**), the midbrain (**E**), the brainstem (**F**) the Pons (**G**) and the medulla (**H**). (**I**) Density of ChAT-positive neurons in the DMV of male and female mice injected with monomeric or αSyn PFF in the stomach (group $F_{5,18}$=5.8739, P=0.002177, sex $F_{1,18}$=0.0141, P=0.9069, interaction $F_{5,18}$=1.885, P=0.147). (**J**) Density of TH-positive neurons in the SNc of male and female mice injected with monomeric αSyn or αSyn PFF in the stomach (group $F_{5,24}$=21.94, P<0.00001, sex $F_{1,24}$=0.0005, P=0.9815, interaction $F_{5,24}$=0.7495, P=0.5944). (**K**) Density of TH-positive neurons in the VTA of male and female mice injected with monomeric αSyn or αSyn PFF in the stomach (group $F_{5,23}$=4.2027, P=0.007362, sex $F_{1,23}$=0.2384, P=0.629, interaction $F_{5,23}$=0.4132, P=0.836).

Data are expressed as mean±SEM.  All individual point represents individual animals. * P<0.05, ** P<0.01, *** P<0.001, **** P<0.0001. Statistical data represent post hoc analyses compared to monomeric αSyn group following 2-way ANOVA.

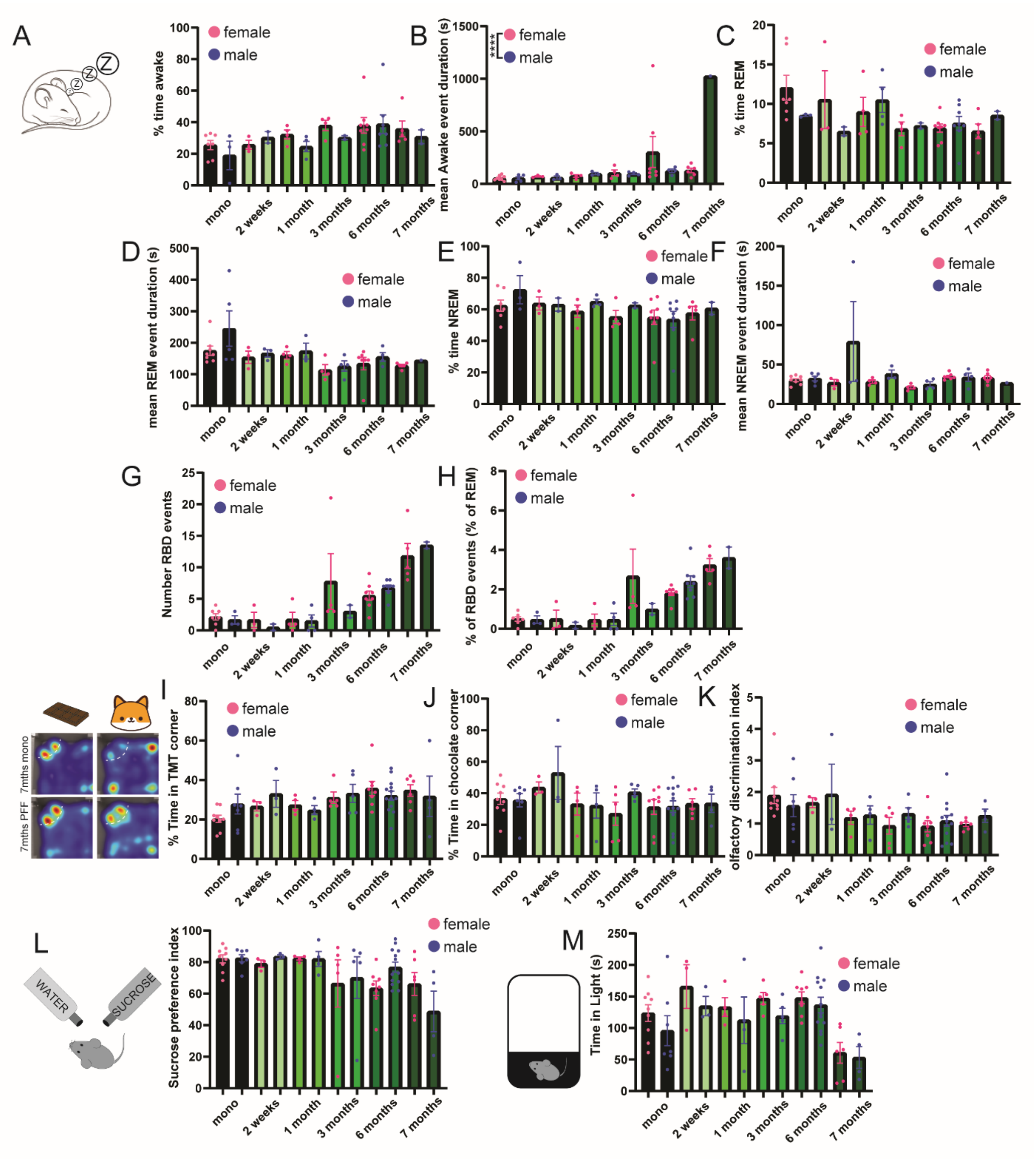


**Fig. S18. Related to Figure 4.**

(**A**) Percentage of time awake in male and female mice injected with monomeric αSyn or αSyn PFF in the stomach (group $F_{5,40}$=2.60, P=0.039299, sex $F_{1,40}$=1.027, P=0.3167, interaction $F_{5,40}$=0.36, P=0.868847). (**B**) Mean wake duration in male and female mice injected with monomeric αSyn or αSyn PFF in the stomach (group $F_{5,40}$=7.27, P=0.000063, sex $F_{1,40}$=5.98, P=0.018934, interaction $F_{5,40}$=6.43, P=0.00018). (**C**) Percentage of time REM in male and female mice injected with monomeric αSyn or αSyn PFF in the stomach (group $F_{5,40}$=1.8754, P=0.12, sex $F_{1,40}$=0.3026, P=0.585, interaction $F_{5,40}$=1.319, P=0.2756). **(D)** Mean REM event duration in male and female mice injected with monomeric αSyn or αSyn PFF in the stomach (group $F_{5,40}$=3.499, P=0.010223, sex $F_{1,40}$=2.14, P=0.1506, interaction $F_{5,40}$=0.507, P=0.769). (**E**) Percentage of time NREM in male and female mice injected with monomeric αSyn or αSyn PFF in the stomach (group $F_{5,40}$=1.845, P=0.126, sex $F_{1,40}$=1.474, P=0.2317, interaction $F_{5,40}$=0.416, P=0.834819). **(F)** Mean NREM event duration in male and female mice injected with monomeric αSyn or αSyn PFF in the stomach (group $F_{5,40}$=1.55, P=0.194, sex $F_{1,40}$=2.37, P=0.1314, interaction $F_{5,40}$=1.619, P=0.1769). (**G**) Number of RBD event in male and female mice injected with monomeric αSyn or αSyn PFF in the stomach (group $F_{5,40}$=11.80, P=0.0000001, sex $F_{1,40}$=0.38, P=0.5395, interaction $F_{5,40}$=0.836, P=0.532). (**H**) Percentage of RBD events over REM events in male and female mice injected with monomeric αSyn or αSyn PFF in the stomach. (group $F_{5,40}$=11.62, P=0.000001, sex $F_{1,40}$=0.377, P=0.5424, interaction $F_{5,40}$=1.22, P=0.317). (**I**) Percentage of time in corner paired with TMT smell in male and female mice injected with monomeric αSyn or αSyn PFF in the stomach (group $F_{5,59}$=2.456, P=0.043, sex $F_{1,59}$=0.2081, P=0.6499, interaction $F_{5,59}$=0.8416, P=0.5256). (**J**) Percentage of time in corner paired with chocolate smell in male and female mice injected with monomeric αSyn or αSyn PFF in the stomach (group $F_{5,59}$=1.65, P=0.1587, sex $F_{1,59}$=1.11, P=0.2951, interaction $F_{5,59}$=0.57, P=0.722). (**K**) Olfactory discrimination index in male and female mice injected with monomeric αSyn or αSyn PFF in the stomach (group $F_{5,59}=3.075$, $P=0.01558$, sex $F_{1,59}=0.68$, $P=0.411$, interaction $F_{5,59}=0.478$, $P=0.7907$). (**L**) Sucrose preference index in male and female mice injected with monomeric αSyn or αSyn PFF in the stomach (group $F_{5,59}=3.958$, $P=0.003667$, sex $F_{1,59}=0.03$, $P=0.863$, interaction $F_{5,59}=1.208$, $P=0.3166$). (**M**) Time in the light zone during the dark-light test in male and female mice injected with monomeric αSyn or αSyn PFF in the stomach (group $F_{5,59}=6.147$, $P=0.00012$, sex $F_{1,59}=3.57$, $P=0.0635$, interaction $F_{5,59}=0.145$, $P=0.9806$). Data are expressed as mean±SEM. All individual point represents individual animals. * $P<0.05$, ** $P<0.01$, *** $P<0.001$, **** $P<0.0001$. Statistical data represent 2-way ANOVA sex x group interaction.

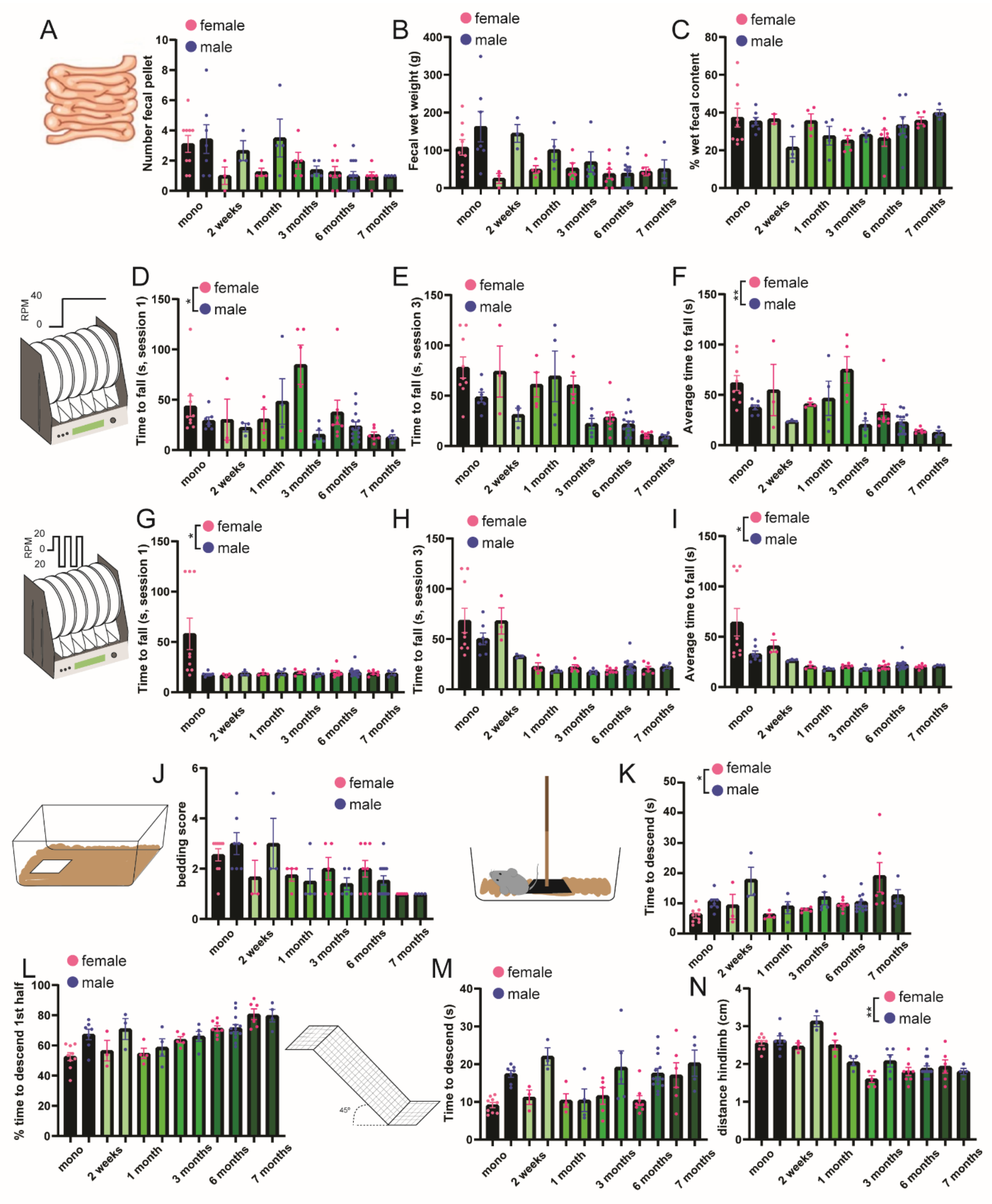


**Fig. S19. Related to Figure 4-5.**

(**A**) Number of fecal pellets in male and female mice injected with monomeric αSyn or αSyn PFF in the stomach (group $F_{5,59}$=5.33, P=0.00041, sex $F_{1,59}$=2.43, P=0.123, interaction $F_{5,59}$=1.46, P=0.214). **(B)** Dry fecal weight in male and female mice injected with monomeric αSyn or αSyn PFF in the stomach (group $F_{5,59}$=5.10, P=0.000589, sex $F_{1,59}$=9.3, P=0.00341, interaction $F_{5,59}$=1.72, P=0.144). (**C**) Percentage of water content in fecal pellets in male and female mice injected with monomeric αSyn or αSyn PFF in the stomach (group $F_{5,50}$=2.10, P=0.08, sex $F_{1,50}$=0.47, P=0.493, interaction $F_{5,50}$=1.38, P=0.246). (**D**) Latency to fall in constant speed 40 RPM rotarod schedule session 1 in male and female mice injected with monomeric αSyn or αSyn PFF in the stomach (group $F_{5,59}$=2.33, P=0.0534, sex $F_{1,59}$=5.21, P=0.026, interaction $F_{5,59}$=3.15, P=0.0136). (**E**) Latency to fall in constant speed 40 RPM rotarod schedule session 3 in male and female mice injected with monomeric αSyn or αSyn PFF in the stomach (group $F_{5,59}$=11.53, P=0.0000001, sex $F_{1,59}$=10.74, P=0.00175, interaction $F_{5,59}$=2.16, P=0.07). (**F**) Average latency to fall across all session of constant speed 40 RPM rotarod schedule in male and female mice injected with monomeric αSyn or αSyn PFF in the stomach (group $F_{5,59}$=6.59, P=0.000061, sex $F_{1,59}$=16.568, P=0.000142, interaction $F_{5,59}$=3.47, P=0.00804). (**G**) Latency to fall in docking speed 20 RPM rotarod schedule session 1 in male and female mice injected with monomeric αSyn or αSyn PFF in the stomach (group $F_{5,59}$=2.76, P=0.0261, sex $F_{1,59}$=2.08, P=0.153, interaction $F_{5,59}$=3.18, P=0.0129). (**H**) Latency to fall in docking speed 20 RPM rotarod schedule session 3 in male and female mice injected with monomeric αSyn or αSyn PFF in the stomach (group $F_{5,59}$=16.37, P=0.0000001, sex $F_{1,59}$=5.22, P=0.0258, interaction $F_{5,59}$=2.101, P=0.077). (**I**) Average latency to fall across all session of docking speed 20 RPM rotarod schedule in male and female mice injected with monomeric αSyn or αSyn PFF in the stomach (group $F_{5,59}$=7.71, P=0.000012, sex $F_{1,59}$=3.64, P=0.061, interaction $F_{5,59}$=2.48, P=0.0416). (**J**) Bedding score in male and female mice injected with monomeric αSyn or αSyn PFF in the stomach (group $F_{5,59}$=6.42, P=0.000079, sex $F_{1,59}$=0.12, P=0.72, interaction $F_{5,59}$=1.58, P=0.179). (**K**) Time to descend in the Pole test in male and female mice injected with monomeric αSyn or αSyn PFF in the stomach (group $F_{5,59}$=3.98, P=0.0035, sex $F_{1,59}$=3.17, P=0.08, interaction $F_{5,59}$=2.37, P=0.0497). (**L**) Percentage of time to descend the first half of the pole test compared to the entire duration in male and female mice injected with monomeric αSyn or αSyn PFF in the stomach (group $F_{5,59}$=11.23, P=0.0000001, sex $F_{1,59}$=7.198, P=0.009456, interaction $F_{5,59}$=2.108, P=0.077). (**M**) Time to descend in the inclined platform test in male and female mice injected with monomeric αSyn or αSyn PFF in the stomach (group $F_{5,59}$=2.77, P=0.025, sex $F_{1,59}$=21.33, P=0.000021, interaction $F_{5,59}$=1.17, P=0.33). (**N**) Average distance of hindlimb during the inclined platform test in male and female mice injected with monomeric αSyn or αSyn PFF in the stomach (group $F_{5,59}$=24.825, P=0.0000001, sex $F_{1,59}$=2.75, P=0.102, interaction $F_{5,59}$=4.185, P=0.0025). Data are expressed as mean±SEM. All individual points represent individual animals. * P<0.05, ** P<0.01. Statistical data represent 2-way ANOVA sex x group interaction.

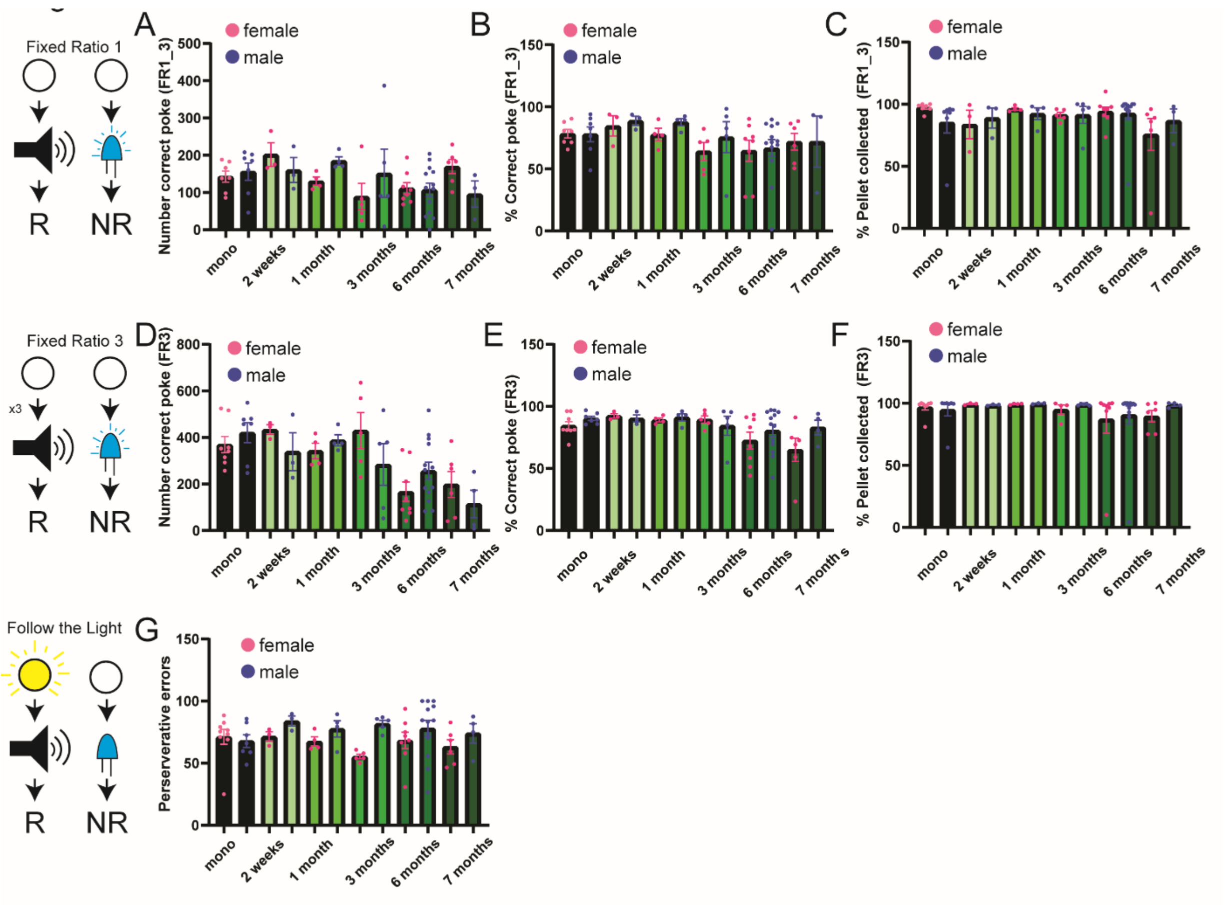


**Fig. S20. Related to Figure 5.**

**(A)** Number of correct pokes in the third session of the Fixed Ratio 1 schedule in male and female mice injected with monomeric αSyn or αSyn PFF in the stomach (group $F_{5,56}$=1.65, P=0.16, sex $F_{1,56}$=0.0099, P=0.921, interaction $F_{5,56}$=1.39, P=0.24). (**B**) Percentage of correct poke in the third session of the Fixed Ratio 1 schedule in male and female mice injected with monomeric αSyn or αSyn PFF in the stomach (group $F_{5,56}$=1.69, P=0.151, sex $F_{1,56}$=0.73, P=0.395, interaction $F_{5,56}$=0.165, P=0.974). (**C**) Percentage of pellet collected within 10s in the third session of the Fixed Ratio 1 schedule in male and female mice injected with monomeric αSyn or αSyn PFF in the stomach (group $F_{5,56}$=0.786, P=0.563, sex $F_{1,56}$=0.002, P=0.967, interaction $F_{5,56}$=0.56, P=0.729). (**D**) Number of correct pokes in the Fixed Ratio 3 schedule in male and female mice injected with monomeric αSyn or αSyn PFF in the stomach (group $F_{5,58}$=7.37, P=0.000021, sex $F_{1,58}$=0.4798, P=0.491, interaction $F_{5,58}$=1.69, P=0.149). (**E**) Percentage of correct poke in the Fixed Ratio 3 schedule in male and female mice injected with monomeric αSyn or αSyn PFF in the stomach (group $F_{5,58}$=2.896, P=0.021, sex $F_{1,58}$=1.493, P=0.226, interaction $F_{5,58}$=0.801, P=0.553). (**F**) Percentage of pellet collected within 10s in the Fixed Ratio 3 schedule in male and female mice injected with monomeric αSyn or αSyn PFF in the stomach (group $F_{5,56}$=0.659, P=0.656, sex $F_{1,56}$=0.262, P=0.6109, interaction $F_{5,56}$=0.141, P=0.9817). (**G**) Number of perseverative errors in the "Follow the light" schedule in male and female mice injected with monomeric αSyn or αSyn PFF in the stomach (group $F_{5,59}$=0.405, P=0.8433, sex $F_{1,59}$=6.826, P=0.0113, interaction $F_{5,59}$=1.065, P=0.388). Data are expressed as mean±SEM. All individual point represents individual animals. Statistical data represent 2-way ANOVA sex x group interaction.

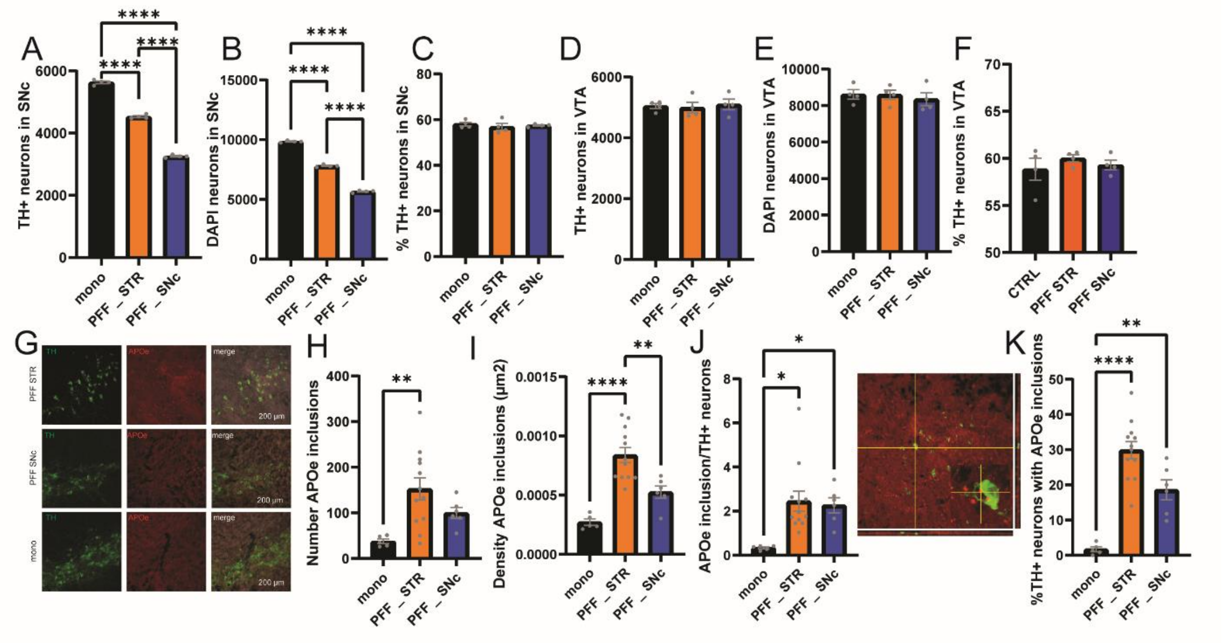


**Fig. S21. Related to Figure 7.**

(**A**) Number of TH-positive neurons in the SNc in WT mice injected with monomeric αSyn or αSyn PFF in the Striatum (orange) or the SNc (blue) $F_{2,9}$=1385, P<0.0001. (**B**) Number of DAPI-positive neurons in the SNc in WT mice injected with monomeric αSyn or αSyn PFF in the Striatum (orange) or the SNc (blue) $F_{2,9}$=1816, P<0.0001. (**C**) Percentage of DAPI neurons expressing TH in the SNc in WT mice injected with monomeric αSyn or αSyn PFF in the Striatum (orange) or the SNc (blue) $F_{2,9}$=0.3376, P=0.7221. **(D)** Number of TH-positive neurons in the VTA in WT mice injected with monomeric αSyn or αSyn PFF in the Striatum (orange) or the SNc (blue) $F_{2,9}$=0.1122, P=0.8951. (**E**) Number of DAPI-positive neurons in the VTA in WT mice injected with monomeric αSyn or αSyn PFF in the Striatum (orange) or the SNc (blue) $F_{2,9}$=0.2433, P=0.789. (**F**) Percentage of DAPI neurons expressing TH in the VTA in WT mice injected with monomeric or αSyn PFF in the Striatum (orange) or the SNc (blue) $F_{2,9}$=0.5842, P=0.5774. (**G**) Confocal images showing the presence of ApoE inclusions in the vicinity of TH-positive neurons located in the SNc, following injection of monomeric αSyn or αSyn PFF in the SNc or Striatum. (**H**) Total number of ApoE inclusion in the SNc of mice injected with monomeric αSyn or αSyn PFF in the Striatum (orange) or the SNc (blue) $F_{2,20}$=5.806, P=0.0103. (**I**) Density of APOe inclusion in the SNc of mice injected with monomeric αSyn or αSyn PFF in the Striatum (orange) or the SNc (blue) $F_{2,20}$=19.61, P<0.0001. (**J**) Number of APOe inclusion over the number of TH-positive neurons in the SNC of mice injected with monomeric αSyn or αSyn PFF in the Striatum (orange) or the SNc (blue) $F_{2,20}$=5.529, P=0.0123. (**K**) Confocal images of APOe inclusion and TH-positive neurons, with orthogonal projection showing that the inclusion is located within the neuron. Quantification of TH-positive neurons located in the SNc that contain at least one inclusion within its soma $F_{2,20}$=27.50, P<0.0001. Data are expressed as mean±SEM. All individual point represents individual animals. * $P<0.05$, ** $P<0.01$, **** $P<0.0001$. Statistical data represent post hoc analyses compared to monomeric αSyn group following 1-way ANOVA.

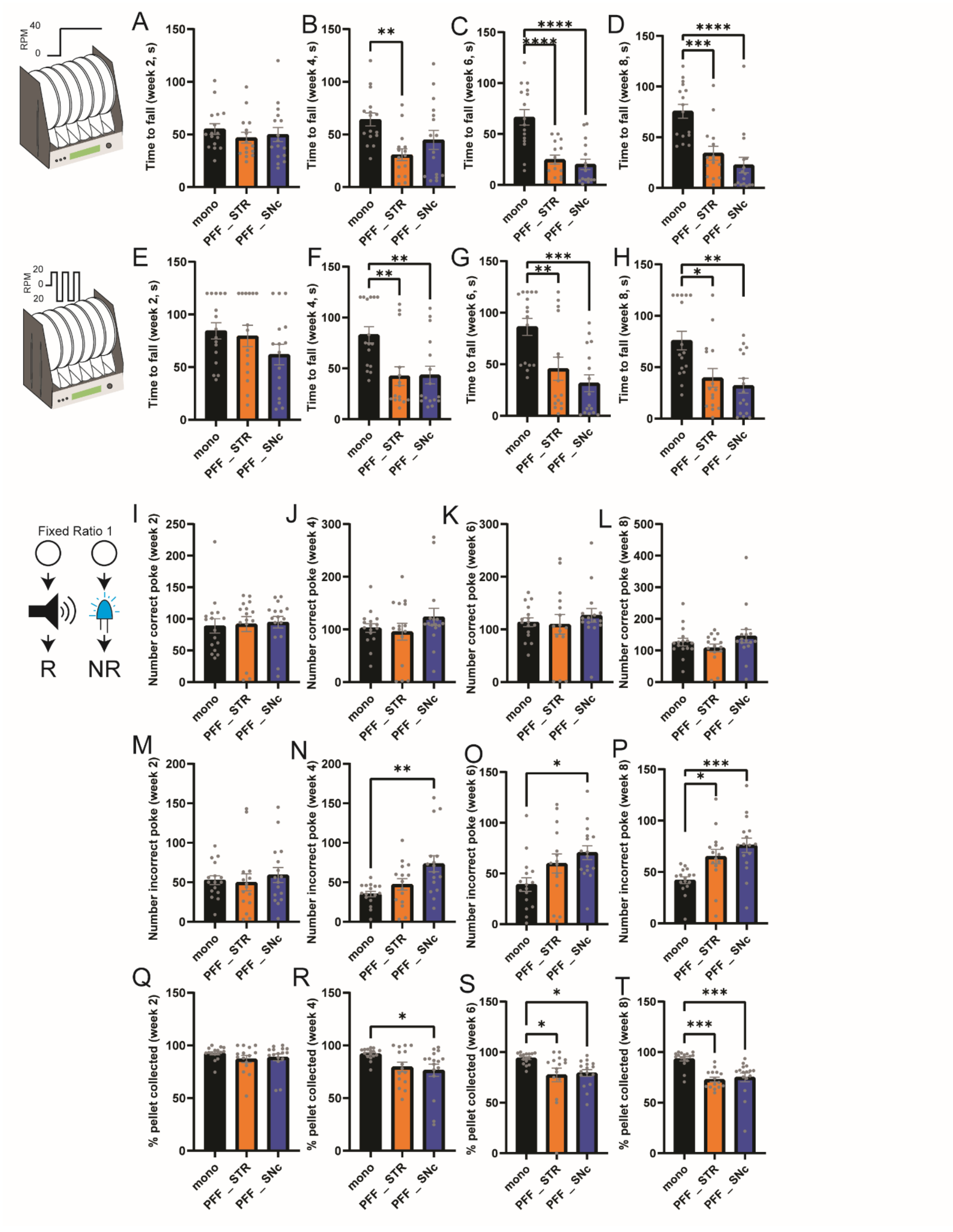


**Fig. S22. Related to Figure 7.**

Latency to fall in the constant speed 40 RPM rotatord schedule following 2 (**A**, $F_{2,44}$=0.5268, P=0.5942), 4 (**B**, $F_{2,44}$=5.492, P=0.0074), 6 (**C**, $F_{2,44}$=18.53, P<0.0001) or 8 (**D**, $F_{2,44}$=15.46, P<0.0001) weeks post injection of monomeric αSyn or αSyn PFF in the Striatum (orange) or the SNc (blue). Latency to fall in the docking speed 20 RPM rotatord schedule following 2 (**E** $F_{2,44}$=1.680, P=0.1982), 4 (**F**, $F_{2,44}$=7.390, P=0.0017), 6 (**G**, $F_{2,44}$=9.548, P=0.0004) or 8 (**H**, $F_{2,44}$=7.670, P=0.0014) weeks post injection of monomeric αSyn or αSyn PFF in the Striatum (orange) or the SNc (blue). Number of correct poke in the Fixed Ratio 1 following 2 (**I**, $F_{2,44}$=0.08087, P=0.9224), 4 (**J**, $F_{2,44}$=1.106, P=0.3399), 6 (**K**, $F_{2,44}$=0.4296, P=0.6535) or 8 (**L**, $F_{2,44}$=1.371, P=0.2645) weeks post injection of monomeric αSyn or αSyn PFF in the Striatum (orange) or the SNc (blue). Number of incorrect poke in the Fixed Ratio 1 following 2 (**M**, $F_{2,44}$=0.2924, P=0.7479), 4 (**N**, $F_{2,44}$=6.860, P=0.0026), 6 (**O**, $F_{2,44}$=4.488, P=0.0168) or 8 (**P**, $F_{2,44}$=8.689, P=0.0007) weeks post injection of monomeric αSyn or αSyn PFF in the Striatum (orange) or the SNc (blue). Number of pellets collected within 10s in the Fixed Ratio 1 following 2 (**Q**, $F_{2,44}$=0.1450, P=0.8654), 4 (**R**, $F_{2,44}$=2.772, P=0.0735), 6 (**S**, $F_{2,44}$=1.177, P=0.3177) or 8 (**T**, $F_{2,44}$=4.687, P=0.0143) weeks post injection of monomeric αSyn or αSyn PFF in the Striatum (orange) or the SNc (blue). Data are expressed as mean±SEM. All individual point represents individual animals. * P<0.05, ** P<0.01, *** P<0.001, **** P<0.0001. Statistical data represent post hoc analyses compared to monomeric αSyn group following 1-way ANOVA.

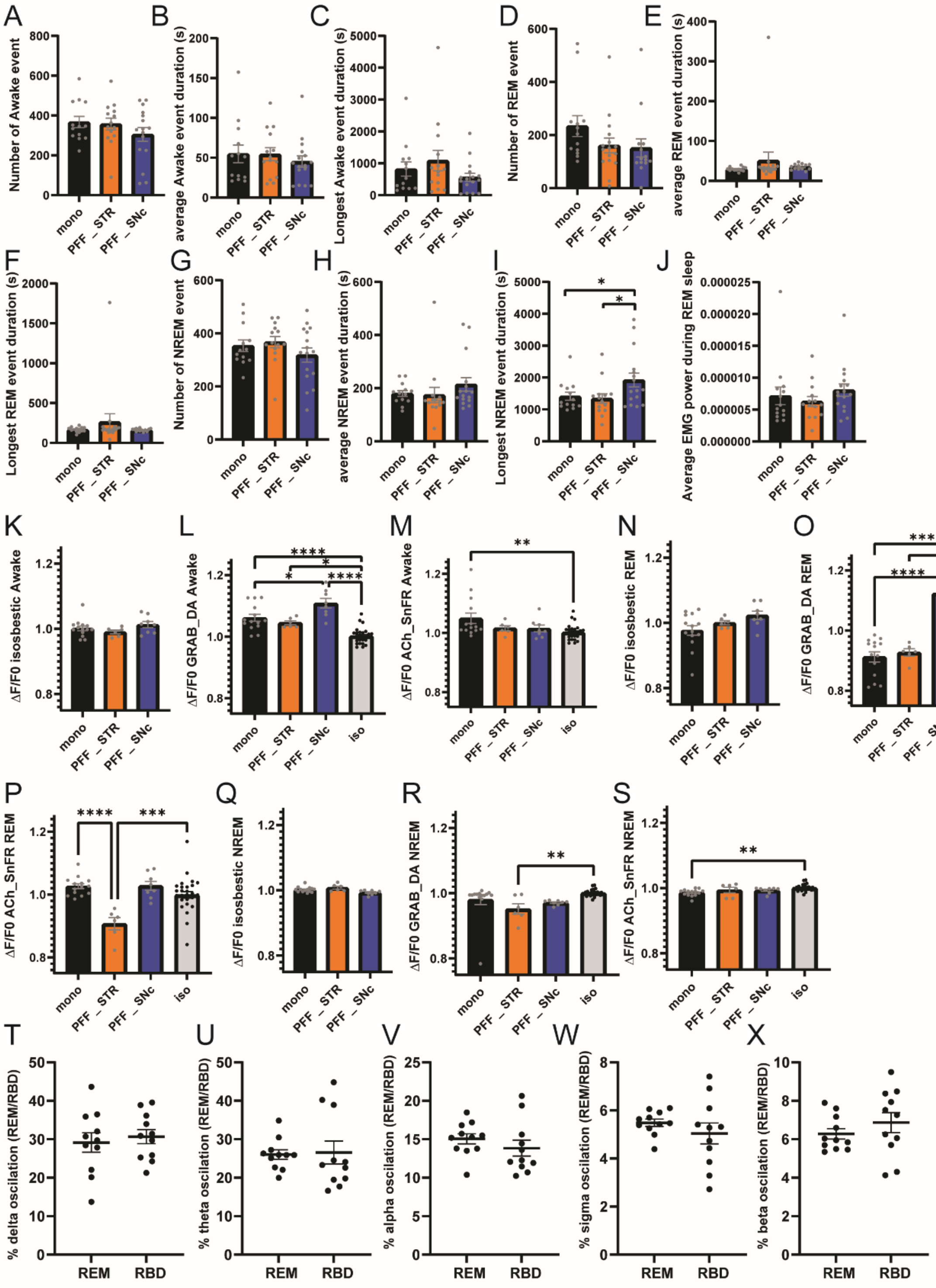


**Fig. S23. Related to Figure 7.**

(**A**) Number of awake events during the 24h recording in mice injected with monomeric αSyn or αSyn PFF in the Striatum (orange) or the SNc (blue) ($F_{2,40}$=1.171, P=0.3204). (**B**) Average awake event duration during the 24h recording in mice injected with monomeric αSyn or αSyn PFF in the Striatum (orange) or the SNc (blue) ($F_{2,40}$=0.4081, P=0.6677). (**C**) Longest awake event duration during the 24h recording in mice injected with monomeric αSyn or αSyn PFF in the Striatum (orange) or the SNc (blue) ($F_{2,40}$=1.374, P=0.2648). (**D**) Number of REM events during the 24h recording in mice injected with monomeric αSyn or αSyn PFF in the Striatum (orange) or the SNc (blue) ($F_{2,40}$=1.737, P=0.1890). (**E**) Average REM event duration during the 24h recording in mice injected with monomeric αSyn or αSyn PFF in the Striatum (orange) or the SNc (blue) ($F_{2,39}$=0.7329, P=0.4870). (**F**) Longest REM event duration during the 24h recording in mice injected with monomeric αSyn or αSyn PFF in the Striatum (orange) or the SNc (blue) ($F_{2,39}$=0.8026, P=0.4554). (**G**) Number of NREM event during the 24h recording in mice injected with monomeric αSyn or αSyn PFF in the Striatum (orange) or the SNc (blue) ($F_{2,40}$=1.200, P=0.3119). (**H**) Average NREM event duration during the 24h recording in mice injected with monomeric αSyn or αSyn PFF in the Striatum (orange) or the SNc (blue) ($F_{2,40}$=0.8806, P=0.4224). (**I**) Longest NREM event duration during the 24h recording in mice injected with monomeric αSyn or αSyn PFF in the Striatum (orange) or the SNc (blue) ($F_{2,40}$=3.415, P=0.0427, here post hoc test is LSD). (**J**) Average EMG signal during REM event during the 24h recording in mice injected with monomeric αSyn or αSyn PFF in the Striatum (orange) or the SNc (blue) ($F_{2,41}$=0.7103, P=0.4975). (**K**) Fiber photometry signal ΔF/F0 in the DMS of the isosbestic signal during awake events in mice injected with monomeric αSyn or αSyn PFF in the Striatum (orange) or the SNc (blue) ($F_{2,25}$=1.453, P=0.2530). (**L**) Fiber photometry signal ΔF/F0 in the DMS of the rGRAB-DA signal during awake events in mice injected with monomeric αSyn or αSyn PFF in the Striatum (orange) or the SNc (blue). Grey histogram represent the average isosbestic signal during awake events of all animals (Mono αSyn and αSyn PFF) ($F_{3,52}$=26.04, P<0.0001). (**M**) Fiber photometry signal ΔF/F0 in the DMS of the Ach-SnFr signal during awake events in mice injected with monomeric αSyn or αSyn PFF in the Striatum (orange) or the SNc (blue). Grey histogram represent the average isosbestic signal during awake events of all animals (Mono and αSyn PFF) ($F_{3,52}$=4.381, P=0.0080). (**N**) Fiber photometry signal ΔF/F0 in the DMS of the isosbestic signal during REM events in mice injected with monomeric αSyn or αSyn PFF in the Striatum (orange) or the SNc (blue) ($F_{2,25}$=2.865, P=0.0758). (**O**) Fiber photometry signal ΔF/F0 in the DMS of the rGRAB -DA signal during REM events in mice injected with monomeric αSyn or αSyn PFF in the Striatum (orange) or the SNc (blue). Grey histogram represent the average isosbestic signal during REM events of all animals (Mono and αSyn PFF) ($F_{3,52}$=28.34, P<0.00001). (**P**) Fiber photometry signal ΔF/F0 in the DMS of the Ach-SnFr signal during REM events in mice injected with monomeric αSyn or αSyn PFF in the Striatum (orange) or the SNc (blue). Grey histogram represent the average isosbestic signal during REM events of all animals (Mono and αSyn PFF) ($F_{3,52}$=9.936, P<0.0001). (**Q**) Fiber photometry signal ΔF/F0 in the DMS of the isosbestic signal during NREM events in mice injected with monomeric αSyn or αSyn PFF in the Striatum (orange) or the SNc (blue) ($F_{2,25}$=5.776, P=0.0087). (**R**) Fiber photometry signal ΔF/F0 in the DMS of the rGRAB-DA signal during NREM events in mice injected with monomeric αSyn or αSyn PFF in the Striatum (orange) or the SNc (blue). Grey histogram represent the average isosbestic signal during NREM events of all animals (Mono and αSyn PFF) ($F_{3,52}$=5.045, P=0.0038). (**S**) Fiber photometry signal ΔF/F0 in the DMS of the Ach-SnFr signal during NREM events in mice injected with monomeric αSyn or αSyn PFF in the Striatum (orange) or the SNc (blue). Grey histogram represent the average isosbestic signal during NREM events of all animals (Mono and αSyn PFF) ($F_{3,52}$=5.358,

P=0.0027). Comparison delta (**T**, $T_{20}$=0.4848, P=0.6332), theta (**U**, $T_{20}$=0.1594, P=0.8750), alpha (**V**, $T_{20}$=0.9830, P=0.3373), sigma (**W**, $T_{20}$=0.9656, P=0.3458) and beta (**X**, $T_{20}$=0.9954, P=0.3314) periodogram oscillation expressed as a percentage of all other oscillation in the frontal cortex between REM and RBD events in mice injected with αSyn PFF in the SNc. Data are expressed as mean±SEM. All individual point represents individual animals. * $P<0.05$, ** $P<0.01$, *** $P<0.001$, **** $P<0.0001$. Statistical data represent post hoc analyses compared to the monomeric group following 1-way ANOVA. Periodogram oscillation are compared using paired t-test.
